# Generating Synthetic Task-based Brain Fingerprints for Population Neuroscience Using Deep Learning

**DOI:** 10.1101/2024.08.03.606469

**Authors:** Emin Serin, Kerstin Ritter, Gunter Schumann, Tobias Banaschewski, Andre Marquand, Henrik Walter, environMENTAL consortium

## Abstract

Task-based functional magnetic resonance imaging (tb-fMRI) reveals individual differences in the neural basis of cognitive functions by linking specific tasks to neural responses. However, scaling tb-fMRI to population-level studies is challenging due to its cognitive demands, variations in task design across studies, and the limited scope of tasks in large datasets. To address this, we propose DeepTaskGen, a deep-learning approach that generates non-acquired task-based contrast maps from resting-state fMRI (rs-fMRI) data. Our approach enables generating synthetic task images for non-acquired tasks within the study protocol. We validate this approach using the Human Connectome Project lifespan data, then generate 47 contrast maps from 7 different cognitive tasks for over 20,000 individuals from UK Biobank. DeepTaskGen outperforms several benchmarks in generating synthetic task-contrast maps, exhibiting superior reconstruction performance while retaining inter-individual variation essential for biomarker development. Notably, we further showed that synthetic task contrast maps achieved similar or greater performance compared to actual task contrast maps and resting-state connectomes for predicting a wide range of demographic, cognitive, and clinical variables. This approach will facilitate the study of individual differences and the generation of task-related biomarkers by enabling the generation of arbitrary functional cognitive tasks from readily available rs-fMRI data.

## Introduction

Characterizing inter-individual variability in brain activity patterns has become crucial to understanding individual differences in cognition, behavior, and the risk for or resilience to mental disorders. Over the years, many techniques have been developed to discover the neural basis of individual differences. Among these, task-based functional magnetic resonance imaging (tb-fMRI) has emerged as a powerful technique as it can non-invasively measure changes in brain activity associated with a specific cognitive task. For example, the emotional face-matching task has been shown to activate several brain regions, particularly the amygdala^1^, which has associations with mental disorders such as depression and anxiety^2^. Researchers can potentially identify individuals at risk of mental disorders by examining individual differences in task-related brain activation. Moreover, an increasing number of studies highlight that task-related brain activity improves prediction performance for individual differences in cognitive and behavioral traits, including fluid intelligence, cognitive flexibility, working memory capacity, and general cognitive ability, compared to task-free (resting-state) brain activity^3–7^. The flexibility of tb-fMRI in eliciting task-specific activations, coupled with its improved predictive power and promising findings regarding reliability^8,9^, has established its role as a robust tool for understanding individual differences in brain activity^10^.

Despite its advantages, task-based brain imaging has several limitations. Firstly, it is less scalable than other brain imaging techniques like anatomical imaging or resting-state fMRI (rs-fMRI) due to its cognitive demands, the need for subject training and compliance, and because different studies often use different task variants, which complicates aggregation across studies. Therefore, large consortium studies such as the Human Connectome Project^11^, IMAGEN^12^, and ABCD^13^ mostly have data from only a few (usually simple) tasks, with some notable exceptions, such as the Human Connectome Project (HCP), which provides a comprehensive task battery^11^. This is even more pronounced in populations that have difficulty in performing experimental tasks, such as children or individuals with neurological or psychiatric conditions. The second limitation arises from variations in experimental task designs. Reliable and advanced biomarker discovery requires large-scale, multi-site datasets to represent individual variations on a population scale. However, this introduces the challenge of standardization, as researchers must control for differences in scanning sites, parameters, and variations in task designs. In many instances it is simply not possible to aggregate task data from different cohorts because the differences between paradigms used in each cohort are too large. This challenge is especially pressing in big-data studies that require large samples to provide sufficient power to detect subtle effects or to characterize inter-individual variability.

In this paper, we address these limitations by generating *synthetic* task-based brain images from task-free resting-state data. This approach builds upon the discovery that resting-state and task-based brain activity share the same intrinsic network architecture.^14–16^ Preliminary work has aimed to predict inter-individual variations in task-induced brain activity from resting-state data^15,17^ Although interesting, these approaches have important limitations in that they either fail to fully capture the complexity necessary for retaining individual-specific information, lack effective methods for transferring learned parameters to new datasets or do not cover the entire brain. Most importantly, all such approaches have only been applied to cases where the true task data were acquired. It is much more challenging to generate contrast images in cases where the tasks were not acquired at all. We also argue that this feature is essential for synthetic task images to be useful for biomarker construction. To demonstrate usefulness, it is of utmost importance to show the predictive utility of synthetic task images for clinically relevant outcomes, which is the aim of this study.

Building upon prior work^17^, we present a volumetric neural network architecture, DeepTaskGen, that (i) addresses all the limitations mentioned above, (ii) provides the ability to generate synthetic task-based images in cohorts where these tasks were not acquired, and (iii) provides superior reconstruction performance in several datasets compared to different baseline methods, while simultaneously retaining individual differences that are essential for biomarker development. To do so, we first train DeepTaskGen on the Human Connectome Project Young-Adult (HCP-YA)^11^ and demonstrate competitive performance in retaining individual differences while predicting task-based contrast maps (i.e., task-based brain activity). Next, we apply our trained network to the Human Connectome Project Development (HCP-D) dataset^18^ to show its generalizability by predicting task-based contrast maps absent from this dataset. Next, we apply our trained network to generate synthetic task images for a comprehensive battery of cognitive functions for one of the largest population-based datasets currently available (UK Biobank).^19^ Finally, we demonstrate the utility of these synthetic data as biomarkers by predicting subjects’ age, sex, fluid intelligence, grip strength, and overall health, as well as several clinical measures, including hypertension and depression diagnosis, alcohol use frequency, anxiety, and depressive symptom scores. Remarkedly, we show that in many cases, synthetic images achieve similar or greater performance than actual images.

## Results

### DeepTaskGen demonstrates overall better reconstruction performance and discriminability on the training dataset

First, we trained DeepTaskGen on the HCP-YA dataset using a train set (𝑛 = *827*) and a validation set (𝑛 = *92*). We used performance metrics assessing two distinct characteristics of synthetic contrast maps. The reconstruction performance, measured by similarity metrics such as Pearson’s correlation or Dice Coefficient, quantize overlapping or association between predicted and actual maps. These metrics are widely adopted to evaluate generative methods in the literature^15,17,20–22^. Discriminability (or subject identification) metrics evaluate the subject-specific variation among predicted images, which is essential for biomarker discovery. To evaluate it, we used the diagonality index, a metric adopted in previous studies^17,20,22^. In this study, we assessed DeepTaskGen’s reconstruction performance using Pearson’s correlation, and diagonality index scores on a separate test set (𝑛 = *39*). Additionally, while not specific to this context, fingerprinting scores are widely accepted for assessing subject identifiability from brain images^23^. Therefore, we present fingerprinting scores along with the Dice AUC score in the Supplementary Material (Supplementary Figure 2, and Table 4,6). Next, we used permutation testing to compare these performance measures with group-average task contrasts, a linear model^15^, and retest scans (which can be seen as a theoretical upper bound on prediction ability). Figure 1c displays the results for seven representative main task contrasts of seven different paradigms, while Supplementary Figure 1 and Supplementary Tables 3,4 provide the performance for all 47 task contrasts and their test statistics, respectively. Additionally, we present the actual and predicted group-average maps for the seven representative main task contrasts in Figure 2a.

**Fig. 1:**
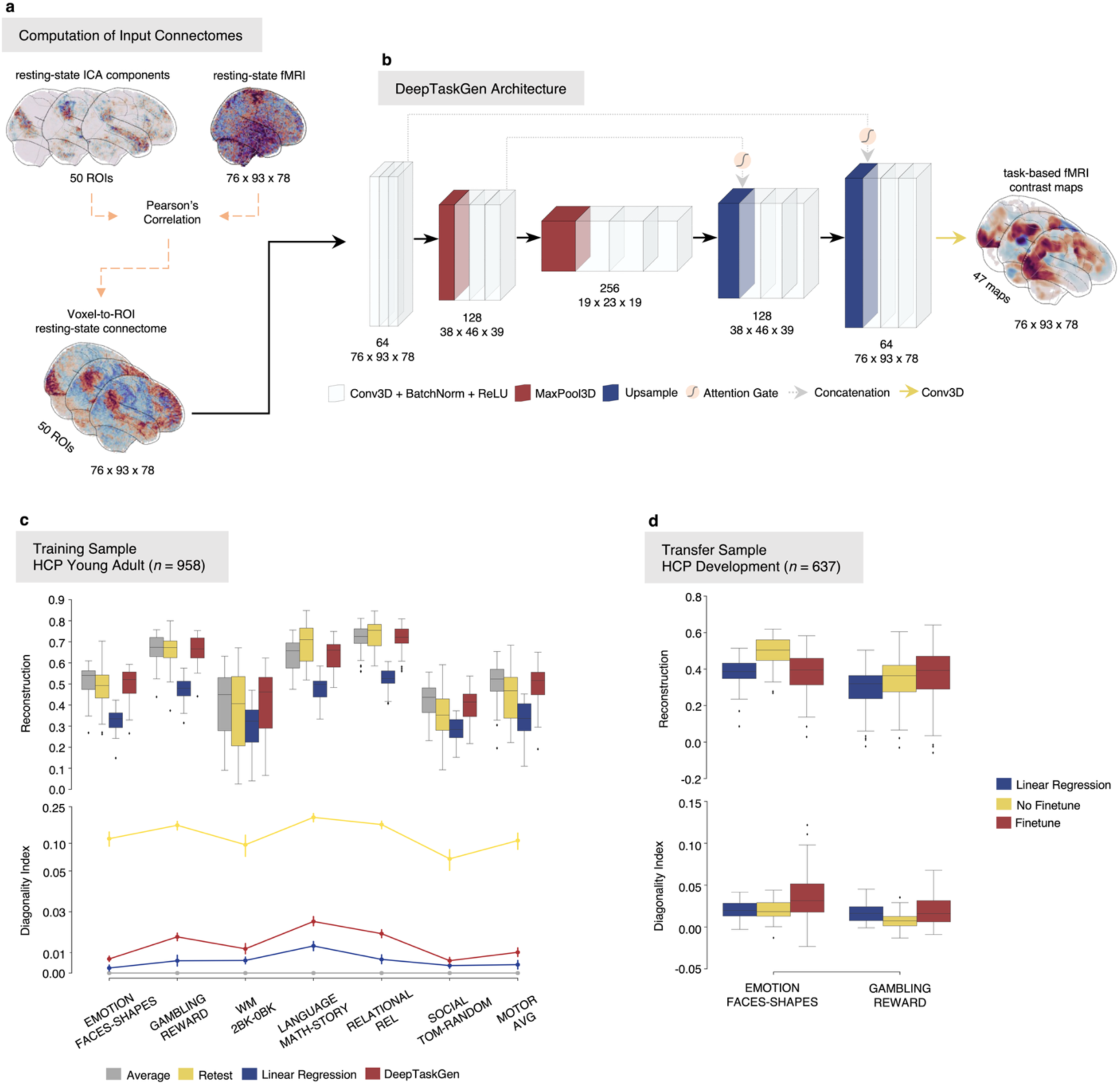
Input, DeepTaskGen architecture, and model evaluation on training and independent samples. **a.** We utilized voxel-to-ROI rs-fMRI connectomes as input. A connectome was constructed for each subject by calculating the full correlation between the averaged time series from 50 ICA-based ROIs and the time series of individual voxels. **b.** Task-contrast maps for various tasks were predicted from rs-fMRI connectomes using our proposed DeepTaskGen architecture. DeepTaskGen is a volumetric U-Net model with attention mechanism that processes the input resting-state connectome through a series of convolutional blocks, each comprising a 3D convolution layer, batch normalization, and a non-linear activation function. By utilizing max pooling, the model compresses images while preserving task-relevant patterns, and then, it up-samples the images to align with the output task contrast maps. The numbers below each block represent the output shape of each block and the number of feature maps (above). The details of the architecture are presented in Supplementary Table 1. **c.** We trained and evaluated DeepTaskGen on the HCP Young Adult dataset. The figure above shows the reconstruction performance computed by taking Pearson’s correlation between predicted and actual contrast maps for representative contrasts from seven distinct tasks. The figure below displays the diagonality index (the difference between the on-diagonal and the mean off-diagonal elements in a correlation matrix, normalized by the mean on-diagonal values), evaluating models’ discriminability performance. We compared DeepTaskGen with methods like group-averaged contrast maps, retest scans, and a linear model (each depicted in distinct colors). **d.** We further fine-tuned the trained DeepTaskGen model on the HCP Development dataset using either task contrasts (e.g., GAMBLING REWARD). We predicted the other contrast (e.g., EMOTION FACES-SHAPES). The fine-tuned model was compared to the non-fine-tuned DeepTaskGen and linear models (shown in distinct colors). Reconstruction performance and discriminability were again used to assess the models’ performance for each task contrast.

**Fig. 2:**
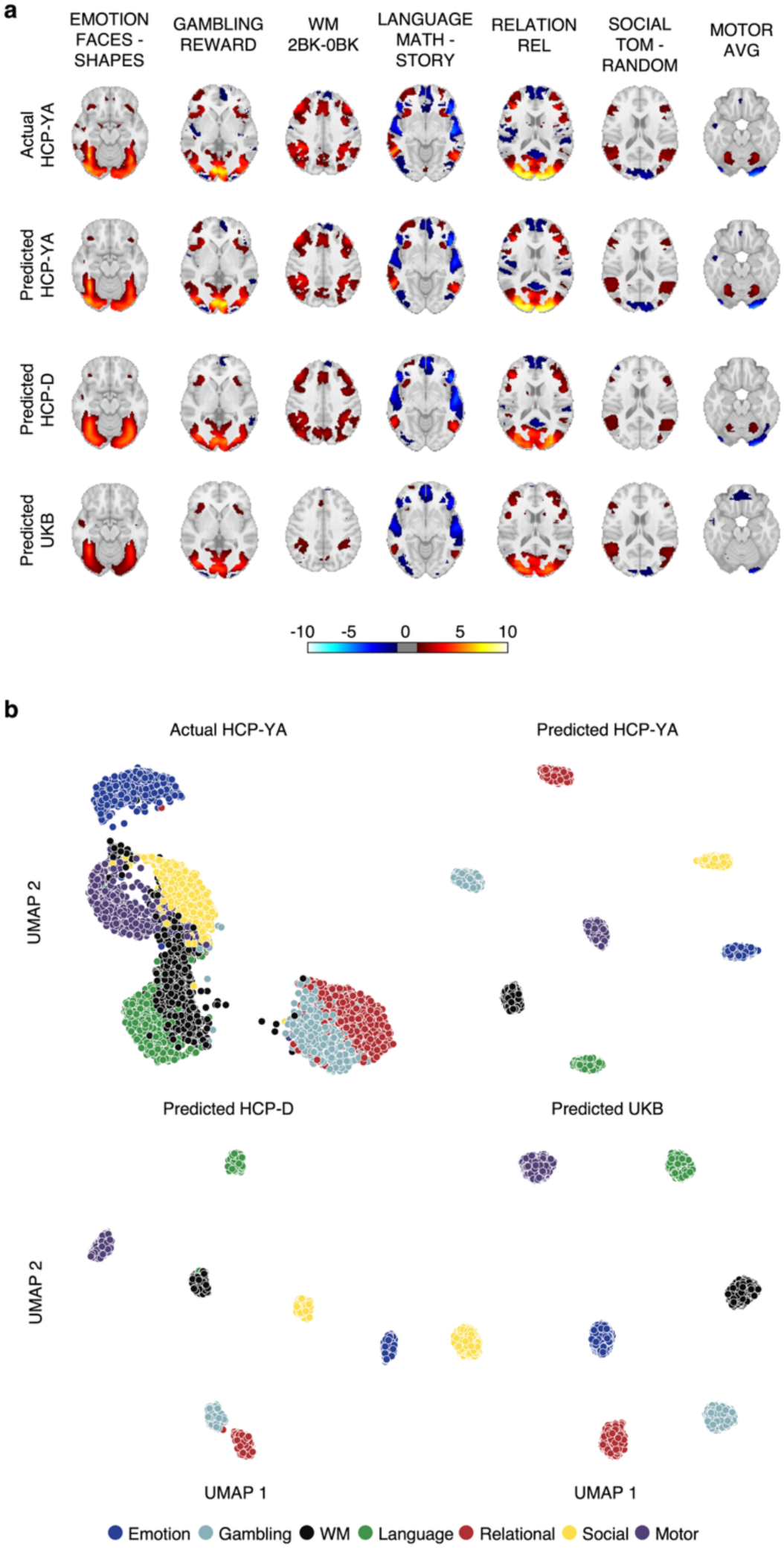
Visualization of group-level contrast maps and similarity (or dissimilarity) among subjects. **a.** Group-level contrast maps for seven representative tasks were generated for predicted maps on three datasets (HCP Young Adult, HCP Development, and UK Biobank) and compared to actual task contrast maps from HCP-YA. Activations and deactivations are displayed using scaled z-scores, with colors indicating the magnitude of the effect (shown in the color gradient bar). **b.** To visualize within- and between-task variance among subjects, we employed Uniform Manifold Approximation and Projection (UMAP), a non-linear dimensionality reduction technique. UMAP projects high-dimensional task contrast maps into two dimensions for clear visualization, allowing us to assess if the projected datasets contain individual- and task-specific information necessary for downstream analyses. Each dot represents a subject’s task contrast map for the seven tasks, colored accordingly. Similar subjects are positioned closer together, indicating similarity, while distant dots represent dissimilarity. Accordingly, wider spread within tasks and greater distance between tasks represent higher within- and between-task variability. Note that we fitted a separate UMAP model to each dataset, as indicated in each column and row.

DeepTaskGen produced consistently higher reconstruction performance than the linear baseline (Fig 1b, Supplementary Figure 1 and 2). It achieved the highest reconstruction performance on the RELATION REL (𝑟 = .*711*, 𝜎 = .*067*), WM PLACE (𝑟 = .*707*, 𝜎 = .*051*), RELATIONAL MATCH (𝑟 = .*697*, 𝜎 = .*071*), and SOCIAL TOM (𝑟 = .*697*, 𝜎 = .*059*) contrasts, while it yielded the lowest performance on the GAMBLING PUNISH-REWARD (𝑟 = .*106*, 𝜎 = .*080*), and RELATIONAL MATCH-REL (𝑟 = .*238*, 𝜎 = .*146*) contrasts. We also observed a notable difference between the contrast maps with the highest (RELATION REL) and lowest (GAMBLING PUNISH-REWARD) performance (𝑟 = .*605*). We achieved the highest reconstruction accuracy in the SOCIAL task, while the MOTOR task yielded the lowest performance. Notably, this trend was also observed in the retest scans, indicating significant variations in reliability across different task-based contrast maps. Additionally, DeepTaskGen (µ = .*516*, 𝜎 = .*167*), where *µ* and *σ* are the mean and standard deviation scores across 47 contrast maps), achieved significantly greater reconstruction performance than the linear model (µ = .*367*, 𝜎 = .*127*) in all 47 task contrasts, indicating an advantage of our network in predicting task contrast maps compared to the linear model after FDR correction (Supplementary Figures 1, 2). DeepTaskGen also performed well relative to retest scans. It significantly outperformed retest scans (µ = .*470*, 𝜎 = .*206*) in 29 task contrasts while achieving worse performance in 4 task contrasts. However, we found no significant difference between DeepTaskGen and retest scans in 14 task contrasts in terms of reconstruction performance. The improved performance of DeepTaskGen over retest scans can be attributed to scan-specific noise present in retest scans or shared session-specific information across both task-based and task-free brain activity retained by DeepTaskGen. The group-average contrast maps (µ = .*524*, 𝜎 = .*165*) achieved the highest scores in reconstruction performance, significantly surpassing DeepTaskGen in 31 tasks while 16 tasks showed no significant difference. However, these group average maps do not provide any inter-individual variance whatsoever. This emphasizes that reconstruction performance, although necessary, is not sufficient for downstream analyses such as prediction and biomarker development, which require individual-specific information.

DeepTaskGen achieved the highest discriminability, measured with normalized diagonality index, in the LANGUAGE MATH-STORY task (µ = .*025*, 𝜎 = .*008*), whereas the GAMBLING PUNISH-REWARD task yielded the lowest (µ = .*0008*, 𝜎 = .*002*). Consistently, the LANGUAGE task showed the highest discriminability (µ = .*016*, 𝜎 = .*010*), while the MOTOR task had the lowest (µ = .*006*, 𝜎 = .*006*), which was the task having lowest discriminability in retest scans. The same pattern of results was observed using the fingerprinting score (Supplementary Figure 2). Additionally, DeepTaskGen (µ = .*011*, 𝜎 = .*009*) consistently yielded higher discriminability than the linear model (µ = .*004*, 𝜎 = .*006*) in 40 of 47 task contrasts while providing similar discriminability in the remaining 7 task contrasts. This indicates our model’s significant advantage in retaining individual differences compared to the linear model. However, retest scans (µ = .*097*, 𝜎 = .*071*) showed the highest discriminability, suggesting they provide the most distinct task contrasts. However, this high discriminability may be influenced by individual-specific noise.

In summary, compared to various baseline methods, DeepTaskGen achieves a good balance between reconstruction performance and maintaining individual-specific variation. The test results for all 47 task contrasts are given in Supplementary Table 3-6. Although it was not the goal of this work to systematically compare surface-based approaches with volumetric approaches, we present error metrics from comparable surface-based deep learning approaches in Supplementary Figure 5 and 6. These show an advantage for volumetric approaches in terms of reconstruction, whereas surface-based approaches show better discriminability independent of model type.

### DeepTaskGen allows for generating contrast maps on an unseen dataset

Next, we evaluated DeepTaskGen’s performance in generating task contrasts on a dataset not included in its original training. To do this, we used the HCP-D dataset, which is ideal because it shares multiple tasks with the HCP-YA dataset used for training the model. This, in turn, allows us to assess the quality of the map generation against ground truth. To adapt the pre-trained model to the new dataset, we fine-tuned it using a single task contrast common to both datasets (see Methods). We then used the fine-tuned model to predict the task contrast that we did not use during fine-tuning. For example, the model for the GAMBLING REWARD contrast was fine-tuned using EMOTION FACES-SHAPES, and vice versa for EMOTION FACES-SHAPES. Furthermore, we assessed to what extent the trained model performed on an unseen dataset without fine-tuning, which we referred to as “No Finetune.”

Fig 1d depicts the reconstruction and discriminability performance of the models for the task contrasts mentioned above. Overall, fine-tuning improved discriminability according to the diagonality index, and had mixed results for reconstruction. In more detail, DeepTaskGen-based models showed significantly better reconstruction performance (i.e., correlation) than the linear model in the “GAMBLING REWARD” contrast (𝑡 = *10*.*225*, 𝑝_1000_ = .*004*, 𝛿 = .*340*) (Supplementary Table 7), while providing similar performance for EMOTION FACES-SHAPES. While the fine-tuned model with EMOTION FACES-SHAPES outperformed the non-fine-tuned model in the GAMBLING REWARD contrast (𝑡 = *3*.*675*, 𝑝_1000_ = .*005*, 𝛿 = .*142*), the fine-tuned model with GAMBLING REWARD performed significantly worse in the EMOTION FACES-SHAPES contrast (𝑡 = −*11*.*059*, 𝑝_1000_ = .*004*, 𝛿 = −.*624*) compared to the non-fine-tuned model. A similar pattern of results was observed for Dice AUC (Supplementary Figure 2).

Regarding discriminability, the fine-tuned model significantly outperformed both the non-fine-tuned (𝑡 = *4*.*392*, 𝑝_1000_ = .*003*, 𝛿 = .*333*) and linear models (𝑡 = *3*.*998*, 𝑝_1000_ = .*003*, 𝛿 = .*347*) in the EMOTION FACES-SHAPES task according to the diagonality score. For the the GAMBLING REWARD contrast, the fine-tuned model significantly surpassed the non-fine-tuned model (𝑡 = *5*.*743*, 𝑝_1000_ = .*003*, 𝛿 = .*357*), yet performed comparably relative to the linear model (𝑡 = *1*.*362*, 𝑝_1000_ = .*200*) (Supplementary Table 8). The reduced discriminability advantage of the fine-tuned model over the linear model in the GAMBLING REWARD task is likely due to dataset-specific differences in the GAMBLING task (see Discussion). The test results for the additional performance metrics are given in Supplementary Figure 3 and Supplementary Table 9, 10. Interestingly, fine tuned models show a lower discriminability under the fingerprinting score. This is not unexpected given they are sensitive to different aspects of the data. In most cases we prefer the diagonality index, since it is sensitive to magnitude differences, whilst the fingerprinting score is a ‘winner takes all’ approach.

In summary, DeepTaskGen-based models enable the prediction of trained task contrast maps on datasets lacking such maps. However, the benefit and optimal approach for fine-tuning varies depending on the specific task contrast and therefore should be done with care.

### Predicted task contrast maps achieve stronger or comparable performance in predicting demographic, cognitive, and clinical variables

Next, we sought to demonstrate the utility of synthetic task images for downstream prediction tasks within the UK Biobank dataset. To achieve this, we fine-tuned our model on the UK Biobank dataset using EMOTION FACES-SHAPES task contrast (µ_reconstruction_ = 0.428, σ_reconstruction_ = 0.140, discriminability = 0.189) and predicted seven representative contrasts from each task available in HCP-YA (Figs. 3-4).

**Fig. 3:**
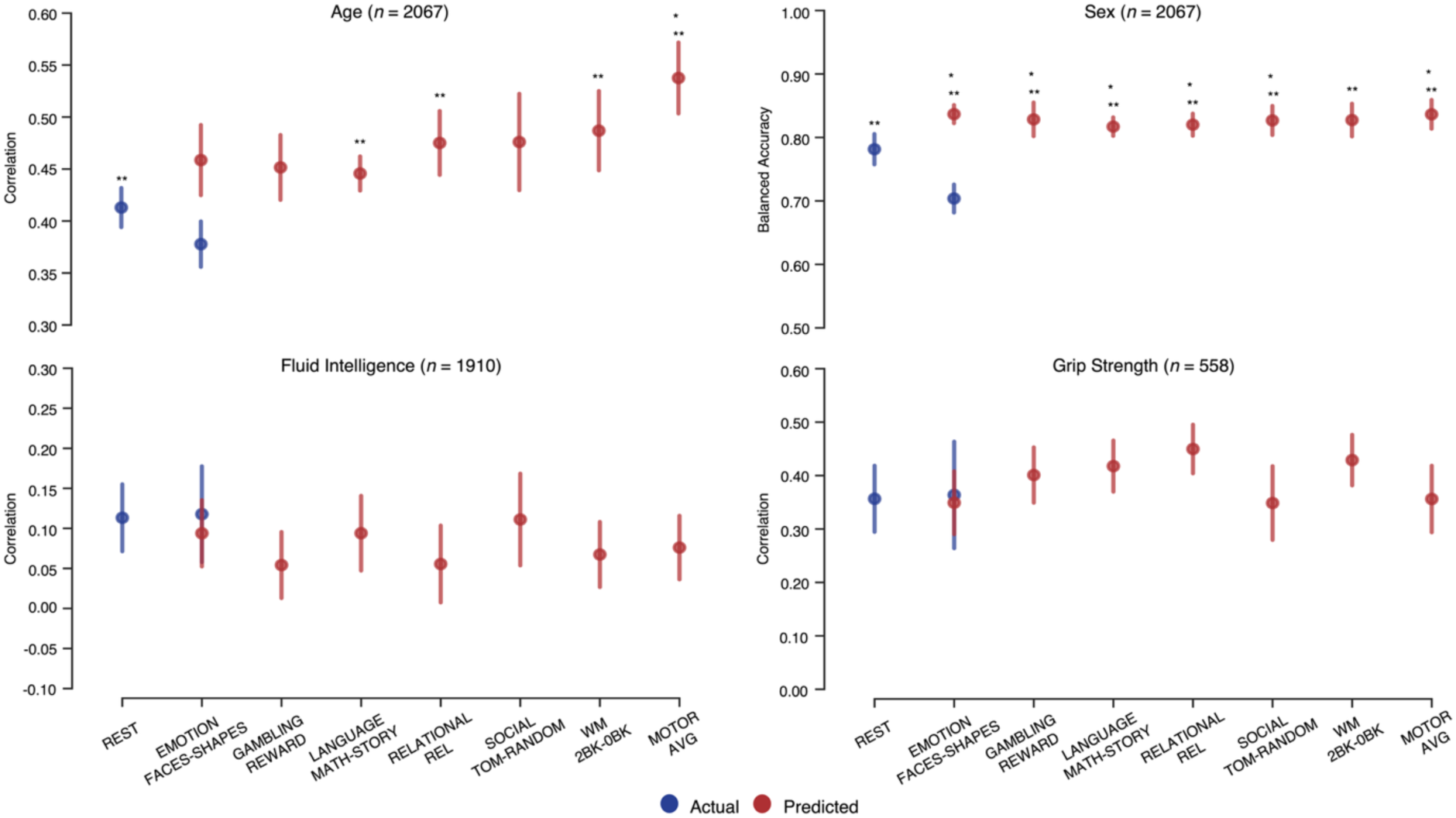
Prediction of subjects’ age, sex, fluid intelligence, and dominant hand grip strength using task contrast maps and resting-state connectome on UK Biobank. We predicted subjects’ demographic and cognitive measures in the UK Biobank dataset using three modalities: resting-state connectome, actual contrast map from the EMOTION task, and seven synthetic task contrast maps. All predictions were made using L2 regularized regression (i.e., ridge regression) within a 5-fold cross-validation framework with permutation testing (𝑃 = *1000*). Blue represents resting-state connectome and actual task contrast maps, while red represents predicted task contrast maps. Note that only the EMOTION task has both actual and predicted contrast maps on UKB, indicated in blue and red, while the remaining six tasks are indicated in red as UKB does not provide them. Significant predictions based on permutation testing are highlighted. Error bars indicate the standard deviation of prediction performance across five CV folds. Balanced accuracy was used for sex classification, while Pearson’s correlation assessed the other variables. Sample sizes for all analyses are indicated in each figure. An asterisk (*) indicates a significant difference in prediction performance between annotated maps and resting-state connectome data, while two asterisks (**) represent a significant difference between the annotated maps and actual task-contrast maps. Detailed test statistics are provided in Supplementary Tables 12, 14-17.

**Fig. 4:**
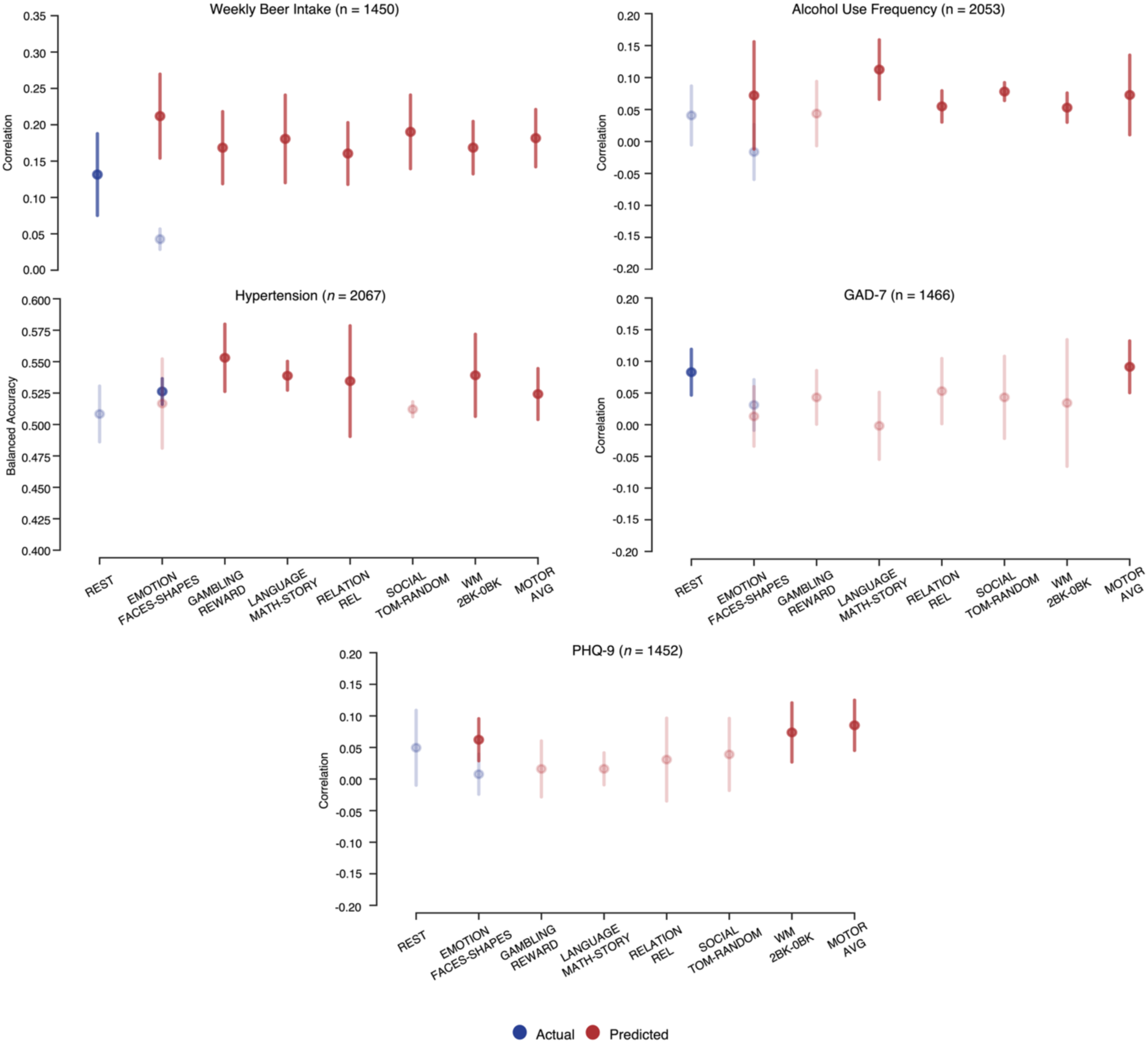
Prediction of subjects’ clinical measures using task contrast maps and resting-state connectome on UK Biobank. We predicted subjects’ clinical measures in the UK Biobank dataset using three modalities: synthetic task contrast maps, actual task contrast maps, and resting-state connectome data. resting-state connectome, actual contrast map from the EMOTION task, and seven synthetic task contrast maps. All predictions were made using L2 regularized regression (i.e., ridge regression) within a 5-fold cross-validation framework. Permutation testing (𝑃 = *1000*) was used to assess the significance of out-of-sample performance against a null distribution. Actual and synthetic brain measures are depicted in blue and red colors, respectively. Note that only the EMOTION task has both actual and predicted contrast maps on UKB, indicated in blue and red, while the remaining six tasks are indicated in red as UKB does not provide them. Significant predictions based on permutation testing are highlighted. Error bars indicate the standard deviation of prediction performance across five CV folds. Balanced accuracy was used to measure depression classification performance, while Pearson’s correlation was employed to assess other variables. Sample sizes for all analyses are indicated in each figure. Test statistics were only performed between predictions surviving permutation testing. The detailed test statistics are given in Supplementary Tables 12,18-20. The results for the additional clinical measures are depicted in Supplementary Figure 4.

To ensure that the predicted task contrast maps retain overall network structure and capture individual- and task-related variance for downstream analyses, we visualized them in two ways, along with the actual task contrast maps from the HCP-YA dataset as a reference. First, we generated group-average maps and observed excellent correspondence with the canonical networks for each task^24^ and the actual task contrasts (Fig. 2a). We also noted variability across datasets (e.g., reduced WM-related activation in UKB, which comprises mid and late adults, compared to other datasets). Then, we fit Uniform Manifold Approximation and Projection (UMAP)^25^ to each dataset to visualize high-dimensional differences between individuals and experimental tasks in a more intuitive two-dimensional space (Fig. 2b). We found that the predicted task contrast maps generated by DeepTaskGen contain lower within-task variance yet improved between-task variance compared to actual task contrast maps. This improvement can be attributed to the extensive denoising performed by our neural network during reconstruction, enhancing the signal-to-noise ratio (see Discussion).

We then compared the prediction performance of each of the seven predicted task contrast maps to that of actual task contrast maps and resting-state connectome data using permutation testing (𝑃 = *1000*). Specifically, we aimed to predict individuals’ demographic, cognitive, and clinical measures, including age, sex, fluid intelligence, dominant hand grip strength, overall health, alcohol use frequency, weekly beer intake, hypertension diagnosis, depression diagnosis, depressive symptoms (Patient Health Questionnaire [PHQ]-9^26^ and Recent Depressive Symptoms [RDS]-4^27^), anxiety symptoms (Generalized Anxiety Disorder Assessment [GAD]-7^28^) and neuroticism from predicted and actual task contrast maps, as well as resting-state connectome data. These were chosen because other studies have reported high prediction accuracy using these tasks^29,30^ and they are measures representing individuals’ mental health. The exact IDs for the variables are given in Supplementary 11.

The results, presented in Figures 3-4 and Supplementary Tables 12-20, show that the predicted task contrasts achieved greater or comparable prediction performance relative to actual EMOTION FACES-SHAPES task contrasts and resting-state connectome data for all variables. Specifically, we observed a large improvement in sex and age prediction using synthetic task contrasts, indicating that DeepTaskGen retains task activity patterns related to unique biological traits and amplifies their detection due to enhanced signal-to-noise ratios. Notably, for several clinically relevant measures, including RDS-4 (Supplementary Figure 4), PHQ-9, hypertension diagnosis, alcohol use frequency, synthetic task contrasts achieved significant prediction, while actual brain measures could not survive permutation testing. This enhanced performance indicates that synthetic task contrast maps can retain amplified complex activation patterns related to subjects’ clinical characteristics. Additionally, despite low prediction performance, only the actual EMOTION FACES-SHAPES withstand permutation testing for overall health prediction. None of the brain measures could survive permutation testing for the prediction of neuroticism and depression diagnosis (Supplementary Figure 4).

## Discussion

In this work, we present DeepTaskGen, a robust neural network architecture designed to predict task-based contrast maps using volumetric resting-state connectivity. This architecture enables the prediction of task contrasts on unseen datasets, even when task contrasts are unavailable or have not been acquired. We employed this method to generate task data for a comprehensive battery of cognitive tasks for the UK Biobank dataset and show that these synthetic task images retain individual-specific information essential for biomarker discovery.

We showed that our approach accurately predicts task-based contrast maps from resting-state connectivity in both the training dataset (HCP-YA) and unseen datasets (HCP-D and UKB). DeepTaskGen showed excellent reconstruction performance in that it outperformed retest scans in terms of more than half of the task contrasts and performed consistently better than a linear baseline model, which aligns with previous work^17^. Importantly, DeepTaskGen’s generated task contrasts preserve individual differences relative to linear models and group-average maps. This indicates that these synthetic contrast images may be better suited as biomarkers for individual-specific differences. Although group average maps show better reconstruction performance, they do not retain such individual-specific differences.

Notably, we observed variable performance when fine-tuning our model on HCP-D, which can be attributed to slight differences in the GAMBLING task between HCP-YA and HCP-D. Surprisingly, we found that non-fine-tuned models often perform as well or better than models fine-tuned for specific contrasts. More specifically, the GUESSING task is an adaptation of the GAMBLING task^31^, with modifications to make it more child-friendly and incorporate a magnitude manipulation^18^. This slight difference in the target task could result in performance loss for the fine-tuned model compared to the non-fine-tuned model in EMOTION. This may be because the slight difference in the adapted task could introduce shifts in feature representations, which the backbone may capture during fine-tuning, potentially leading to suboptimal transfer and reduced performance. Conversely, we observed a significant performance advantage of fine-tuning in GAMBLING, because EMOTION, which is the same in both datasets, was used for fine-tuning. Moreover, the slight task paradigm variation explains the lack of discriminability performance gain of the fine-tuned model over the baseline linear model in GAMBLING and the increased reconstruction performance discrepancy between datasets compared to EMOTION. Collectively, these findings underscore the critical role of domain similarity in fine-tuning outcomes and indicate that fine-tuning should be performed and evaluated carefully.

One of the key contributions of this work is a set of synthetic images in UKB for tasks that were not acquired. We made use of these to perform extensive validation analyses to showcase the advantage of using predicted task contrast maps for predicting individual differences in an extensive set of demographic, cognitive, and clinical variables. Our results indicate that these predicted maps offer equivalent or better performance in all the scenarios we evaluated relative to actual EMOTION FACES-SHAPE task contrast images and resting-state connectome data. This is likely because our approach increases the signal-to-noise ratio in the synthetic data by (i) reducing inherent noise in functional MRI data and (ii) allowing an increase in the temporal degrees of freedom from resting-state scans that are frequently longer than task-based scans. Inherent noise removal (i.e., denoising) occurs on several levels within the U-Net architecture, most notably on the downsampling (i.e., encoding) branch. Similar to other dimensionality reduction methods, the network compresses the information in the input image that is most relevant to the target image, suppressing unrelated information attributed to noise. During training, the weights of these blocks are constantly updated to force the network to extract the most relevant information to the target contrast maps, minimizing overall loss. Due to these inherent mechanisms, U-Net has been widely used to remove noise from biomedical images^32,33^. Moreover, we observed variability in the predictive performance of various synthetic task contrasts, which indicates DeepTaskGen’s capability to retain task-specific variability, also evidenced in the UMAP visualizations (Fig. 2b). This reinforces DeepTaskGen as a robust tool to facilitate predictive neuroscience.

Our method builds directly upon the foundational work of Tavor et al.^15^ and Ngo et al.^17^, who demonstrated the potential to predict task-based brain activity from resting-state brain connectivity. However, these previous methods have additional limitations. Tavor et al. employed a linear model, which fails to capture the complexity of the shared network architecture between task-based and task-free brain activity and does not effectively transfer learned parameters to unseen datasets. Ngo et al.^17^ introduced a neural network-based architecture, but it is limited to the cortical surface, thereby excluding subcortical areas and cerebellum that are crucial for cognitive functions^34–36^ and mental disorders^37–39^. In contrast, our proposed network, DeepTaskGen, overcomes these drawbacks, generating task contrast maps of the entire brain in new cohorts where such maps were not acquired. Additionally, Ngo et al.^17^ proposed a reconstructive-contrastive loss to enhance inter-subject variance in predicted contrast maps while minimizing within-subject differences. While the authors observed increased inter-subject variance, they noted its ineffectiveness for fine-tuning, therefore suggested using the mean squared error loss (MSE), which may still bias predictions toward the group average on target datasets. We also observed a similar pattern, where reconstructive performance decreased with increasing inter-subject contrast. To address this, we introduced Contrast-Regularized Reconstructive Loss (CR-R), which can be applied to both training and target samples without compromising reconstruction performance. It should be noted that even with these modifications, all methods are regularized toward predicting the group average, consistent with earlier findings^15,17,40^. Whilst this approach yields good reconstruction performance, it is clear that sufficient inter-individual variation remains for meaningful behavioral prediction, as shown in Figure 3 and in the diagonality index. Moreover, the improved generalizability to new cohorts and predict tasks that were not acquired is an important contribution of this work.

DeepTaskGen holds a wide range of implications in neuroscience studies. One significant application is the prediction of contrast maps for age groups for whom task data are difficult to acquire, such as children or individuals with neurological and psychiatric disorders. We showed that synthetic contrast maps facilitate predictions and biomarker studies within populations where acquiring task-based fMRI data may be challenging, which creates a significant potential to expand the scope of precision psychiatry. However, the correspondence between resting-state and task-evoked brain activation may vary across different groups, including those with disorders. Therefore, a model trained solely on healthy controls might require careful fine-tuning to ensure accurate predictions in disorder groups, which should be a focus of future studies. Another important application of DeepTaskGen lies in methods that utilize population data, such as normative modeling^41^. A recent study modeling the heterogeneity in the Emotional Face Matching Task using population-wide life-long normative modeling indicates a notable gap in mid-adulthood age groups^42^. In this case, we believe DeepTaskGen can play an important role in this context by augmenting contrast maps for the underrepresented age group in their database. Additionally, while the UKB dataset contributes a large portion of their data due to its sample size, it only includes the Emotion Face Matching Task, limiting the replication of such models for other fMRI tasks. DeepTaskGen can also address this challenge by generating corresponding synthetic task contrasts by requiring only resting-state fMRI data, which can then facilitate the development of more comprehensive normative models. Furthermore, our method can benefit large-scale imaging genetics studies, such as genome-wide association studies (GWAS), where potential associations between genotypes and brain-based phenotypes are studied. For instance, our approach allows pooling data to test for associations between putative genetic variants and specific task-evoked activity patterns. A recent study^43^ revealed significant genotype-phenotype associations using multimodal neuroimaging data from UKB, but it still faces the limitations mentioned above. Thus, DeepTaskGen can also enhance GWAS by enabling more specific and task-related associations using synthetic task contrasts. It is important to note the difference in variance distribution between predicted and actual task-based contrast maps. Actual images contain task-related variation (i.e., signal) and epistemic variation (or noise)^41^. During the prediction process, such noise can be effectively removed, amplifying the variance across experimental tasks by retaining only task-related variance. This can be seen in the UMAP visualizations in Fig. 2b, where predicted maps showed lower within-task contrast variance but higher between-task contrast variance than actual maps. Such removal of epistemic noise results in an increased signal-to-noise ratio, leading to improved prediction performance in our validation analyses. While this may not be critical for traditional statistical methods like group comparisons, it becomes more important when these images are used in complex models where noise is explicitly modeled. Therefore, we strongly recommend careful model designs that consider this difference when incorporating both data types within a single model. Additionally, one could utilize latent information in the resting-state connectome extracted by autoencoders to maximize phenotype prediction performance as this latent information hypothetically contains necessary details to generate task-based contrast maps. However, like prior methods, we provide synthetic contrast maps that can surpass the prediction performance of actual maps, as indicated in this study, while also providing synthetic maps spatially equivalent to actual maps. This ensures that synthetic images remain informative and straightforward to interpret, which is critical for several downstream analyses.

Several methodological considerations should be noted when interpreting our results. First, the spatial Independent Component Analysis (ICA) and task contrast maps provided in the HCP-YA dataset are surface-based (i.e., fs_LR surface). We projected these surface-based images to the volumetric MNI space using a well-established method, Registration Fusion^44^, which, despite its promise, might not be as optimal as direct registration to the coordinate system during preprocessing. Therefore, the projected contrast maps may have potential differences from those originally preprocessed in volumetric space, which might impact prediction performance. Second, despite being more reliable than traditional ICA approaches, group-level ICA maps may still not be optimally reliable due to their data-driven nature. This may be particularly important when group-level ICA maps are estimated separately on different datasets. However, in this study, as mentioned above, the same ICA map provided by HCP was used as the primary map to parcellate rs-fMRI images across datasets. Additionally, parcellation was done in a common MNI space to ensure reliability across datasets. Third, the different dimensionalities of ICA maps used to parcellate rs-fMRI might result in varying prediction performance. Although ICA maps with higher dimensionality could provide voxel-to-region of interest (ROI) connectomes with greater granularity, they would significantly increase the size of the connectome, rendering it computationally infeasible due to excessive GPU memory usage. Conversely, ICA maps with lower dimensionality could serve as a natural feature reduction method; however, the resulting voxel-to-ROI connectome might be too simplistic to retain the spatial information important for predicting task contrast maps. Although this is beyond the scope of the current study, we strongly believe that the effect of input dimensionality on model performance should be investigated in future studies. Forth, while HCP-YA is the largest open dataset with multimodal fMRI, the sample size may still be suboptimal to fully capture the true individualized patterns between rest- and task-based fMRI^45^. UKB offers a significantly larger sample size, including subjects recorded multiple times, which can serve as a reference for test-retest reliability. Although this is beyond the scope of the present study, which aims to present a model that can generate multi-domain synthetic task-based contrasts, evaluating generative models on such a large sample will be necessary in future studies. Moreover, as described in the Methods section, the presented fine-tuning routine is most effective when the latent brain network architecture underlying task activations and resting-state connectome remains relatively stable between the source and target datasets. When applied across substantially different age groups, model transferability might be reduced due to the evolution of cognition and brain architecture over developmental stages, leading to varying brain activation patterns under cognitive tasks^46^. In such cases, fine-tuning the entire network without freezing the final layer could provide a better adaptation strategy, allowing the model to learn age-specific modifications between the task-related final layer and the brain network-related backbone layers, while leveraging shared functional patterns. However, as this is not a direct task activation transfer, researchers should interpret the results more cautiously than those obtained with the presented fine-tuning routine. Another option would be to incorporate multi-task learning by adding auxiliary tasks, such as age prediction, during training and fine-tuning. This approach might force the model to retain age-related information, as has been evidenced by a previous study^47^. Furthermore, while methods using volumetric images provide varying reconstruction performance (depending on task contrast), they exhibit lower discriminability compared to surface-based methods (Supplementary Figure 5,6). Notably, this reduction is not specific to DeepTaskGen but is a characteristic of volumetric-based methods, as DeepTaskGen demonstrated a greater performance gain over the baseline linear model compared to surface-based generative methods (Supplementary Figure 7,8). This difference between surface and volume can be attributed to factors such as variations in preprocessing and brain representation (e.g., surface vs. volume, total number of vertices vs. voxels). Researchers should select surface- or volumetric-based generative methods based on their research scope (e.g., subcortical involvement) and data availability. Additionally, future research should explore methods to reduce the disparity between surface and volumetric data.

In summary, we have demonstrated DeepTaskGen, a general-purpose tool for generating arbitrary synthetic task images from resting-state data. We consider that this will facilitate the study of individual differences and the generation of task-related biomarkers.

## Methods

### Datasets

Supplementary Table 2 provides the sample details and functional scan acquisition parameters for the datasets used in this study. Further preprocessing details are provided in the supplementary material.

#### Human Connectome Project Young-Adult (HCP-YA)

The Human Connectome Project Young-Adult (HCP-YA) 1200 Release was used as a main dataset to train and evaluate models^11^. 958 participants comprising all the task and resting-state sessions were included. Of those 958 subjects, 39 have re-test 3T fMRI data from the second visit, which we used as an evaluation set. Rs-fMRI data were acquired in four 15-minute runs, with 1200 timepoints per run, totaling 4800 timepoints per subject. However, since fewer timepoints (e.g., 300-500) are common in neuroimaging datasets and stable functional connectivity can be computed with shorter resting-state acquisitions^22,23^, we divided resting-state data into eight contiguous parts (600 timepoints) as previously shown^17^. Similar to previous studies^15,17^, we used ROIs from the 50-component parcellation derived from spatial ICA released within HCP-YA to compute individual functional connectivity. As tb-fMRI data, we used 47 unique contrasts from 7 main task domains^24^: WORKING MEMORY, EMOTION, SOCIAL, RELATIONAL, LANGUAGE, MOTOR, and GAMBLING. We used minimally preprocessed and FIX-cleaned fMRI data provided by HCP^48^. While HCP also provides volumetric rs-fMRI data, the spatial ICA maps, and z-transformed task-activation maps are only surface-based. Therefore, we projected surface-based data to volumetric space using the registration fusion (RF-ANTs) method^44^. Registration Fusion is a popular method that allows nonlinear mapping between MNI152 and fsaverage coordinate systems using coordinate mapping computed on 745 subjects from the Brain Genomics Superstruct Project (GSP)^49^. The details of the mapping process are given elsewhere^44^. Briefly, we separated CIFTI files into a cortical surface in fs_LR space (wb_command –cifti-separate) and a subcortex in MNI space, mapped the cortical surface into fsaverage space (wb_command –metric-resample) using HCP Workbench Command^50^. We then projected it into volumetric space using registration fusion and combined it with the subcortex to create a whole-brain volumetric image. Projected contrast maps were then visually checked for alignment to MNI space and corresponding task activation patterns. Note that the projected images only contain gray matter. We also binarize a reference projected image (melodic_IC.dscalar.nii, containing ICA components) to use as a gray matter mask. Subsequently, the scan images were downsampled to 2×2×2 mm voxel resolution and cropped to the cortex (resulting image dimension is 76×93×78) to reduce the computational costs.

#### Human Connectome Project Development (HCP-D)

The evaluation of DeepTaskGen’s generalizability across different populations with various age groups and transferability of learned parameters to unseen datasets was conducted on the HCP-Development (HCP-D) dataset from the HCP-Lifespan project^18^. 637 subjects that had completed four resting-state runs (488 timepoints per run) and Emotion^1^ and Guessing^31^ tasks were included in the study. The choice of Emotion and Guessing tasks was due to the Emotion task performed in both the main HCP-YA and HCP-D projects, while the Guessing task served as an adaptation of the Gambling task in HCP-YA, with some modifications.

#### UK Biobank (UKB)

To assess the practical applicability of generated contrast maps and further generalizability of the trained model on a large-scale sample, we used the UK Biobank dataset^19^. A total of 20,792 subjects with resting-state (490 timepoints) and Emotion task^1^ MRI images were included in this study.

### Voxel-to-ROI functional connectivity

DeepTaskGen employs resting-state voxel-to-ROI connectivity to predict task contrasts. Voxel-to-ROI connectivity is computed as Pearson’s correlation between each voxel’s time series and the average signal of the target ROI^51^. The target ROIs are derived from components obtained through group-level independent component analysis (ICA) applied to resting-state fMRI time series^52^. Voxel-to-ROI connectivity provides an increased spatial resolution compared to the conventional ROI-to-ROI approach, which requires deeper models at the generation step. Although voxel-to-voxel connectivity could produce a higher spatial resolution than voxel-to-ROI connectivity, it is computationally infeasible due to excessive GPU memory use. Additionally, as mentioned earlier, the voxel-to-ROI approach allows for modeling the connectivity of the entire brain, providing a more inclusive solution than the vertex-to-ROI approach.

### DeepTaskGen

The architecture of DeepTaskGen is depicted in Fig 1. DeepTaskGen is a volumetric adaptation of BrainSurfCNN^17^, a surface-based convolutional neural network based on U-Net architecture^53^ widely used in biomedical image segmentation with additional attention mechanism^54^. Like autoencoders, U-Net consists of encoding (i.e., downsampling), decoding (i.e., upsampling) blocks, a bottleneck layer and skip connections. Specifically, DeepTaskGen’s network architecture is as follows: each block in the encoding and decoding arms consists of a 3D convolutional layer, batch normalization, and non-linear ReLU activation function. Additionally, max pooling (MaxPool3d) is applied before each encoding block to reduce the dimensions of the input image by half by retaining task-related information. In contrast, compressed images are upsampled before each decoding block to match the output dimensionality with actual maps. In addition to BrainSurfCNN^17^, we incorporated an attention gate into each skip connection. This enhancement helps the network focus on relevant features from the encoding blocks while suppressing irrelevant activations, thereby improving overall model sensitivity^54^. The details of the architecture are provided in Supplementary Table 1.

### Contrast-Regularized Reconstructive Loss

Unlike the previous study^17^, we did not utilize the reconstructive-contrastive (RC) loss in model training and fine-tuning. Although RC loss enhances the variance among subjects’ generated contrast maps, it can reduce reconstruction performance by over-emphasizing the contrastive component, particularly in transfer samples with smaller datasets, as reported by the authors^17^, who proposed the loss. Instead, we minimize the Contrast-Regularized Reconstructive Loss (CR-R) loss, in which we regularize the contrastive part of the reconstruction loss (RC) and further combine it with the reconstructive loss (i.e., mean squared error, MSE) to prevent the model from losing focus on increasing reconstruction performance. Given a mini-batch of *N* subjects, 𝐵 = {𝑥*_i_* … 𝑥*_N_*}, in which 𝑥*_i_* is the actual contrast maps of subject 𝑖, the CR-R loss 𝐿*_CR-R_* is defined as:

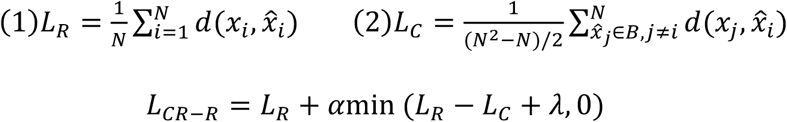

where 𝑑(.) is L2 norm, 𝐿*_R_* is the reconstructive loss calculated as mean-squared error (MSE) between actual and generated contrast maps, and 𝐿*_c_* is the contrastive loss calculated as MSE between subjects generated contrast maps and actual contrast maps of the remaining subjects. 𝐿*_CR-R_* encourages 𝐿*_c_* to be larger than 𝐿*_R_* by a margin 𝜆 and further combines it with 𝐿*_R_* to prevent the model from reducing reconstruction performance while increasing inter-subject contrast. The hyperparameter 𝛼 controls the weight of contrastive part of the in this combined objective. Thanks to its integrated nature, CR-R does not require separate training sessions with MSE and RC loss, further simplifying the training process.

### Model Training

DeepTaskGen was trained on HCP-YA (𝑛*_train_* = 827, 𝑛*_validation_* = 92, 𝑛*_test_* = 39) for 100 epochs using a batch size of 10, CR-R as the loss function (𝜆 = *1* and 𝛼 = *0*.*25*), and the Adam optimizer.

As the task contrasts maps projected from fs_LR space (see Supplementary Material for detailed descriptions) and includes only gray matter, we used only these voxels to minimize the loss function. The model training was conducted on a single Nvidia A100-40GB GPU and took approximately 6 hours.

### Model transfer to HCP-D and UKB

Our main objective for the HCP-D dataset was to validate the performance of DeepTaskGen to predict unavailable arbitrary task contrast maps on an unseen dataset. For instance, we fine-tuned the model using the EMOTION FACES-SHAPES contrast to predict the GAMBLING REWARD contrast. This approach relies on the assumption that the weights in the trained model’s output layer are specific to a given task contrast (e.g., GAMBLING REWARD), while the backbone layers (i.e., all the layers before the output layer) contain information shared across multiple task contrasts yet differ across datasets due to dataset-related features. We also assume that the task-specific output layer is similar across datasets, as it primarily processes task-specific high-level features using dataset-specific low- and mid-level features, such as variations in MRI acquisitions or preprocessing, from the backbone layers, whereas the backbone layers differ due to dataset-specific differences. By adapting only the backbone of the model, we can effectively predict the trained contrasts from HCP-YA. To achieve this, we froze the parameters of our trained model’s output layer and fine-tuned the backbone of our model on the HCP-D dataset using either GAMBLING-REWARD or EMOTION FACES-SHAPES contrast. Specifically, we fine-tuned the pre-trained model using only a single task contrast (e.g., GAMBLING-REWARD) for 50 epochs. We used 515 and 58 subjects for training and validation during fine-tuning, respectively. Subsequently, we replaced the output layer of the fine-tuned model with one that matched the target task contrast (e.g., EMOTION FACES-SHAPES) and predicted the target task contrast maps for 64 test subjects (i.e., holdout set). The model transfer performance of DeepTaskGen was also compared with that of a non-fine-tuned model and linear regression.

The initially trained model was further fine-tuned on UKB (𝑛*_train_* = 16840, 𝑛*_validation_* = 1872, 𝑛*_test_* = 2080) to test the practical applicability of predicted task contrast maps. We fine-tuned the backbone of the trained model using the EMOTION FACES-SHAPES task contrast available in UKB for 50 epochs. Then, we replaced the coefficients in the output layer of the fine-tuned model with corresponding model coefficients of the 47 task contrasts available in the training dataset to predict these task contrast maps for 2080 test subjects. Notably, despite being significantly larger than HCP-YA in terms of sample size, the UK Biobank (UKB) was not utilized for pre-training the main model, as it only provides one fMRI task, EMOTION, which is insufficient for training a multi-modal model.

### Baselines

#### Group-average contrasts

The group-average contrasts were utilized as a reference to represent the features in the task contrast that are shared among individuals. A simple predictive model can effectively capture these common features. By comparing the performance of the predicted task contrasts with the group-average contrasts, we can assess the degree to which the predicted task contrasts capture individual-specific information beyond what is captured by the common features represented in the group-average contrasts. It could also show the degree to which variation would be in each task contrast, e.g., a high correlation between true and group-average task contrasts indicates a reduced level of inter-individual variation.

#### Retest scans

The additional tb-fMRI data acquired in a second run (𝑛 = *39*) was used to evaluate the reliability of the task contrasts.

#### Linear Model

We implemented this baseline model based on the approach presented by Tavor and colleagues^15^, as it is the pioneering study in predicting task-based contrast maps from resting-state connectivity. While the original method was developed and tested using surface brain images (i.e., CIFTI files), we adapted it into volumetric space for an unbiased comparison. Within each of 50 ROIs (defined using ‘melodic_IC_ftb.dlabel.nii’, provided by HCP, projected to MNI using registration fusion), we vectorized the input resting-state connectomes and target contrast maps. A linear regression model was fit to map between input and target vectors, restricted to these regions containing only gray matter (157,461 voxels) to prevent nuisance effects of white matter and CSF signals. Fitting separate linear models to each ROIs resulted in a large weight matrix with dimensions R x S, where R represents the number of ROIs and S represents the number of subjects in the training set and validation set combined (𝑛 = *919*). We averaged this weight matrix across subjects to create a vector containing average weights for ROIs. We then predicted the corresponding task contrast maps for subjects in the test set using this weight vector. This procedure was repeated for each of the 47 task contrasts, resulting in 2.159.650 linear models fit during training.

### Visualization of group-level contrast maps and inter-subject and inter-task similarity

To evaluate the structure of the predicted task contrast maps, we visualized both common structural elements across subjects (Figure 2a) and similarities or differences between subjects for various task contrasts (Figure 2b). Figure 2a displays the common brain structure by presenting the voxel-based sample average of the task contrast maps across all subjects.

We employed a non-linear dimensionality reduction method, Uniform Manifold Approximation and Projection (UMAP)^25^, to visualize the similarity or variance between subjects and across various tasks. UMAP preserves global and local topological structure while embedding high-dimensional data into a lower-dimensional space. Specifically, UMAP was utilized to project the high-dimensional task-based contrast maps into a two-dimensional space (i.e., two components), where each dot represents a subject’s task contrast map. This allows for a straightforward interpretation of similarity or dissimilarity between subjects and fMRI tasks by considering the distance between dots on the two-dimensional plot. We implemented UMAP in Python using the umap-learn package with default hyperparameters (𝑛*_neighbors_* = *15*, 𝑚𝑒𝑡𝑟𝑖𝑐 = 𝑒𝑢𝑐𝑙𝑖𝑑𝑒𝑎𝑛). To retain and visualize the global and topological details and achieve optimal embedding within each dataset, we fit a UMAP model to each dataset separately.

### Prediction of cognitive and demographic data for validation

To validate the clinical relevance of our model, we predicted multiple non-imaging variables using the synthetic images derived from the UK Biobank dataset. Specifically, we aimed to predict three main domains: physical (age, sex, grip strength of the dominant hand, overall health), cognitive (fluid intelligence), and mental health (alcohol use frequency, weekly beer intake, hypertension diagnosis, depression diagnosis and symptoms (PHQ-9^26^ and RDS-4^27^), anxiety symptoms (GAD-7^28^), and neuroticism). These phenotypes cover key aspects of individuals’ demographics and well-being and have been shown to associate brain biomarkers^55^. Additionally, they have been extensively analyzed in the UK Biobank dataset, further supporting their relevance^27,29,30,56^. We used a similar L2 regularized linear model (i.e., ridge regression) and training and evaluation procedure for all these prediction analyses (see below). The exact sample size for each prediction is provided in Figures 3-4.

All regression models were built using predicted contrast maps for seven main task contrasts (EMOTION FACES-SHAPES, GAMBLING REWARD, WM 2BK-0BK, LANGUAGE MATH-STORY, RELATION REL, SOCIAL TOM-RANDOM, MOTOR AVG) as well as subjects’ actual EMOTION FACES-SHAPES contrast maps and resting-state connectome as comparison. The resting-state connectome for each subject was computed as Pearson’s correlation between each time series extracted using the 50-component ICA parcellation described earlier (*50*𝑥*50* correlation matrix for each individual). Feature matrices in prediction analyses were constructed by taking all gray matter voxel values within the brain (155,680 features) or extracting the upper triangle of the correlation matrix (1,225 features) for each subject. We predicted variables using L2 regularized linear regression. To mitigate the possible effect of the difference in the number of features derived from task-based contrast maps and resting-state connectome, the alpha value, which controls regularization strength, was set to a value computed by dividing the number of features by the training sample size. This resulted in significantly more regularization for task-based features than connectome-based features. The 5-fold cross-validation was used to evaluate models’ out-of-sample performance (i.e., generalizability), while permutation testing (𝑃 = *1000*) was used to estimate the significance of models’ performance by testing against a null distribution. Additionally, using permutation testing, we compared the predictive performance of synthetic contrast maps with actual contrast maps and resting-state connectome. Specifically, we predicted variables using true labels and compared 5-fold CV performance using a paired t-test. To create a null sampling distribution for the *t*-values, we shuffled labels and repeated the comparisons 1000 times. Then, we computed the p-value as the proportion of permutations resulting in t-values as extreme as or more extreme than the observed t-values. Note that predictions were performed on the 2080 subjects we spared as a test set during the fine-tuning process explained earlier.

### Measures of model performance

#### Reconstruction performance

Reconstruction performance is one of the primary measures for evaluating a model’s performance. It is determined by calculating Pearson’s correlation between the predicted and actual task contrast maps for unseen subjects (i.e., those not used in training) in the test set.

#### Diagonality Index

While reconstruction performance is essential for evaluating a model’s capability to generate task contrast maps, it is insufficient for assessing its ability to preserve inter-individual differences in these synthetic maps, which is crucial for downstream biomarker development and individual-specific predictions. To address this, we computed the diagonality index^17,20,22^, that measures the difference between the correlations of subjects’ actual and predicted contrast maps (on-diagonal) and the mean correlation between subjects’ predicted maps and the actual maps of other subjects (off-diagonal). However, the raw diagonality index is not directly comparable across methods, as it only reflects the ratio of cases where the correlation between synthetic and actual maps is highest, without accounting for overall map similarity. To ensure comparability, we normalized the diagonality index by dividing it by the on-diagonal correlation scores (i.e., reconstruction performance).

#### Prediction performance

The predictive power of both actual and predicted task contrast maps was evaluated using Pearson’s correlation coefficient for the dominant hand’s grip strength, brain age, fluid intelligence, and overall health. For sex classification, balanced accuracy, defined as the arithmetic mean of the true positive rate and true negative rate, was used as the evaluation metric.

### Statistics and Reproducibility

The Human Connectome Project Young Adult (HCP-YA) and Developmental (HCP-D) datasets comprise 958 and 637 subjects, respectively, while the UK Biobank dataset includes 20,792 subjects. Statistical analyses and reproducibility details for model training and evaluation are provided in the respective sections of the Results and Methods. Performance comparisons of methods on HCP-YA and HCP-D were conducted using permutation-tested *t*-tests (1000 permutations) on the corresponding test samples (HCP-YA: N = 39; HCP-D: N = 64). Significance was set at *p* < 0.05, with False Discovery Rate (FDR) correction applied across all comparison pairs and task contrasts. Effect sizes for significant tests were quantified using Cliff’s Delta (*δ*). Behavioral, cognitive, demographic, and clinical predictions were performed on the UK Biobank test set, with sample sizes indicated above each figure (Figs. 3, 4, Supplementary Figure 4). Prediction performance and significance were evaluated using 5-fold cross-validation and permutation testing (1000 permutations), and average performance is reported in the manuscript. Comparisons of prediction performance between synthetic and actual task-based images were conducted using permutation-tested *t*-tests (1000 permutations), with FDR correction applied across all comparison pairs and task contrasts. Effect sizes for significant tests were again measured using Cliff’s Delta (*δ*). To promote transparency and reproducibility, the complete implementation, including model architecture, training, evaluation, and downstream validation steps, as well as pre-trained models are provided in an open-source repository.

### Technology use disclosure

Large language models were used for proofreading (e.g., grammar, typos, integrity, and clarity checks) and as assistance during Python programming. All authors have read, corrected, and verified all information presented in this manuscript and Supplementary Information.

## Data availability

The minimally preprocessed HCP-YA S1200 data can be accessed and downloaded from the following link: https://www.humanconnectome.org/study/hcp-young-adult/document/1200-subjects-data-release. The ICA-based group-averaged parcellations used to construct resting-state connectomes are available at: https://www.humanconnectome.org/study/hcp-young-adult/document/extensively-processed-fmri-data-documentation. Similarly, preprocessed data for the HCP-D study can be downloaded from https://www.humanconnectome.org/study/hcp-lifespan-development/data-releases. The UK Biobank (UKB) dataset is publicly available at https://www.ukbiobank.ac.uk/.

## Code availability

DeepTaskGen model code can be found on GitHub: https://github.com/eminSerin/DeepTaskGen. Python scripts to reproduce all the results presented in this paper are given at https://github.com/eminSerin/deeptaskgen-paper. Additionally, trained models to generate task contrast maps on various open datasets are publicly available at https://github.com/eminSerin/deeptaskgen-models.

## Acknowledgement

Funded by the European Union (Grant agreement No 101057429). Complementary funding was received by UK Research and Innovation (UKRI) under the UK government’s Horizon Europe funding guarantee (10131373 and 10038599) and the National Key R&D Program of Ministry of Science and Technology of China (MOST 2023YFE0199700). Views and opinions expressed are however those of the author(s) only and do not necessarily reflect those of the European Union, the European Health and Digital Executive Agency (HADEA), UKRI or MOST. Neither the European Union nor HADEA nor UKRI nor MOST can be held responsible for them. This work has also been supported by a Grant from the German Research Foundation to the ENIGMA task-based fMRI Working Group (DFG ER 724/4–1, WA 1539/11–1). Data used in this study were provided in part by the Human Connectome Project, WU-Minn Consortium (principal investigators: D. Van Essen and K. Ugurbil; grant number 1U54MH091657), funded by the 16 National Institutes of Health (NIH) institutes and centers supporting the NIH Blueprint for Neuroscience Research, and by the McDonnell Center for Systems Neuroscience at Washington University. Additionally, this research utilized data obtained from UK Biobank, a large-scale biomedical database.

## References

1. Hariri, A. R., Bookheimer, S. Y. & Mazziotta, J. C. Modulating emotional responses: effects of a neocortical network on the limbic system. Neuroreport 11, 43–48 (2000).

2. Swartz, J. R., Knodt, A. R., Radtke, S. R. & Hariri, A. R. A neural biomarker of psychological vulnerability to future life stress. Neuron 85, 505–511 (2015).

3. Gao, S. Combining multiple connectomes improves predictive modeling of phenotypic measures. 9 (2019).

4. Greene, A. S., Gao, S., Scheinost, D. & Constable, R. T. Task-induced brain state manipulation improves prediction of individual traits. Nat. Commun. 9, 2807 (2018).

5. Gal, S., Coldham, Y., Tik, N., Bernstein-Eliav, M. & Tavor, I. Act natural: Functional connectivity from naturalistic stimuli fMRI outperforms resting-state in predicting brain activity. NeuroImage 258, 119359 (2022).

6. Sripada, C., Angstadt, M., Rutherford, S., Taxali, A. & Shedden, K. Toward a ‘treadmill test’ for cognition: Improved prediction of general cognitive ability from the task activated brain. Hum. Brain Mapp. 41, 3186–3197 (2020).

7. Jiang, R. et al. Task-induced brain connectivity promotes the detection of individual differences in brain-behavior relationships. NeuroImage 207, 116370 (2020).

8. Shah, L. M., Cramer, J. A., Ferguson, M. A., Birn, R. M. & Anderson, J. S. Reliability and reproducibility of individual differences in functional connectivity acquired during task and resting state. Brain Behav. 6, e00456 (2016).

9. Tetereva, A., Li, J., Deng, J. D., Stringaris, A. & Pat, N. Capturing brain-cognition relationship: Integrating task-based fMRI across tasks markedly boosts prediction and test-retest reliability. NeuroImage 263, 119588 (2022).

10. Woo, C.-W., Chang, L. J., Lindquist, M. A. & Wager, T. D. Building better biomarkers: brain models in translational neuroimaging. Nat. Neurosci. 20, 365–377 (2017).

11. Van Essen, D. C. et al. The WU-Minn Human Connectome Project: An overview. NeuroImage 80, 62–79 (2013).

12. Schumann, G. et al. The IMAGEN study: reinforcement-related behaviour in normal brain function and psychopathology. Mol. Psychiatry 15, 1128–1139 (2010).

13. Casey, B. J. et al. The Adolescent Brain Cognitive Development (ABCD) study: Imaging acquisition across 21 sites. Dev. Cogn. Neurosci. 32, 43–54 (2018).

14. Cole, M. W., Ito, T., Bassett, D. S. & Schultz, D. H. Activity flow over resting-state networks shapes cognitive task activations. Nat. Neurosci. 19, 1718–1726 (2016).

15. Tavor, I. et al. Task-free MRI predicts individual differences in brain activity during task performance. Science 352, 216–220 (2016).

16. Smith, S. M. et al. Correspondence of the brain’s functional architecture during activation and rest. Proc. Natl. Acad. Sci. U. S. A. 106, 13040–13045 (2009).

17. Ngo, G. H., Khosla, M., Jamison, K., Kuceyeski, A. & Sabuncu, M. R. Predicting individual task contrasts from resting-state functional connectivity using a surface-based convolutional network. NeuroImage 248, 118849 (2022).

18. Somerville, L. H. et al. The Lifespan Human Connectome Project in Development: A large-scale study of brain connectivity development in 5–21 year olds. NeuroImage 183, 456–468 (2018).

19. Miller, K. L. et al. Multimodal population brain imaging in the UK Biobank prospective epidemiological study. Nat. Neurosci. 19, 1523–1536 (2016).

20. Zheng, Y.-Q. et al. Accurate predictions of individual differences in task-evoked brain activity from resting-state fMRI using a sparse ensemble learner. NeuroImage 259, 119418 (2022).

21. Cohen, A. D., Chen, Z., Parker Jones, O., Niu, C. & Wang, Y. Regression-based machine-learning approaches to predict task activation using resting-state fMRI. Hum. Brain Mapp. 41, 815–826 (2020).

22. Tik, N. et al. Generalizing prediction of task-evoked brain activity across datasets and populations. NeuroImage 276, 120213 (2023).

23. Finn, E. S. et al. Functional connectome fingerprinting: Identifying individuals using patterns of brain connectivity. Nat. Neurosci. 18, 1664–1671 (2015).

24. Barch, D. M. et al. Function in the human connectome: Task-fMRI and individual differences in behavior. NeuroImage 80, 169–189 (2013).

25. McInnes, L., Healy, J. & Melville, J. UMAP: Uniform Manifold Approximation and Projection for Dimension Reduction. (2018) doi:10.48550/ARXIV.1802.03426.

26. Kroenke, K., Spitzer, R. L. & Williams, J. B. W. The PHQ-9: Validity of a brief depression severity measure. J. Gen. Intern. Med. 16, 606–613 (2001).

27. Dutt, R. K. et al. Mental health in the UK Biobank: A roadmap to self-report measures and neuroimaging correlates. Hum. Brain Mapp. 43, 816–832 (2022).

28. Spitzer, R. L., Kroenke, K., Williams, J. B. W. & Löwe, B. A Brief Measure for Assessing Generalized Anxiety Disorder: The GAD-7. Arch. Intern. Med. 166, 1092 (2006).

29. Gong, W., Bai, S., Zheng, Y.-Q., Smith, S. M. & Beckmann, C. F. Supervised Phenotype Discovery From Multimodal Brain Imaging. IEEE Trans. Med. Imaging 42, 834–849 (2023).

30. He, T. et al. Meta-matching as a simple framework to translate phenotypic predictive models from big to small data. Nat. Neurosci. 25, 795–804 (2022).

31. Delgado, M. R., Nystrom, L. E., Fissell, C., Noll, D. C. & Fiez, J. A. Tracking the hemodynamic responses to reward and punishment in the striatum. J. Neurophysiol. 84, 3072–3077 (2000).

32. Zhang, J., Niu, Y., Shangguan, Z., Gong, W. & Cheng, Y. A novel denoising method for CT images based on U-net and multi-attention. Comput. Biol. Med. 152, 106387 (2023).

33. Meng, Z., Fan, Z., Zhao, Z. & Su, F. ENS-Unet: End-to-End Noise Suppression U-Net for Brain Tumor Segmentation. in 2018 40th Annual International Conference of the IEEE Engineering in Medicine and Biology Society (EMBC) 5886–5889 (IEEE, Honolulu, HI, 2018). doi:10.1109/EMBC.2018.8513676.

34. Koziol, L. F. & Budding, D. E. Subcortical Structures and Cognition. (Springer New York, New York, NY, 2009). doi:10.1007/978-0-387-84868-6.

35. Janacsek, K. et al. Subcortical Cognition: The Fruit Below the Rind. Annu. Rev. Neurosci. 45, 361–386 (2022).

36. Stoodley, C. J. The Cerebellum and Cognition: Evidence from Functional Imaging Studies. The Cerebellum 11, 352–365 (2012).

37. Okada, N. et al. Subcortical volumetric alterations in four major psychiatric disorders: a mega-analysis study of 5604 subjects and a volumetric data-driven approach for classification. Mol. Psychiatry (2023) doi:10.1038/s41380-023-02141-9.

38. Ho, T. C. et al. Subcortical shape alterations in major depressive disorder: Findings from the ENIGMA major depressive disorder working group. Hum. Brain Mapp. 43, 341–351 (2022).

39. Hoppenbrouwers, S. S., Schutter, D. J. L. G., Fitzgerald, P. B., Chen, R. & Daskalakis, Z. J. The role of the cerebellum in the pathophysiology and treatment of neuropsychiatric disorders: A review. Brain Res. Rev. 59, 185–200 (2008).

40. Kwon, J. et al. Predicting task-related brain activity from resting-state brain dynamics with fMRI Transformer. Imaging Neurosci. 3, imag_a_00440 (2025).

41. Marquand, A. F. et al. Conceptualizing mental disorders as deviations from normative functioning. Mol. Psychiatry 24, 1415–1424 (2019).

42. Savage, H. S., et al. Unpacking the Functional Heterogeneity of the Emotional Face Matching Task: A Normative Modelling Approach. http://biorxiv.org/lookup/doi/10.1101/2023.03.27.534351 (2023) doi:10.1101/2023.03.27.534351.

43. Elliott, L. T. et al. Genome-wide association studies of brain imaging phenotypes in UK Biobank. Nature 562, 210–216 (2018).

44. Wu, J. et al. Accurate nonlinear mapping between MNI volumetric and FreeSurfer surface coordinate systems. Hum. Brain Mapp. 39, 3793–3808 (2018).

45. Schulz, M.-A., Bzdok, D., Haufe, S., Haynes, J.-D. & Ritter, K. Performance reserves in brain-imaging-based phenotype prediction. Cell Rep. 43, 113597 (2024).

46. Rubia, K. Functional brain imaging across development. Eur. Child Adolesc. Psychiatry 22, 719–731 (2013).

47. Zabihi, M. et al. Nonlinear latent representations of high-dimensional task-fMRI data: Unveiling cognitive and behavioral insights in heterogeneous spatial maps. PLOS ONE 19, e0308329 (2024).

48. Glasser, M. F. et al. The minimal preprocessing pipelines for the Human Connectome Project. NeuroImage 80, 105–124 (2013).

49. Holmes, A. J. et al. Brain Genomics Superstruct Project initial data release with structural, functional, and behavioral measures. Sci. Data 2, 150031 (2015).

50. Marcus, D. S. et al. Human Connectome Project informatics: Quality control, database services, and data visualization. NeuroImage 80, 202–219 (2013).

51. Khosla, M., Jamison, K., Kuceyeski, A. & Sabuncu, M. R. Ensemble learning with 3D convolutional neural networks for functional connectome-based prediction. NeuroImage 199, 651–662 (2019).

52. Smith, S. M. et al. Resting-state fMRI in the Human Connectome Project. NeuroImage 80, 144–168 (2013).

53. Ronneberger, O., Fischer, P. & Brox, T. U-Net: Convolutional Networks for Biomedical Image Segmentation. in Medical Image Computing and Computer-Assisted Intervention – MICCAI 2015 (eds. Navab, N., Hornegger, J., Wells, W. M. & Frangi, A. F.) vol. 9351 234–241 (Springer International Publishing, Cham, 2015).

54. Oktay, O., et al. Attention U-Net: Learning Where to Look for the Pancreas. Preprint at 10.48550/ARXIV.1804.03999 (2018).

55. Smith, S. M. et al. A positive-negative mode of population covariation links brain connectivity, demographics and behavior. Nat. Neurosci. 18, 1565–1567 (2015).

56. He, T. Deep neural networks and kernel regression achieve comparable accuracies for functional connectivity prediction of behavior and demographics. 15 (2020).

