## Supplementary Material for "Generating Synthetic Task-based Brain Fingerprints for Population Neuroscience Using Deep Learning"

#### Preprocessing

***Human Connectome Project Development (HCP-D):*** Similarly, the minimally preprocessed volumetric resting state (e.g., rfMRI_REST_hp0_clean.nii.gz) and task-fMRI (e.g., tfMRI_GUESSING_PA_hp0_clean.nii.gz) images provided by the HCP were used in our study. The voxel-to-ROI matrices for each subject within the dataset were computed using the same ICA-based parcellation used in the training dataset. The task-based contrast maps were calculated using "nilearn.glm.first_level.FirstLevelModel" method from Nilearn[1]. Specifically, the first level model with a high-pass filter (0.01 Hz), "glover" hemodynamic response function, smoothing with 6 mm FWHM Gaussian kernel, and z-transformation was applied to minimally preprocessed task-based images. Nuisance correction was done using the Friston 24 motion parameters.

***UK Biobank (UKB):*** We used the consortium-processed dataset, which was preprocessed using an FSL-based pipeline, which includes unwarping, motion correction, fieldmap correction, registration, normalization, automatic noise selection as implemented in ICA-AROMA[2], and smoothing using a 5 mm FWHM Gaussian kernel. Following preprocessing, the same ICA-based parcellation used in the previous analyses was utilized to extract general-purpose resting-state images and compute voxel-to-ROI time series. Z-transformed contrast maps for the Emotion task were computed using FSL. Subjects with a mean relative root mean square head motion greater than 0.5 mm were excluded.

#### Additional Performance Metrics

***Dice AUC Score****:* The Dice coefficient[3] (also known as the F1 score) is a widely used metric in medical image segmentation that measures the voxel-wise overlap between predicted and actual task-based contrast activation patterns (i.e., thresholded contrast maps), thereby capturing the spatial distribution similarity between images. To compute the Dice coefficient, both predicted and actual contrast maps were thresholded with a series of activity thresholds to generate binary maps of activated voxels. For each threshold $t$, the Dice coefficient is calculated as:

$$Dice_t = \frac{2|Predicted\ Map_t \cap Actual\ Map_t|}{|Predicted\ Map_t| + |Actual\ Map_t|}$$

where $|Predicted\ Map_t|$ and $|Actual\ Map_t|$ are the number of activated voxels in the predicted and actual contrast maps, respectively, and $|Predicted\ Map_t \cap Actual\ Map_t|$ denotes the overlap between them. To account for variability in activation thresholds, Dice scores over a range of thresholds (5% to 50% with 5% interval, indicating the most activated voxels) were integrated to create Area Under the Dice Curve (Dice AUC)[4].

***Fingerprinting Score****:* Fingerprinting score, introduced by Finn et al.[5], is another metric used to evaluate a model's ability to predict subject-specific contrast maps. It is computed by correlating each subject's predicted task contrast with both their actual task contrast and the actual task contrasts of all other subjects. The score represents the fraction of subjects whose predicted contrast achieves the highest correlation with their own actual contrast (i.e., argmax), quantifying individual predictability. Similar to the discriminability score, we normalize the diagonality index by dividing it by reconstruction performance. A discriminability score of 1 indicates perfect similarity to a subject's own actual contrast map and perfect separation from other subjects, suggesting that synthetic maps capture individual-specific characteristics beyond average information.

| Input Layer # | Layer # | Layer Type | Normalization | Activation | Output Shape |
|---|---|---|---|---|---|
| - | 1 | Input Data | - | - | 50 x 76 x 93 x 78 |
| 1 | 2 | Conv3D | BatchNorm3D | ReLU | 64 x 76 x 93 x 78 |
| 2 | 3 | MaxPool3D | - | - | 64 x 38 x 46 x 39 |
| 3 | 4 | Conv3D | BatchNorm3D | ReLU | 128 x 38 x 46 x 39 |
| 4 | 5 | MaxPool3D | - | - | 128 x 19 x 23 x 19 |
| 5 | 6 | Conv3D | BatchNorm3D | ReLU | 256 x 19 x 23 x 19 |
| 6 | 7 | ConvTranspose3D (Upsample) | - | - | 128 x 38 x 46 x 38 |
| 4 | 8 | Attention + Concat | - | - | 256 x 38 x 46 x 39 |
| 8 | 9 | Conv3D | BatchNorm3D | ReLU | 128 x 38 x 46 x 39 |
| 9 | 10 | ConvTranspose3D (Upsample) | - | - | 64 x 76 x 92 x 78 |
| 2 | 11 | Attention + Concat | - | - | 128 x 76 x 93 x 78 |
| 11 | 12 | Conv3D | BatchNorm3D | ReLU | 64 x 76 x 93 x 78 |
| 12 | 13 | Conv3D | - | - | 47 x 76 x 93 x 78 |

**Supplementary Table 1:** DeepTaskGen Architecture. The table is divided into individual convolutional blocks by horizontal lines. Layers 2-6 form the encoding block, which includes the bottleneck, while layers 6-12 constitute the decoding block. Layer 13 consists of a Conv3D with a kernel size of 1. The parameters for convolutional layers from the encoding and decoding blocks are as follows: kernel size = 3, padding = 1, stride = 1. Skip connections from the encoding blocks are filtered through attention gates before being concatenated with their corresponding decoding blocks. During this concatenation, trilinear interpolation is applied to address any slight dimensional discrepancies between the corresponding encoding and decoding layers. Total number of trainable parameters is 3,447,875.

| Site | Sample Size | Sex (F) | Age (μ(σ)) | Task | TR/TE (ms) | Flip Angle | Multiband factor | Thickness (mm) | Frames |
|---|---|---|---|---|---|---|---|---|---|
| HCP Young Adult | 958 | 504 (52,6%) | 28.66 ($\sigma$ = 3.71) | Resting State | 720/33.01 ms | 52° | 8 | 2.0 mm | 4800 |
| | | | | Working Memory | | | | | 405 |
| | | | | Motor | | | | | 284 |
| | | | | Language | | | | | 316 |
| | | | | Social Cognition | | | | | 274 |
| | | | | Relational | | | | | 232 |
| | | | | Emotion Processing | | | | | 176 |
| | | | | Gambling | | | | | 253 |
| HCP Development | 637 | 343 (53,7%) | 14.49 ($\sigma$ = 4.05) | Resting State | 800/37 ms | 52° | 8 | 2.0 mm | 976 |
| | | | | Guessing | | | | | 280* |
| | | | | Emotion Processing | | | | | 178 |
| UK Biobank | 20,792 | 11,214 | 60.82 | Resting State | 735/39 ms | 52° | 8 | 2.4 mm | 490 |

| | | | | | | | | | |
|---|---|---|---|---|---|---|---|---|---|
| | | (53,9%) | ($\sigma$ = 7.45) | Emotion Processing | | | | | 332 |

**Supplementary Table 2:** Sample details and functional scan acquisition parameters.
*Only Run 1 PA was used to match the Emotion Processing Task.

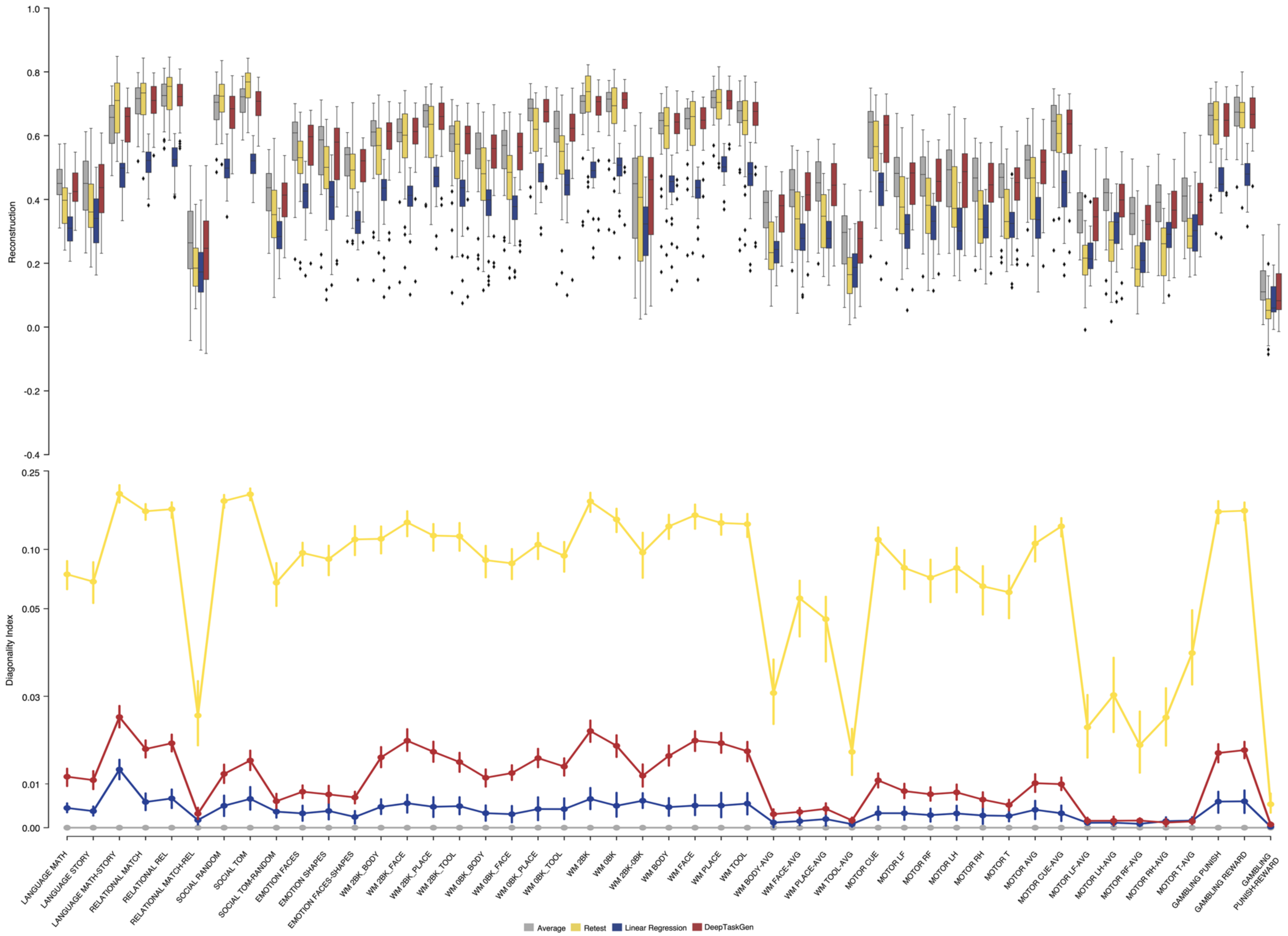
Reconstruction
1.0
0.8
0.6
0.4
0.2
0.0
-0.2
-0.4
Diagonality Index
0.25
0.10
0.05
0.03
0.01
0.00
LANGUAGE MATH
LANGUAGE STORY
LANGUAGE MATH-STORY
RELATIONAL MATCH
RELATIONAL REL
RELATIONAL MATCH-REL
SOCIAL RANDOM
SOCIAL TOM
SOCIAL TOM-RANDOM
EMOTION FACES
EMOTION SHAPES
EMOTION FACES-SHAPES
WM 2BK_BODY
WM 2BK_FACE
WM 2BK_PLACE
WM 2BK_TOOL
WM 0BK_BODY
WM 0BK_FACE
WM 0BK_PLACE
WM 0BK_TOOL
WM 2BK
WM 0BK
WM 2BK-0BK
WM BODY
WM FACE
WM PLACE
WM TOOL
WM BODY-AVG
WM FACE-AVG
WM PLACE-AVG
WM TOOL-AVG
MOTOR CUE
MOTOR LF
MOTOR RF
MOTOR LH
MOTOR RH
MOTOR T
MOTOR AVG
MOTOR CUE-AVG
MOTOR LF-AVG
MOTOR LH-AVG
MOTOR RF-AVG
MOTOR RH-AVG
MOTOR T-AVG
GAMBLING PUNISH
GAMBLING REWARD
GAMBLING PUNISH-REWARD
Average
Retest
Linear Regression
DeepTaskGen

**Supplementary Figure 1.** Reconstruction performance computed by taking Pearson's correlation between predicted and actual contrast maps and the diagonality index (the difference between the on-diagonal and the mean off-diagonal elements in a correlation matrix, normalized by the mean on-diagonal values) of DeepTaskGen and various baselines for 47 task contrasts from HCP-YA.

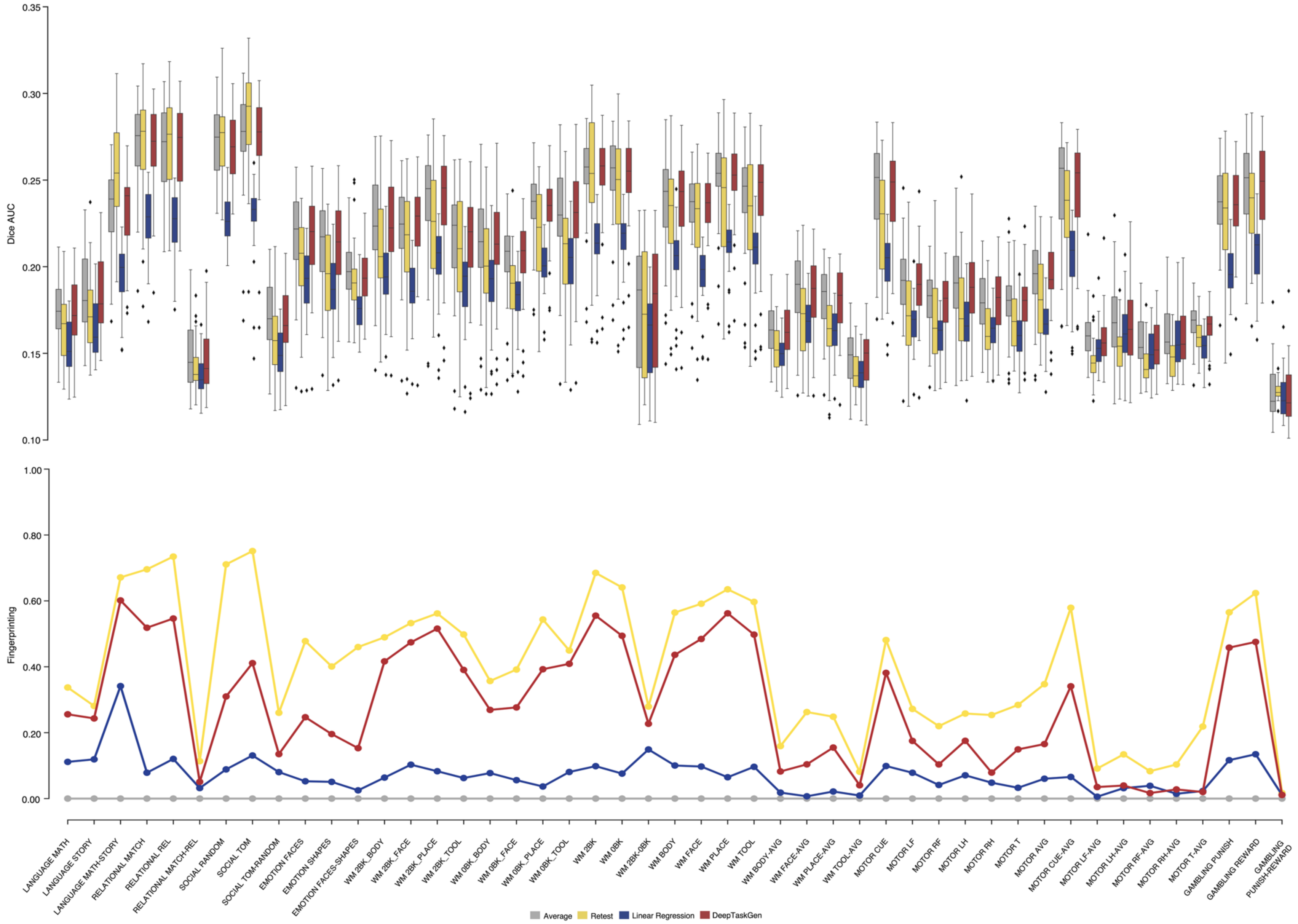

Dice AUC
0.35
0.30
0.25
0.20
0.15
0.10
Fingerprinting
1.00
0.80
0.60
0.40
0.20
0.00
LANGUAGE MATH
LANGUAGE STORY
LANGUAGE MATH-STORY
RELATIONAL MATCH
RELATIONAL REL
RELATIONAL MATCH-REL
SOCIAL RANDOM
SOCIAL TOM
SOCIAL TOM-RANDOM
EMOTION FACES
EMOTION SHAPES
EMOTION FACES-SHAPES
WM 2BK_BODY
WM 2BK_FACE
WM 2BK_PLACE
WM 2BK_TOOL
WM 0BK_BODY
WM 0BK_FACE
WM 0BK_PLACE
WM 0BK_TOOL
WM 2BK
WM 0BK
WM 2BK-0BK
WM BODY
WM FACE
WM PLACE
WM TOOL
WM BODY-AVG
WM FACE-AVG
WM PLACE-AVG
WM TOOL-AVG
MOTOR CUE
MOTOR LF
MOTOR RF
MOTOR LH
MOTOR RH
MOTOR T
MOTOR AVG
MOTOR CUE-AVG
MOTOR LF-AVG
MOTOR LH-AVG
MOTOR RF-AVG
MOTOR RH-AVG
MOTOR T-AVG
GAMBLING PUNISH
GAMBLING REWARD
GAMBLING PUNISH-REWARD
Average
Retest
Linear Regression
DeepTaskGen

**Supplementary Figure 2.** Dice AUC and fingerprinting score of DeepTaskGen and various baselines for 47 task contrasts from HCP-YA.

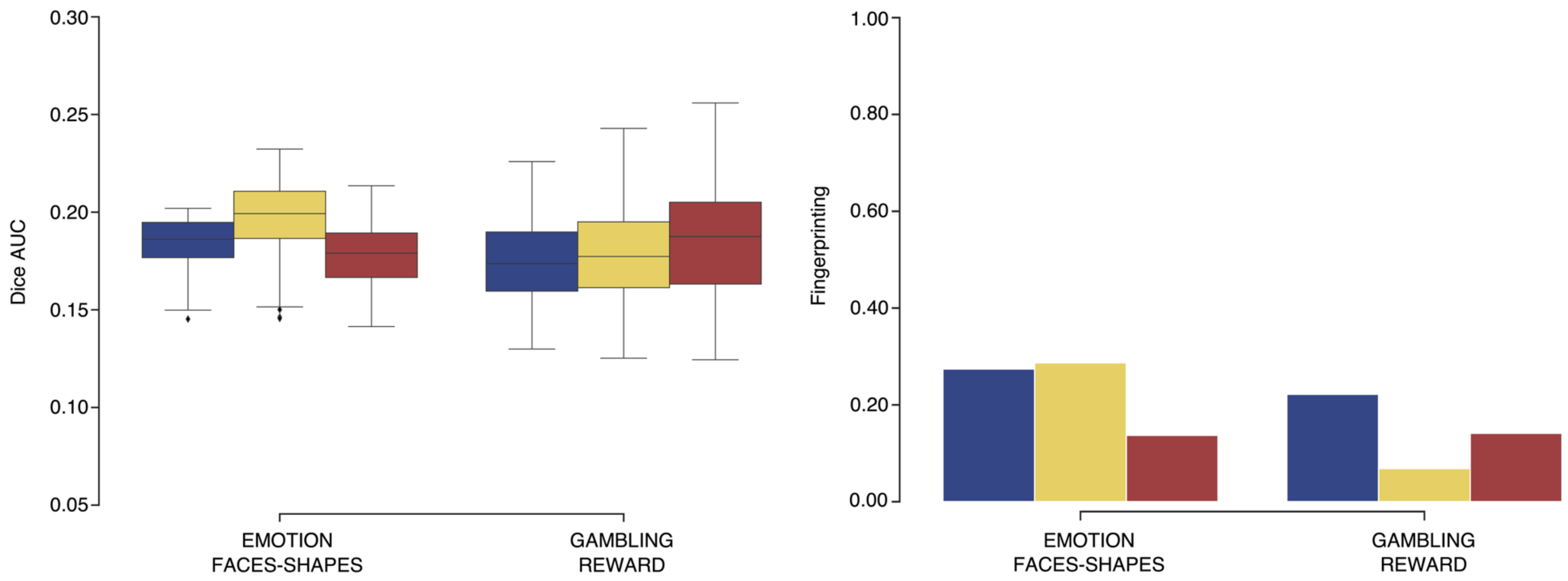


**Supplementary Figure 3.** Dice AUC and fingerprinting scores for fine-tuned and non-fine-tuned DeepTaskGen models, as well as the baseline linear model, for the EMOTION FACES-SHAPES and GAMBLING REWARD task contrasts from HCP-D.

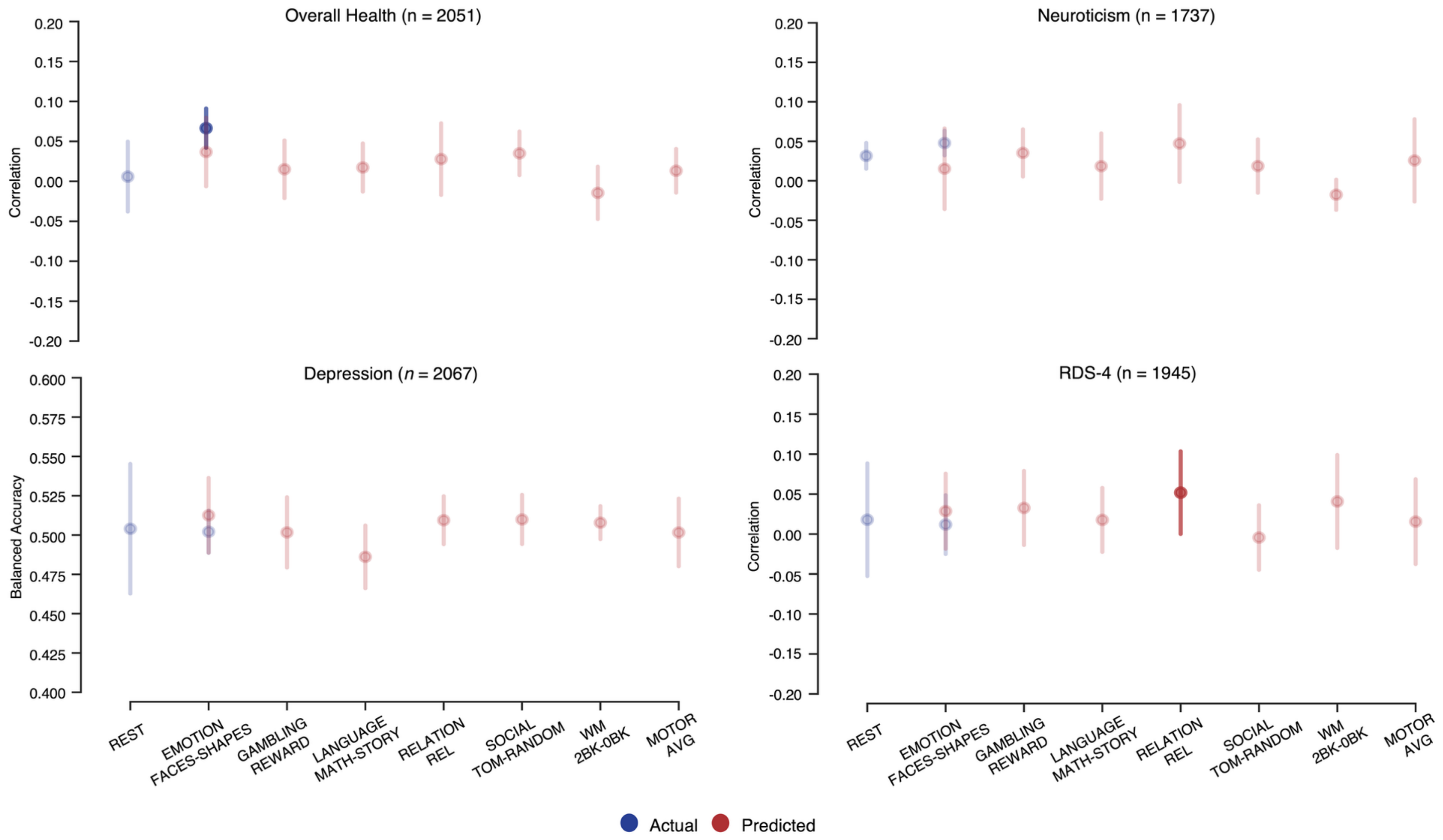


**Supplementary Figure 4.** Additional predictions of subjects' overall health, neuroticism, depression, and Recent Depression Symptoms-4 (RDS-4) scores using task contrast maps and resting-state connectome on UK Biobank. Actual and synthetic brain measures are depicted in blue and red colors, respectively. Significant predictions based on permutation testing are highlighted. Error bars indicate the standard deviation of prediction performance across five CV folds. Balanced accuracy was used to measure depression classification performance, while Pearson's correlation was employed to assess other variables. Sample sizes for all analyses are indicated in each figure.

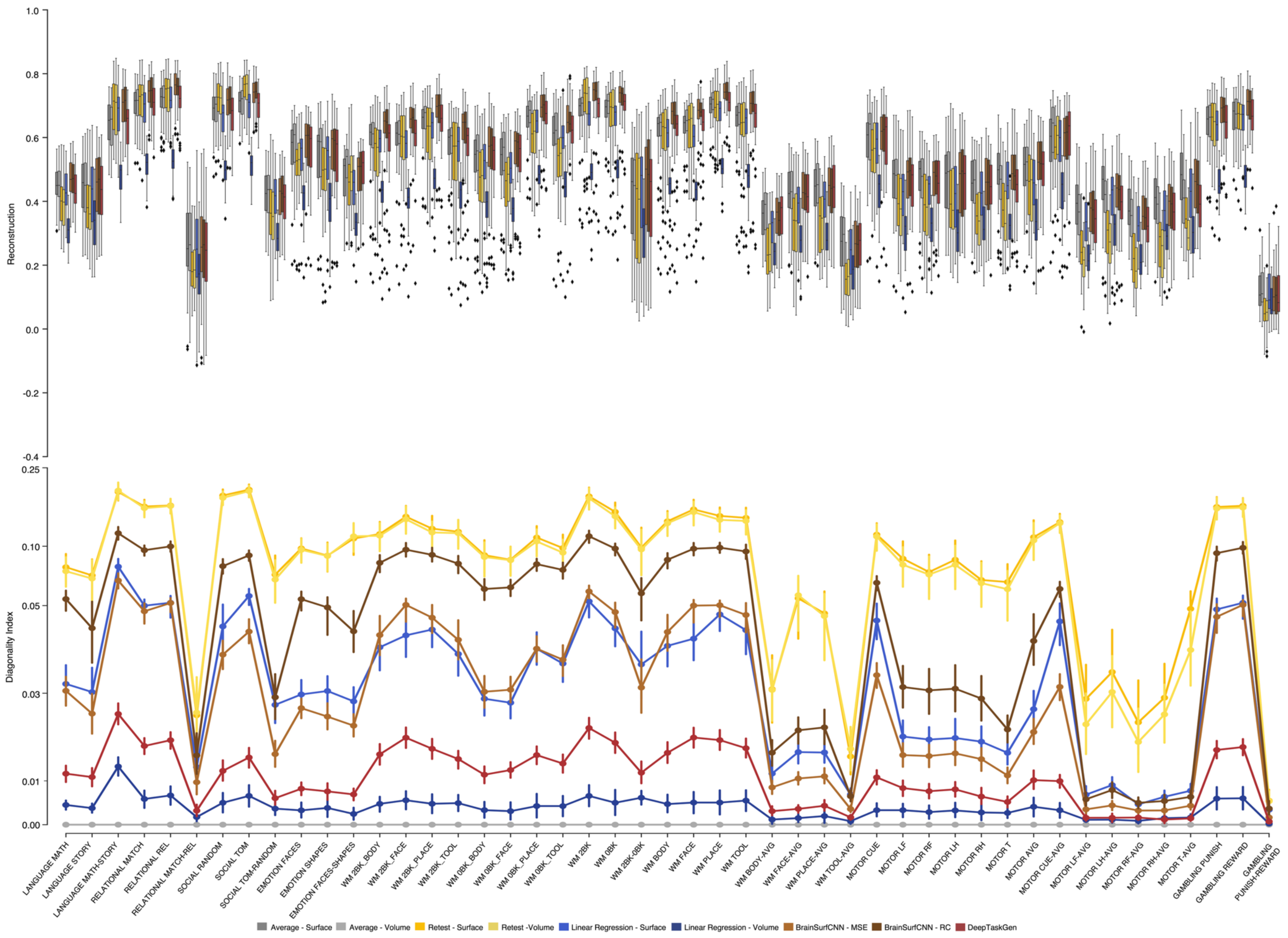
Reconstruction
Diagonality Index
LANGUAGE MATH
LANGUAGE STORY
LANGUAGE MATH-STORY
RELATIONAL MATCH
RELATIONAL REL
RELATIONAL MATCH-REL
SOCIAL RANDOM
SOCIAL TOM
SOCIAL TOM-RANDOM
EMOTION FACES
EMOTION SHAPES
EMOTION FACES-SHAPES
WM 2BK_BODY
WM 2BK_FACE
WM 2BK_PLACE
WM 2BK_TOOL
WM 0BK_BODY
WM 0BK_FACE
WM 0BK_PLACE
WM 0BK_TOOL
WM 2BK
WM 0BK
WM 2BK-0BK
WM BODY
WM FACE
WM PLACE
WM TOOL
WM BODY-AVG
WM FACE-AVG
WM PLACE-AVG
WM TOOL-AVG
MOTOR CUE
MOTOR LF
MOTOR RF
MOTOR LH
MOTOR RH
MOTOR T
MOTOR AVG
MOTOR CUE-AVG
MOTOR LF-AVG
MOTOR LH-AVG
MOTOR RF-AVG
MOTOR RH-AVG
MOTOR T-AVG
GAMBLING PUNISH
GAMBLING REWARD
GAMBLING PUNISH-REWARD
Average - Surface
Average - Volume
Retest - Surface
Retest -Volume
Linear Regression - Surface
Linear Regression - Volume
BrainSurfCNN - MSE
BrainSurfCNN - RC
DeepTaskGen

**Supplementary Figure 5.** Reconstruction performance and diagonality index of surface- and volumetric-based methods and baselines for 47 task contrasts from HCP-YA. Average: Group Average; Retest: Test-Retest subjects $(n = 39)$; Linear Regression: Tavor et al., 2016[6]; BrainSurfCNN - MSE: 50 epochs with MSE loss; BrainSurfCNN - RC: Fine-tuned model with 50 epochs using reconstructive-contrastive loss[4]; DeepTaskGen - proposed method.

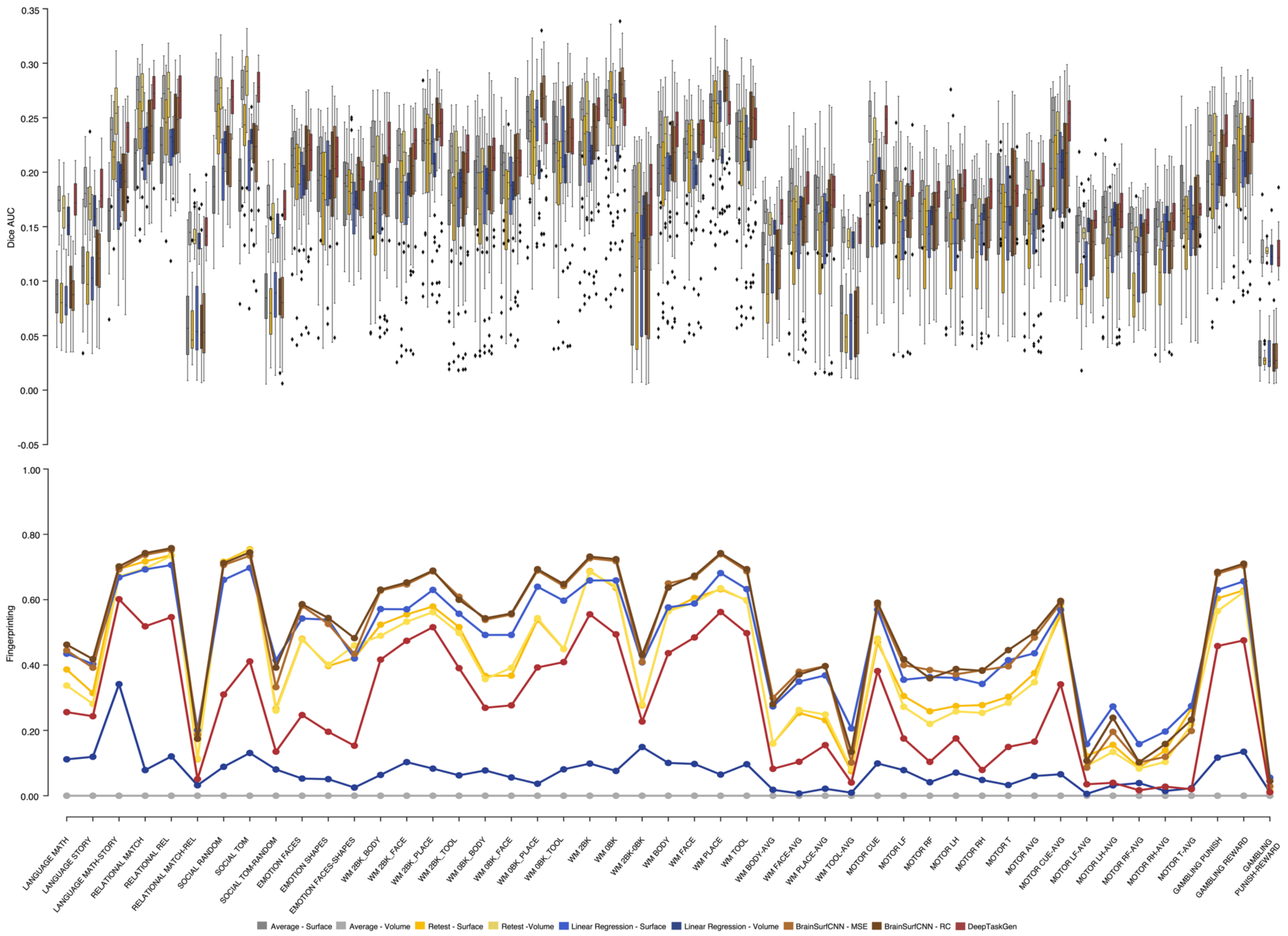
Dice AUC
Fingerprinting
LANGUAGE MATH
LANGUAGE STORY
LANGUAGE MATH-STORY
RELATIONAL MATCH
RELATIONAL REL
RELATIONAL MATCH-REL
SOCIAL RANDOM
SOCIAL TOM
SOCIAL TOM-RANDOM
EMOTION FACES
EMOTION SHAPES
EMOTION FACES-SHAPES
WM 2BK_BODY
WM 2BK_FACE
WM 2BK_PLACE
WM 2BK_TOOL
WM 0BK_BODY
WM 0BK_FACE
WM 0BK_PLACE
WM 0BK_TOOL
WM 2BK
WM 0BK
WM 2BK-0BK
WM BODY
WM FACE
WM PLACE
WM TOOL
WM BODY-AVG
WM FACE-AVG
WM PLACE-AVG
WM TOOL-AVG
MOTOR CUE
MOTOR LF
MOTOR RF
MOTOR LH
MOTOR RH
MOTOR T
MOTOR AVG
MOTOR CUE-AVG
MOTOR LF-AVG
MOTOR LH-AVG
MOTOR RF-AVG
MOTOR RH-AVG
MOTOR T-AVG
GAMBLING PUNISH
GAMBLING REWARD
GAMBLING PUNISH-REWARD
Average - Surface
Average - Volume
Retest - Surface
Retest -Volume
Linear Regression - Surface
Linear Regression - Volume
BrainSurfCNN - MSE
BrainSurfCNN - RC
DeepTaskGen

**Supplementary Figure 6.** Dice AUC and fingerprinting scores of surface- and volumetric-based methods and baselines for 47 task contrasts from HCP-YA. Average: Group Average; Retest: Test-Retest subjects $(n = 39)$; Linear Regression: Tavor et al., 2016[6]; BrainSurfCNN - MSE: 50 epochs with MSE loss; BrainSurfCNN - RC: Fine-tuned model with 50 epochs using reconstructive-contrastive loss[4]; DeepTaskGen - proposed method.

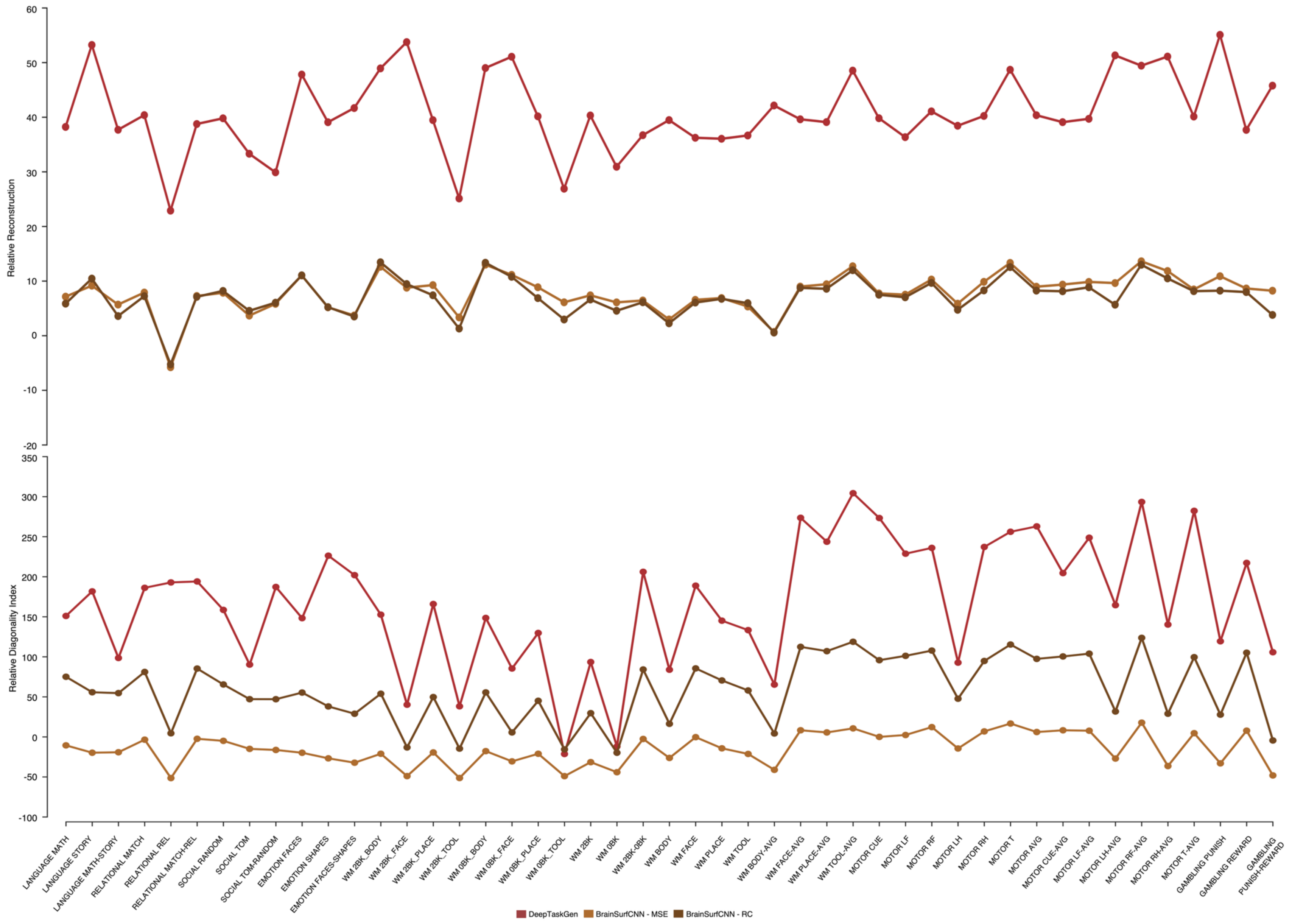
Relative Reconstruction
60
50
40
30
20
10
0
-10
-20
Relative Diagonality Index
350
300
250
200
150
100
50
0
-50
-100
LANGUAGE MATH
LANGUAGE STORY
LANGUAGE MATH-STORY
RELATIONAL MATCH
RELATIONAL REL
RELATIONAL MATCH-REL
SOCIAL RANDOM
SOCIAL TOM
SOCIAL TOM-RANDOM
EMOTION FACES
EMOTION SHAPES
EMOTION FACES-SHAPES
WM 2BK_BODY
WM 2BK_FACE
WM 2BK_PLACE
WM 2BK_TOOL
WM 0BK_BODY
WM 0BK_FACE
WM 0BK_PLACE
WM 0BK_TOOL
WM 2BK
WM 0BK
WM 2BK-0BK
WM BODY
WM FACE
WM PLACE
WM TOOL
WM BODY-AVG
WM FACE-AVG
WM PLACE-AVG
WM TOOL-AVG
MOTOR CUE
MOTOR LF
MOTOR RF
MOTOR LH
MOTOR RH
MOTOR T
MOTOR AVG
MOTOR CUE-AVG
MOTOR LF-AVG
MOTOR LH-AVG
MOTOR RF-AVG
MOTOR RH-AVG
MOTOR T-AVG
GAMBLING PUNISH
GAMBLING REWARD
GAMBLING PUNISH-REWARD
DeepTaskGen
BrainSurfCNN - MSE
BrainSurfCNN - RC

**Supplementary Figure 7.** Relative reconstruction performance and diagonality index scores of BrainSurfCNN and DeepTaskGen for 47 task contrasts from HCP-YA. Relative scores are computed against the corresponding baseline linear regression[6] model's performance, indicating the models' performance gain or loss compared to the baseline linear model (i.e., $Relative\ Performance = \frac{Model\ Performance - Baseline\ Performance}{Baseline\ Performance} * 100$). Positive values indicate a performance gain over a linear regression model, while negative values indicate a loss. DeepTaskGen: proposed method; BrainSurfCNN - MSE: 50 epochs with MSE loss; BrainSurfCNN - RC: Fine-tuned model with 50 epochs using reconstructive-contrastive loss[4]. The raw performance of the models is indicated in Supplementary Figure 5. DeepTaskGen provides a greater performance gain over the linear model compared to the surface-based BrainSurfCNN.Despite varying degrees, all models exhibit a similar pattern of gains and losses compared to the linear model across 47 task contrasts (i.e., similar dips and peaks), indicating variability across the task contrast maps.

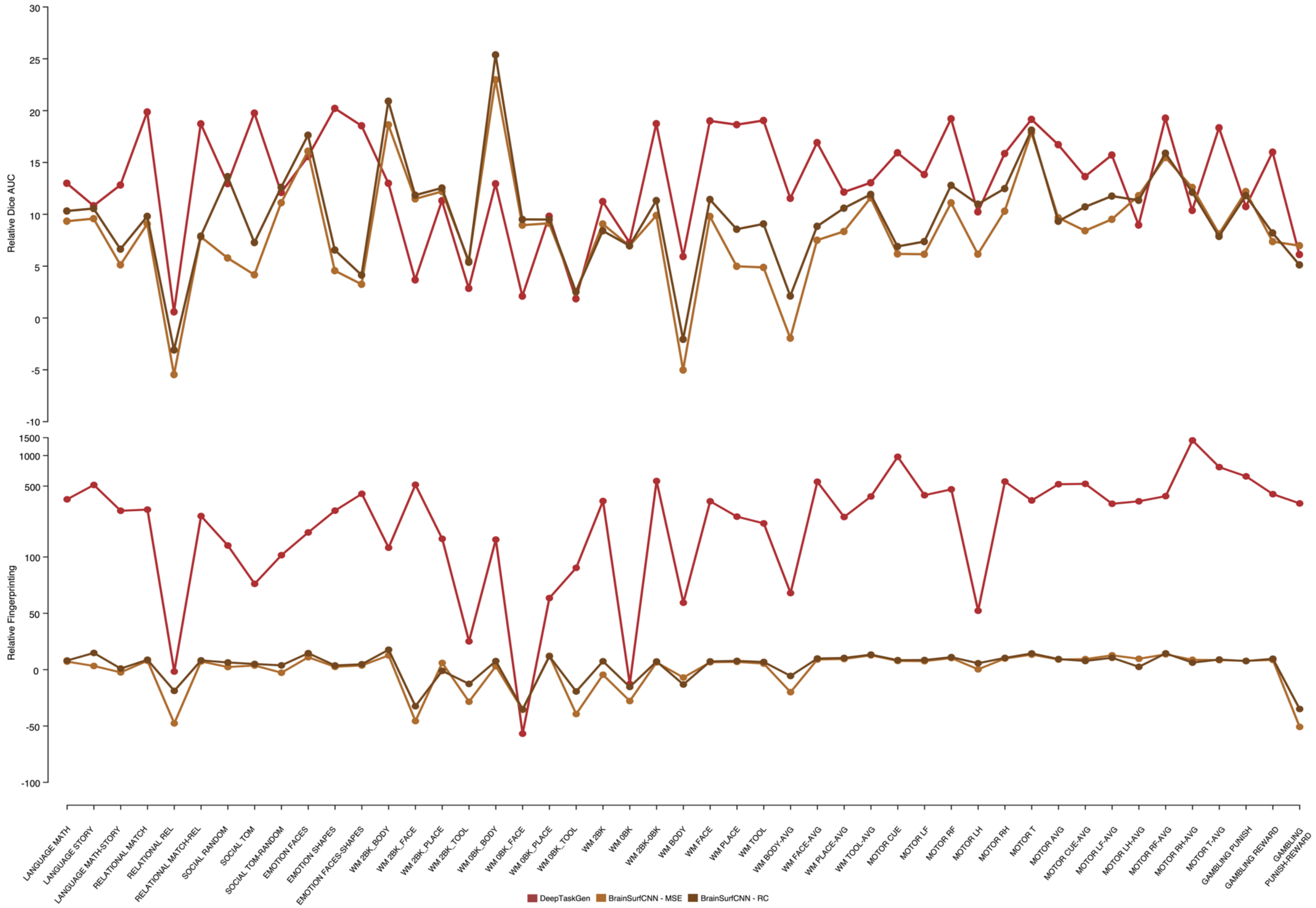
Relative Dice AUC
30
25
20
15
10
5
0
-5
-10
Relative Fingerprinting
1500
1000
500
100
50
0
-50
-100
LANGUAGE MATH
LANGUAGE STORY
LANGUAGE MATH-STORY
RELATIONAL MATCH
RELATIONAL REL
RELATIONAL MATCH-REL
SOCIAL RANDOM
SOCIAL TOM
SOCIAL TOM-RANDOM
EMOTION FACES
EMOTION SHAPES
EMOTION FACES-SHAPES
WM 2BK_BODY
WM 2BK_FACE
WM 2BK_PLACE
WM 2BK_TOOL
WM 0BK_BODY
WM 0BK_FACE
WM 0BK_PLACE
WM 0BK_TOOL
WM 2BK
WM 0BK
WM 2BK-0BK
WM BODY
WM FACE
WM PLACE
WM TOOL
WM BODY-AVG
WM FACE-AVG
WM PLACE-AVG
WM TOOL-AVG
MOTOR CUE
MOTOR LF
MOTOR RF
MOTOR LH
MOTOR RH
MOTOR T
MOTOR AVG
MOTOR CUE-AVG
MOTOR LF-AVG
MOTOR LH-AVG
MOTOR RF-AVG
MOTOR RH-AVG
MOTOR T-AVG
GAMBLING PUNISH
GAMBLING REWARD
GAMBLING PUNISH-REWARD
DeepTaskGen
BrainSurfCNN - MSE
BrainSurfCNN - RC

**Supplementary Figure 8.** Relative Dice AUC and fingerprinting scores of BrainSurfCNN and DeepTaskGen for 47 task contrasts from HCP-YA. Relative scores are computed against the corresponding baseline linear regression[6] model's performance, indicating the models' performance gain or loss compared to the baseline linear model (i.e., $Relative\ Performance = \frac{Model\ Performance - Baseline\ Performance}{Baseline\ Performance} * 100$. Positive values indicate a performance gain over a linear regression model, while negative values indicate a loss. DeepTaskGen: proposed method; BrainSurfCNN - MSE: 50 epochs with MSE loss; BrainSurfCNN - RC: Fine-tuned model with 50 epochs using reconstructive-contrastive loss[4]. The raw performance of the models is indicated in Supplementary Figure 6. DeepTaskGen provides a greater performance gain over the linear model compared to the surface-based BrainSurfCNN.Despite varying degrees, all models exhibit a similar pattern of gains and losses compared to the linear model across 47 task contrasts (i.e., similar dips and peaks), indicating variability across the task contrast maps.

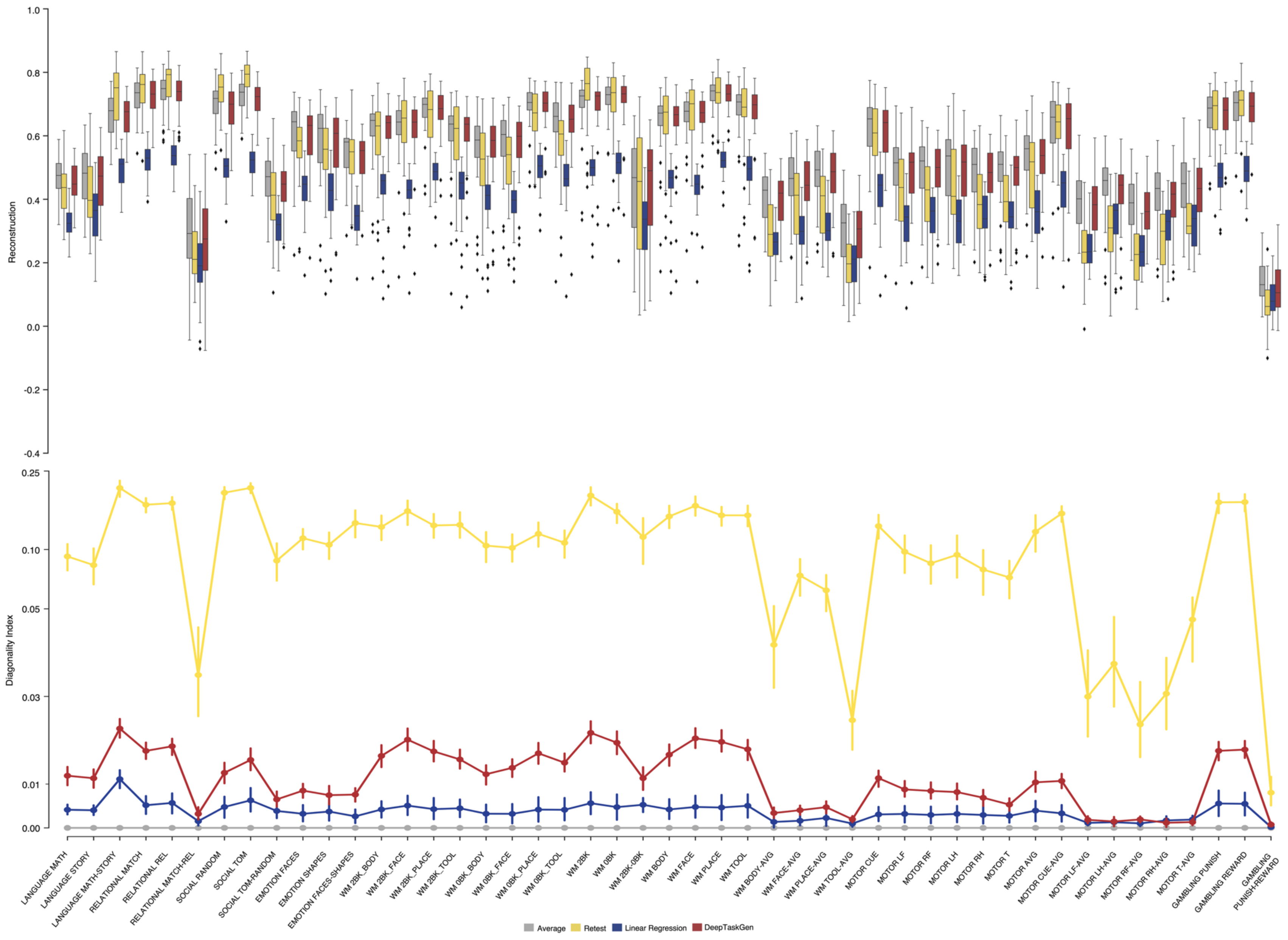
Reconstruction
1.0
0.8
0.6
0.4
0.2
0.0
-0.2
-0.4
Diagonality Index
0.25
0.10
0.05
0.03
0.01
0.00
LANGUAGE MATH
LANGUAGE STORY
LANGUAGE MATH-STORY
RELATIONAL MATCH
RELATIONAL REL
RELATIONAL MATCH-REL
SOCIAL RANDOM
SOCIAL TOM
SOCIAL TOM-RANDOM
EMOTION FACES
EMOTION SHAPES
EMOTION FACES-SHAPES
WM 2BK_BODY
WM 2BK_FACE
WM 2BK_PLACE
WM 2BK_TOOL
WM 0BK_BODY
WM 0BK_FACE
WM 0BK_PLACE
WM 0BK_TOOL
WM 2BK
WM 0BK
WM 2BK-0BK
WM BODY
WM FACE
WM PLACE
WM TOOL
WM BODY-AVG
WM FACE-AVG
WM PLACE-AVG
WM TOOL-AVG
MOTOR CUE
MOTOR LF
MOTOR RF
MOTOR LH
MOTOR RH
MOTOR T
MOTOR AVG
MOTOR CUE-AVG
MOTOR LF-AVG
MOTOR LH-AVG
MOTOR RF-AVG
MOTOR RH-AVG
MOTOR T-AVG
GAMBLING PUNISH
GAMBLING REWARD
GAMBLING PUNISH-REWARD
Average
Retest
Linear Regression
DeepTaskGen

**Supplementary Figure 9.** Reconstruction performance and the diagonality index of DeepTaskGen and various baselines for 47 cortical task contrasts from HCP-YA. Subcortical areas were masked out during computation of the performance metrics.

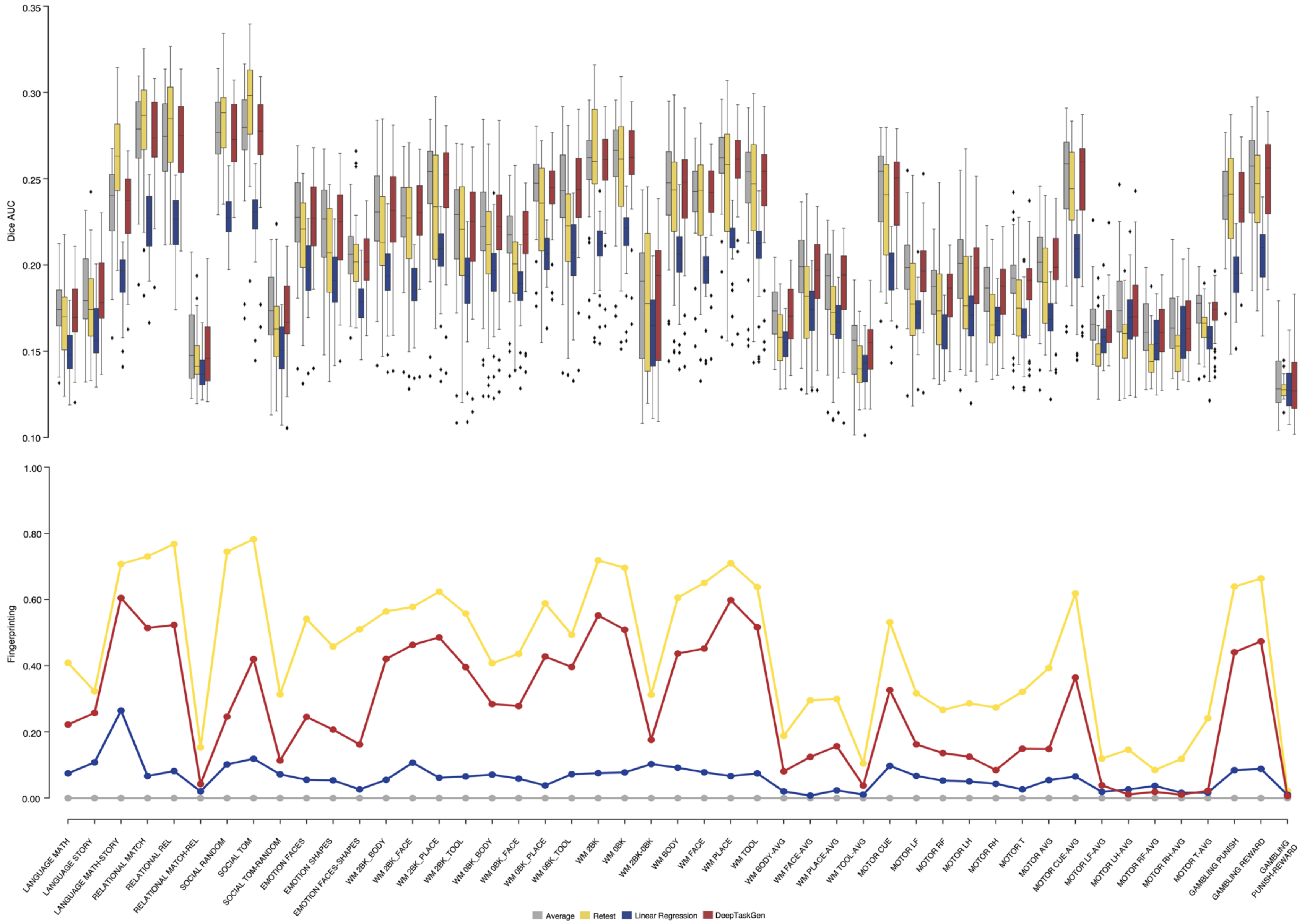
Dice AUC
0.35
0.30
0.25
0.20
0.15
0.10
Fingerprinting
1.00
0.80
0.60
0.40
0.20
0.00
LANGUAGE MATH
LANGUAGE STORY
LANGUAGE MATH-STORY
RELATIONAL MATCH
RELATIONAL REL
RELATIONAL MATCH-REL
SOCIAL RANDOM
SOCIAL TOM
SOCIAL TOM-RANDOM
EMOTION FACES
EMOTION SHAPES
EMOTION FACES-SHAPES
WM 2BK_BODY
WM 2BK_FACE
WM 2BK_PLACE
WM 2BK_TOOL
WM 0BK_BODY
WM 0BK_FACE
WM 0BK_PLACE
WM 0BK_TOOL
WM 2BK
WM 0BK
WM 2BK-0BK
WM BODY
WM FACE
WM PLACE
WM TOOL
WM BODY-AVG
WM FACE-AVG
WM PLACE-AVG
WM TOOL-AVG
MOTOR CUE
MOTOR LF
MOTOR RF
MOTOR LH
MOTOR RH
MOTOR T
MOTOR AVG
MOTOR CUE-AVG
MOTOR LF-AVG
MOTOR LH-AVG
MOTOR RF-AVG
MOTOR RH-AVG
MOTOR T-AVG
GAMBLING PUNISH
GAMBLING REWARD
GAMBLING PUNISH-REWARD
Average
Retest
Linear Regression
DeepTaskGen

**Supplementary Figure 10.** Dice AUC and fingerprinting score of DeepTaskGen and various baselines for 47 cortical task contrasts from HCP-YA. Subcortical areas were masked out during computation of the performance metrics.

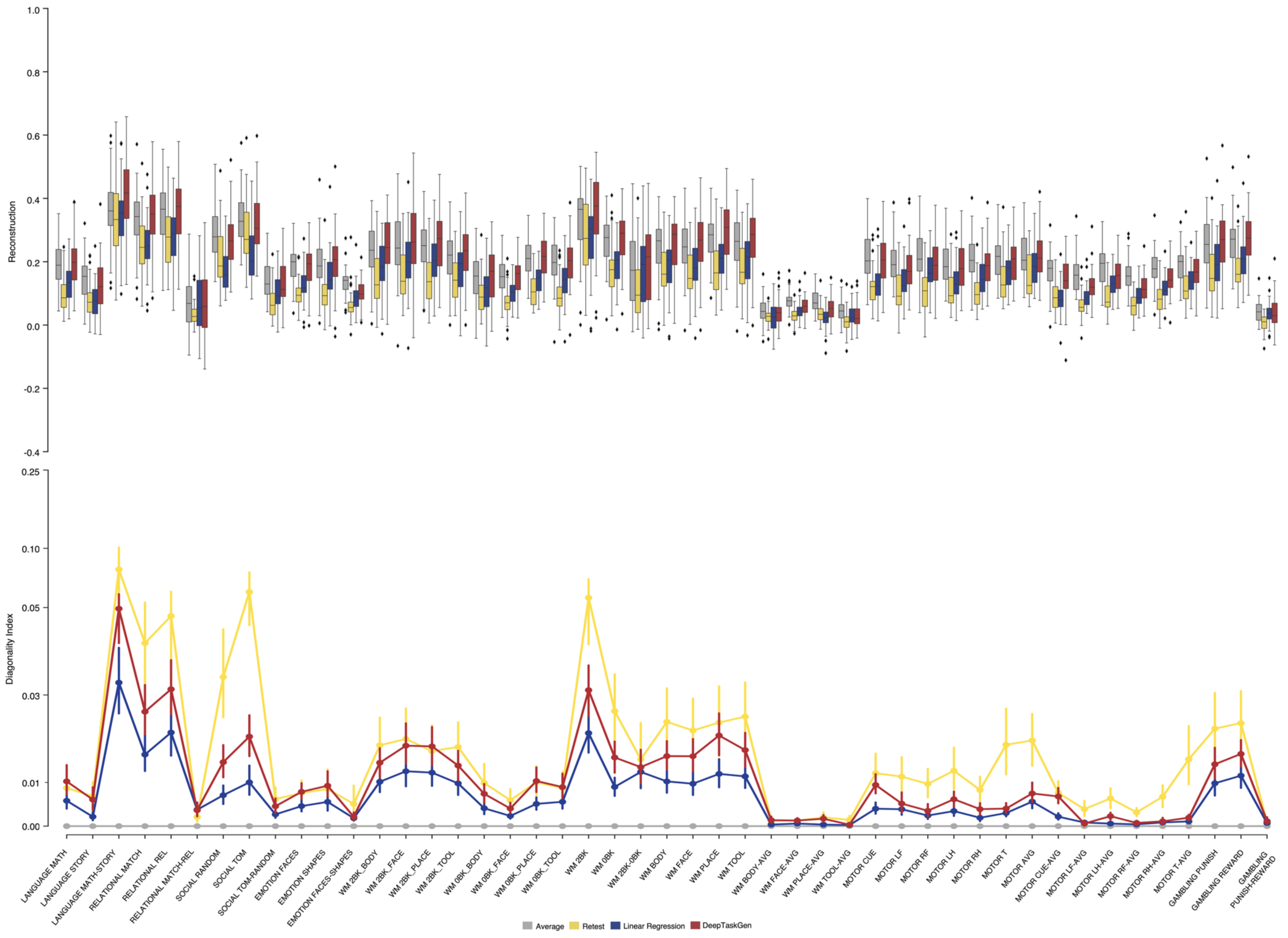
Reconstruction
1.0
0.8
0.6
0.4
0.2
0.0
-0.2
-0.4
Diagonality Index
0.25
0.10
0.05
0.03
0.01
0.00
LANGUAGE MATH
LANGUAGE STORY
LANGUAGE MATH-STORY
RELATIONAL MATCH
RELATIONAL REL
RELATIONAL MATCH-REL
SOCIAL RANDOM
SOCIAL TOM
SOCIAL TOM-RANDOM
EMOTION FACES
EMOTION SHAPES
EMOTION FACES-SHAPES
WM 2BK_BODY
WM 2BK_FACE
WM 2BK_PLACE
WM 2BK_TOOL
WM 0BK_BODY
WM 0BK_FACE
WM 0BK_PLACE
WM 0BK_TOOL
WM 2BK
WM 0BK
WM 2BK-0BK
WM BODY
WM FACE
WM PLACE
WM TOOL
WM BODY-AVG
WM FACE-AVG
WM PLACE-AVG
WM TOOL-AVG
MOTOR CUE
MOTOR LF
MOTOR RF
MOTOR LH
MOTOR RH
MOTOR T
MOTOR AVG
MOTOR CUE-AVG
MOTOR LF-AVG
MOTOR LH-AVG
MOTOR RF-AVG
MOTOR RH-AVG
MOTOR T-AVG
GAMBLING PUNISH
GAMBLING REWARD
GAMBLING PUNISH-REWARD
Average
Retest
Linear Regression
DeepTaskGen

**Supplementary Figure 11.** Reconstruction performance and the diagonality index of DeepTaskGen and various baselines for 47 subcortical task contrasts from HCP-YA. Cortical areas were masked out during computation of the performance metrics.

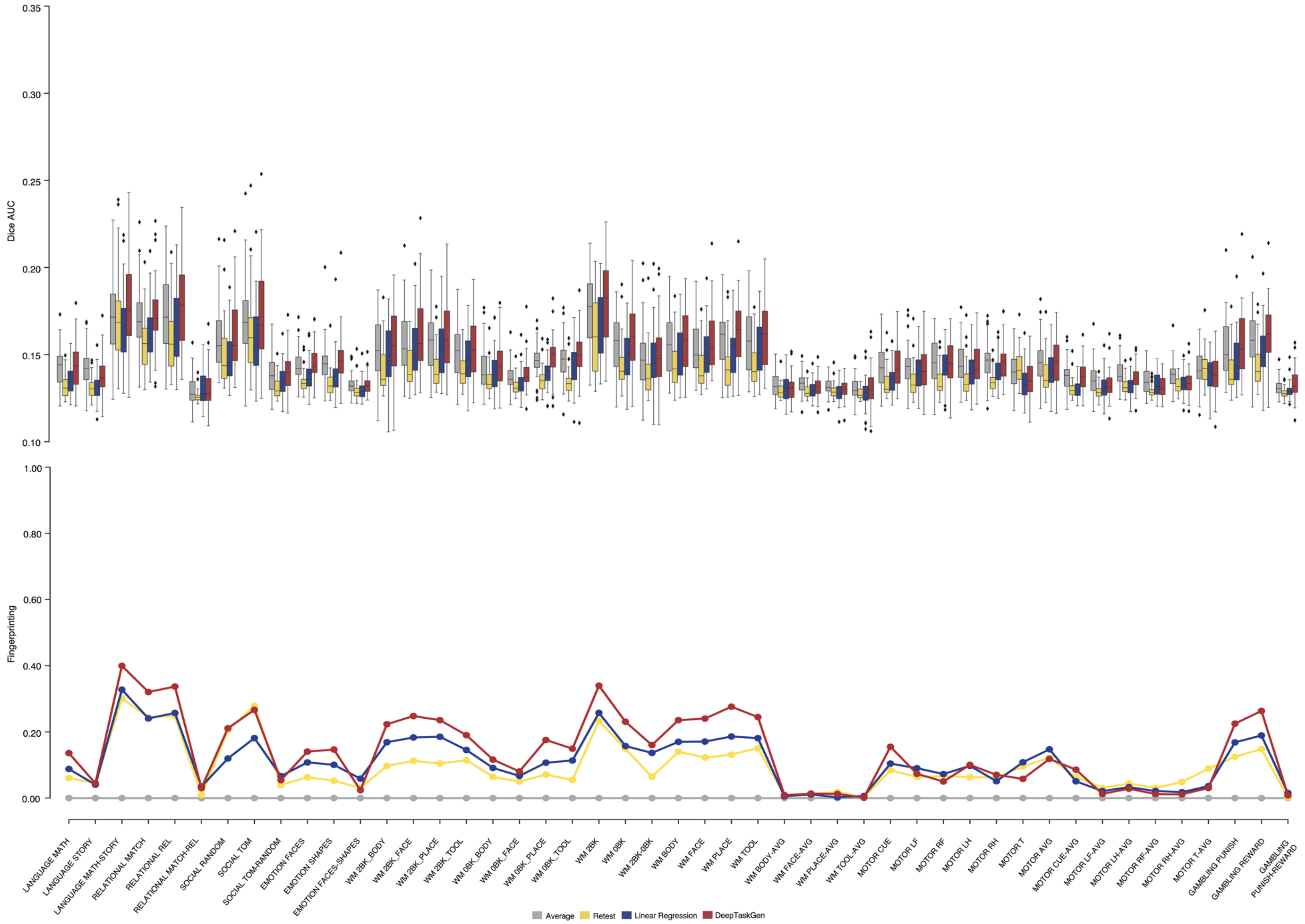
Dice AUC
0.35
0.30
0.25
0.20
0.15
0.10
Fingerprinting
1.00
0.80
0.60
0.40
0.20
0.00
LANGUAGE MATH
LANGUAGE STORY
LANGUAGE MATH-STORY
RELATIONAL MATCH
RELATIONAL REL
RELATIONAL MATCH-REL
SOCIAL RANDOM
SOCIAL TOM
SOCIAL TOM-RANDOM
EMOTION FACES
EMOTION SHAPES
EMOTION FACES-SHAPES
WM 2BK_BODY
WM 2BK_FACE
WM 2BK_PLACE
WM 2BK_TOOL
WM 0BK_BODY
WM 0BK_FACE
WM 0BK_PLACE
WM 0BK_TOOL
WM 2BK
WM 0BK
WM 2BK-0BK
WM BODY
WM FACE
WM PLACE
WM TOOL
WM BODY-AVG
WM FACE-AVG
WM PLACE-AVG
WM TOOL-AVG
MOTOR CUE
MOTOR LF
MOTOR RF
MOTOR LH
MOTOR RH
MOTOR T
MOTOR AVG
MOTOR CUE-AVG
MOTOR LF-AVG
MOTOR LH-AVG
MOTOR RF-AVG
MOTOR RH-AVG
MOTOR T-AVG
GAMBLING PUNISH
GAMBLING REWARD
GAMBLING PUNISH-REWARD
Average
Retest
Linear Regression
DeepTaskGen

**Supplementary Figure 12.** Dice AUC and fingerprinting score of DeepTaskGen and various baselines for 47 subcortical task contrasts from HCP-YA. Cortical areas were masked out during computation of the performance metrics.

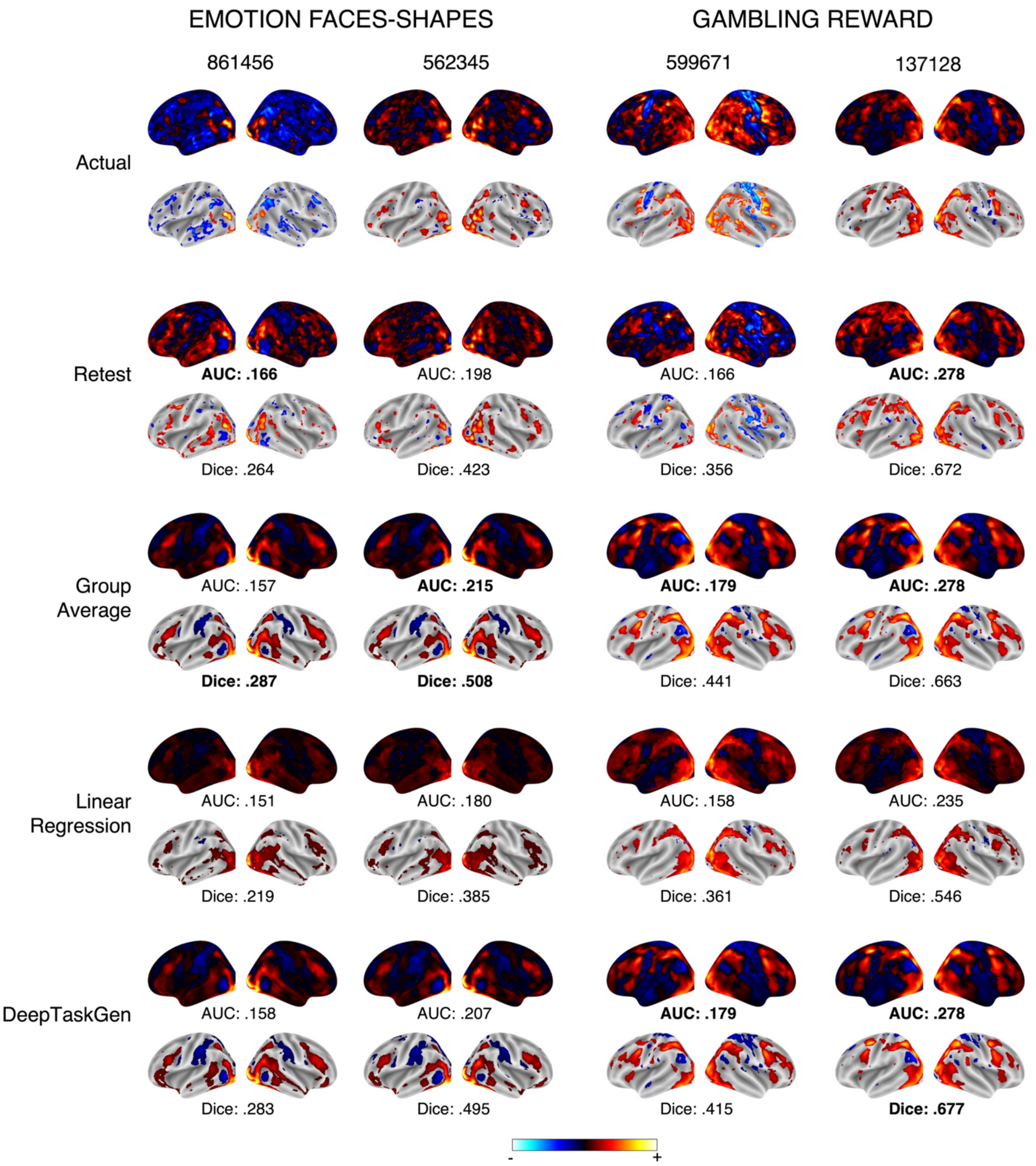


**Supplementary Figure 13.** Unthresholded and thresholded task activations for EMOTION FACES-SHAPES and GAMBLING REWARD contrasts are presented for sample atypical and typical subjects. In each task contrast map, the left column represents a typical subject, while the right column represents an atypical subject, defined by their similarity to the corresponding group average task activations. For each method, unthresholded task activations are displayed at the top, and thresholded activations (top 25% most activated voxels) are shown at the bottom. Dice AUC scores for unthresholded maps and Dice scores for thresholded maps are provided below the corresponding images.

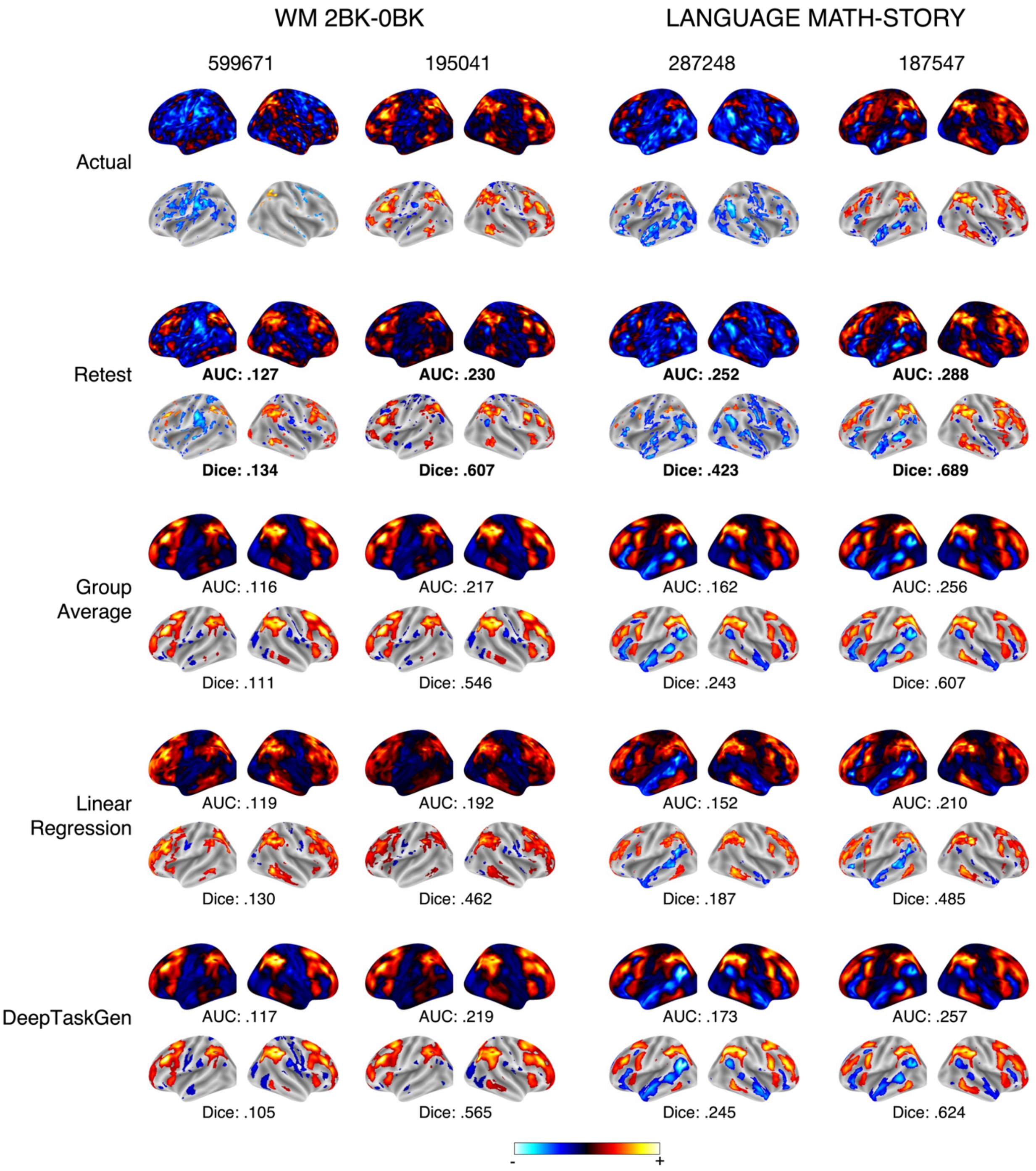


**Supplementary Figure 14.** Unthresholded and thresholded task activations for WM 2BK-0BK and LANGUAGE MATH-STORY contrasts are presented for sample atypical and typical subjects. In each task contrast map, the left column represents a typical subject, while the right column represents an atypical subject, defined by their similarity to the corresponding group average task activations. For each method, unthresholded task activations are displayed at the top, and thresholded activations (top 25% most activated voxels) are shown at the bottom. Dice AUC scores for unthresholded maps and Dice scores for thresholded maps are provided below the corresponding images.

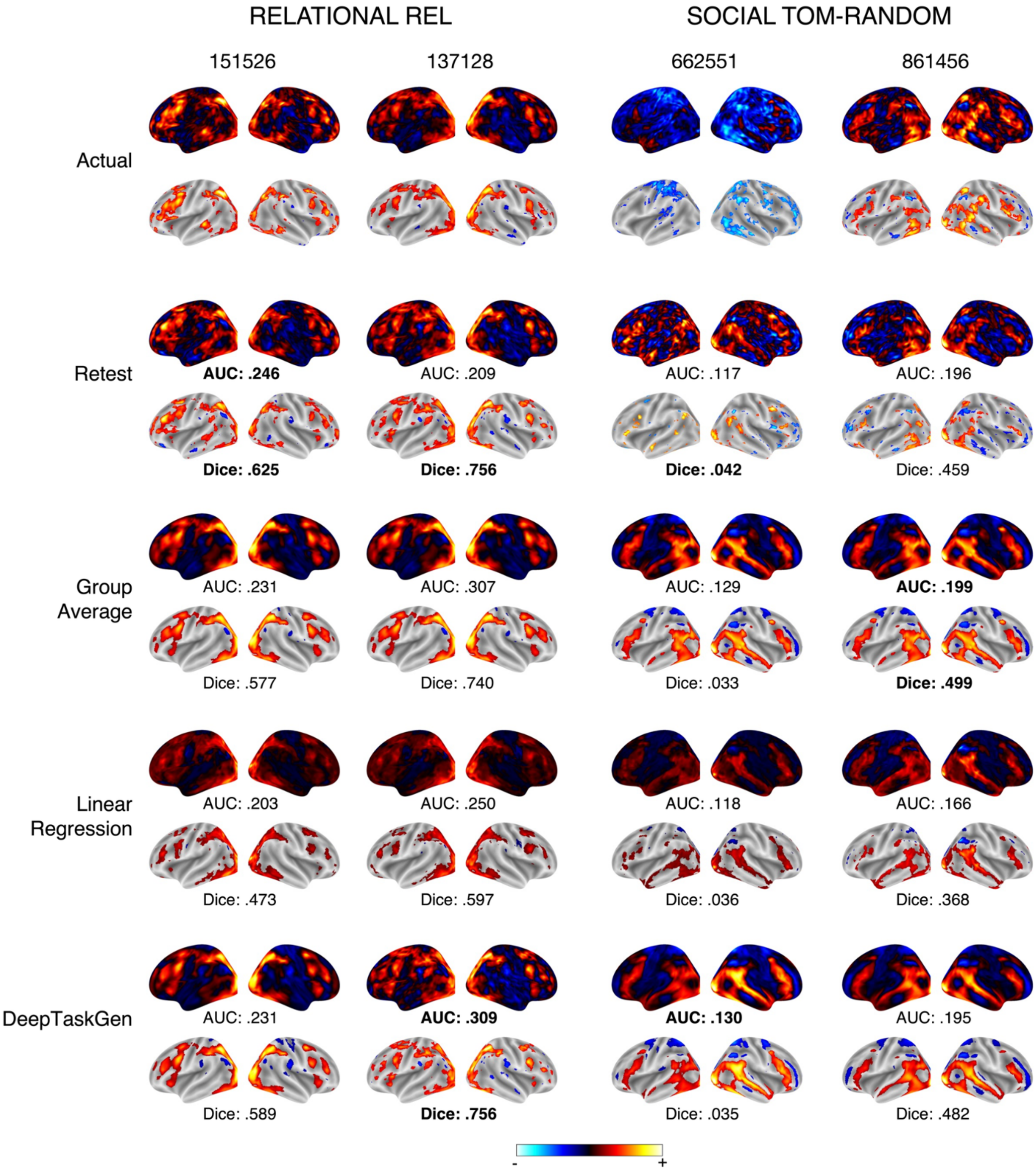


**Supplementary Figure 15.** Unthresholded and thresholded task activations for RELATIONAL REL and SOCIAL TOM-RANDOM contrasts are presented for sample atypical and typical subjects. In each task contrast map, the left column represents a typical subject, while the right column represents an atypical subject, defined by their similarity to the corresponding group average task activations. For each method, unthresholded task activations are displayed at the top, and thresholded activations (top 25% most activated voxels) are shown at the bottom. Dice AUC scores for unthresholded maps and Dice scores for thresholded maps are provided below the corresponding images.

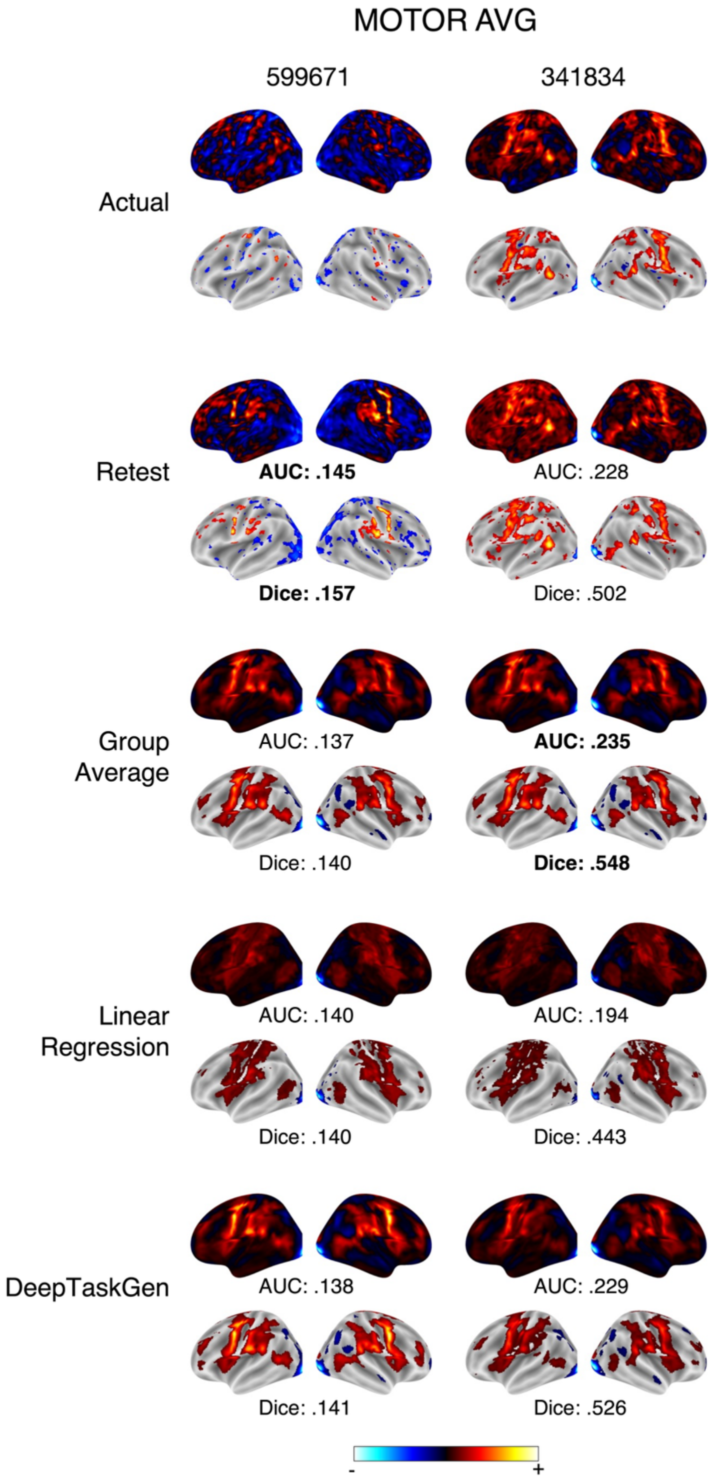

**Supplementary Figure 16.** Unthresholded and thresholded task activations for MOTOR AVG contrast are presented for sample atypical and typical subjects. In each task contrast map, the left column represents a typical subject, while the right column represents an atypical subject, defined by their similarity to the corresponding group average task activations. For each method, unthresholded task activations are displayed at the top, and thresholded activations (top 25% most activated voxels) are shown at the bottom. Dice AUC scores for unthresholded maps and Dice scores for thresholded maps are provided below the corresponding images.

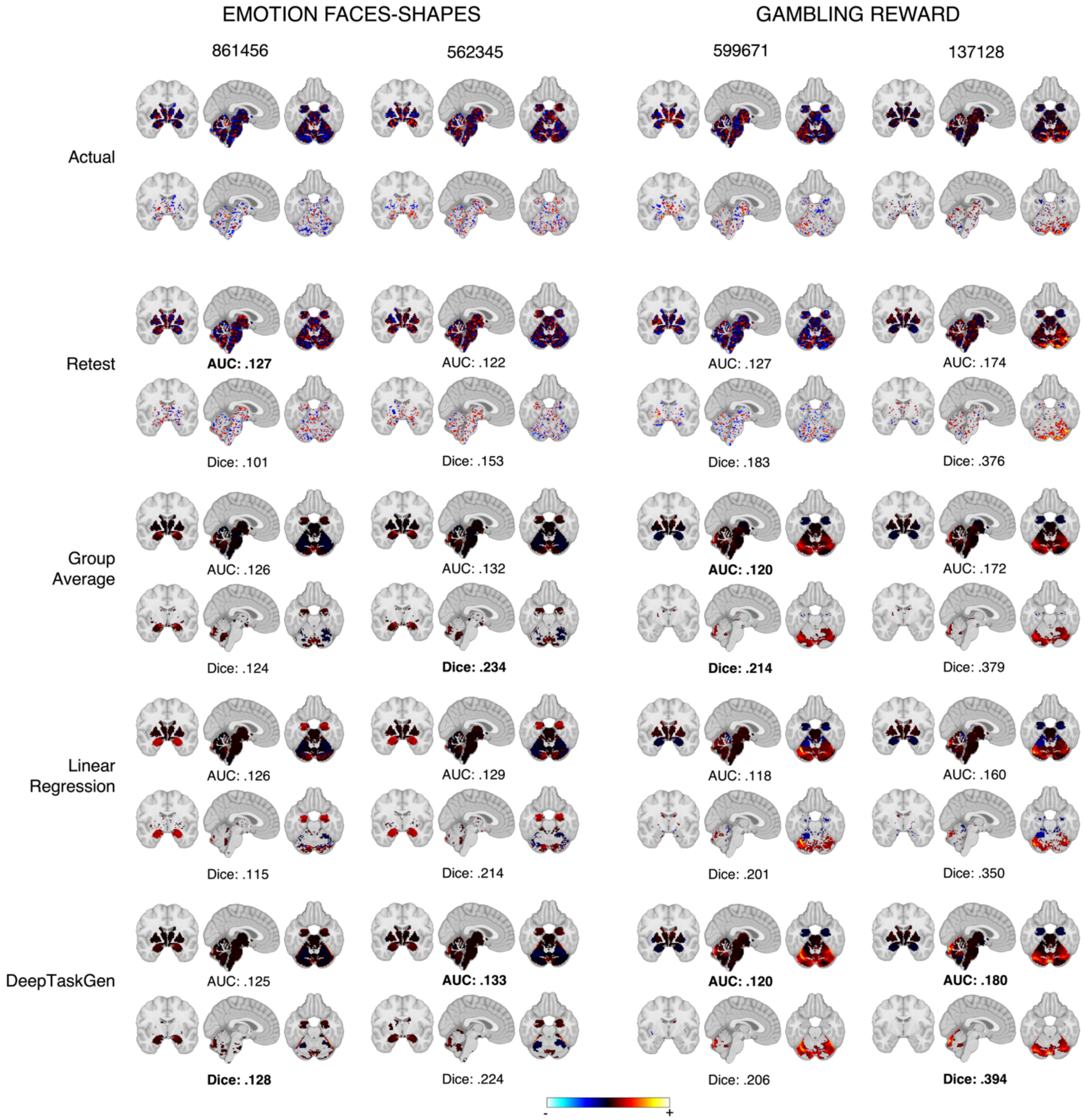


**Supplementary Figure 17.** Unthresholded and thresholded subcortical task activations for EMOTION FACES-SHAPES and GAMBLING REWARD contrasts are presented for sample atypical and typical subjects. In each task contrast map, the left column represents a typical subject, while the right column represents an atypical subject, defined by their similarity to the corresponding group average task activations. For each method, unthresholded task activations are displayed at the top, and thresholded activations (top 25% most activated voxels) are shown at the bottom. Dice AUC scores for unthresholded maps and Dice scores for thresholded maps are provided below the corresponding images.

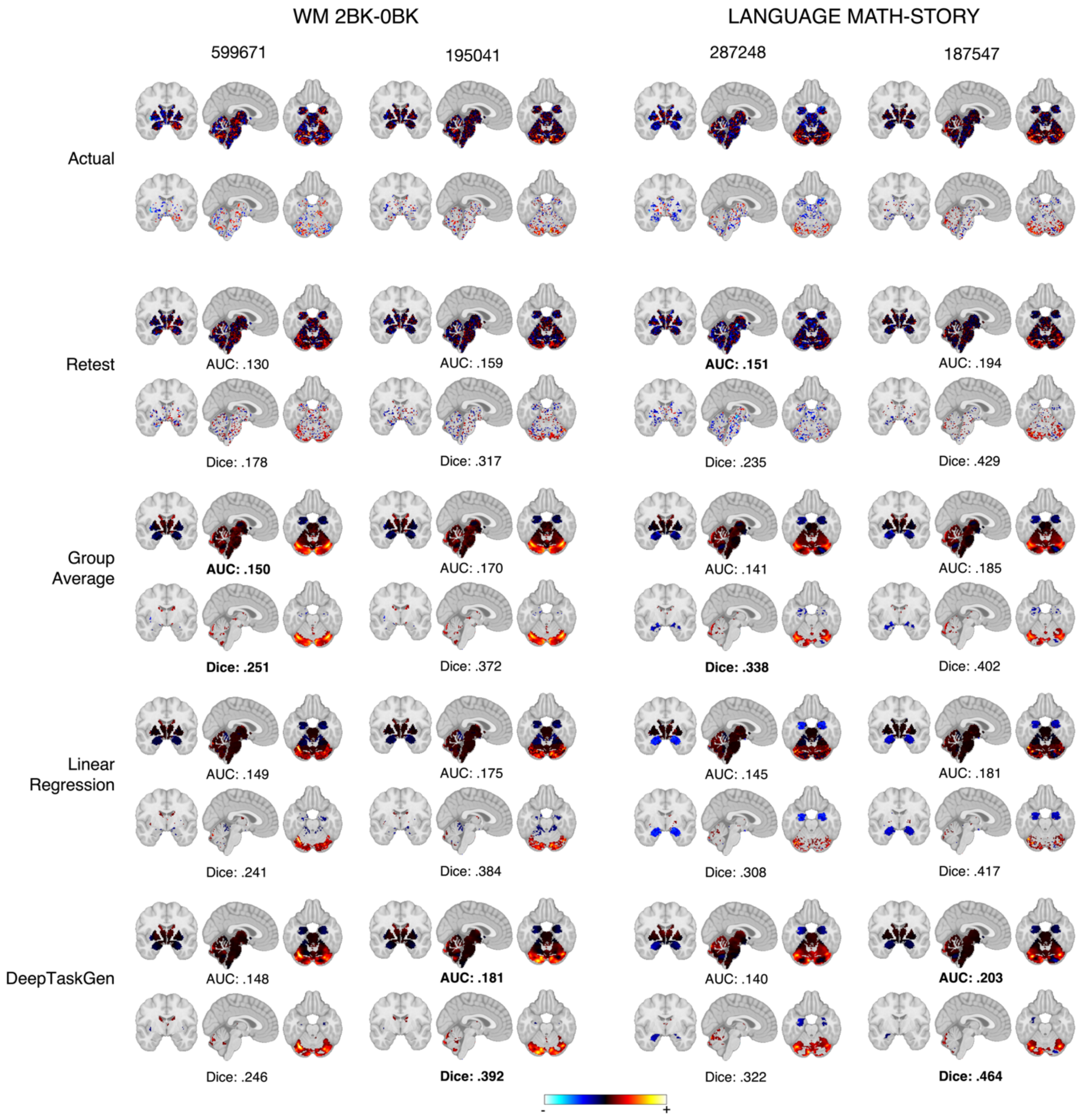


**Supplementary Figure 18.** Unthresholded and thresholded subcortical task activations for WM 2BK-0BK and LANGUAGE MATH-STORY contrasts are presented for sample atypical and typical subjects. In each task contrast map, the left column represents a typical subject, while the right column represents an atypical subject, defined by their similarity to the corresponding group average task activations. For each method, unthresholded task activations are displayed at the top, and thresholded activations (top 25% most activated voxels) are shown at the bottom. Dice AUC scores for unthresholded maps and Dice scores for thresholded maps are provided below the corresponding images.

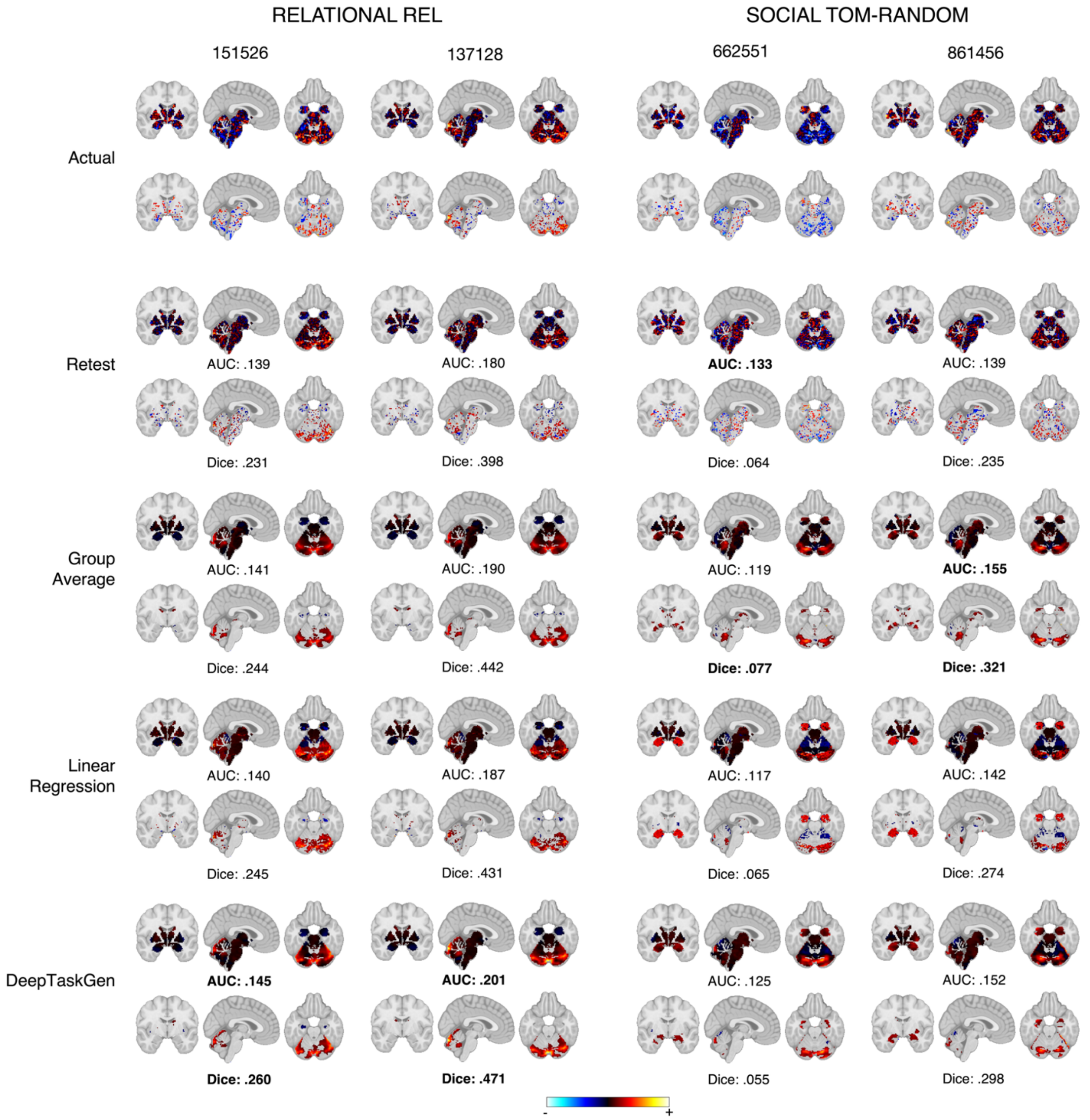


**Supplementary Figure 19.** Unthresholded and thresholded subcortical task activations for RELATIONAL REL and SOCIAL TOM-RANDOM contrasts are presented for sample atypical and typical subjects. In each task contrast map, the left column represents a typical subject, while the right column represents an atypical subject, defined by their similarity to the corresponding group average task activations. For each method, unthresholded task activations are displayed at the top, and thresholded activations (top 25% most activated voxels) are shown at the bottom. Dice AUC scores for unthresholded maps and Dice scores for thresholded maps are provided below the corresponding images.

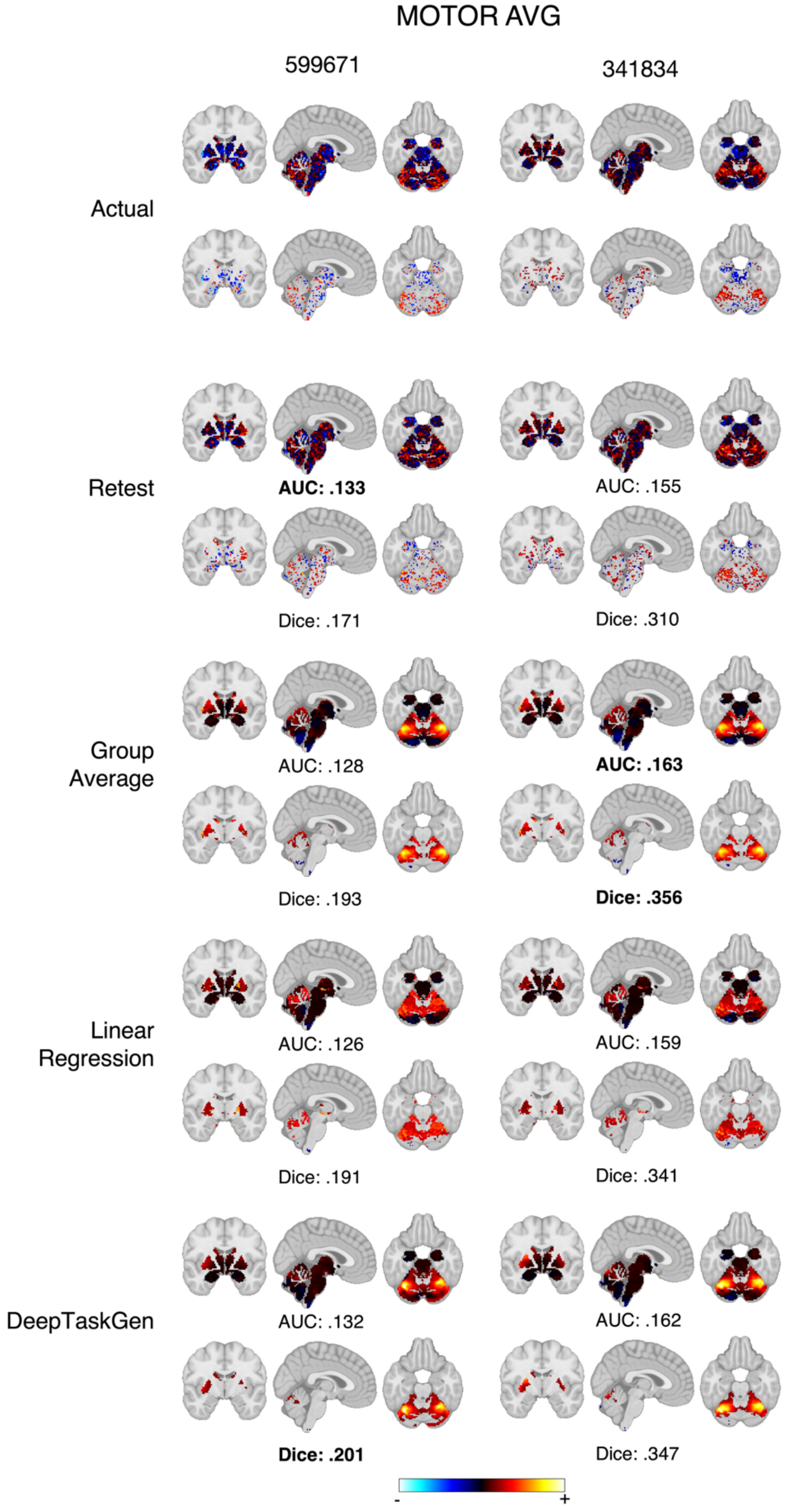

**Supplementary Figure 20.** Unthresholded and thresholded subcortical task activations for MOTOR AVG contrast are presented for sample atypical and typical subjects. In each task contrast map, the left column represents a typical subject, while the right column represents an atypical subject, defined by their similarity to the corresponding group average task activations. For each method, unthresholded task activations are displayed at the top, and thresholded activations (top 25% most activated voxels) are shown at the bottom. Dice AUC scores for unthresholded maps and Dice scores for thresholded maps are provided below the corresponding images.

| Task Contrast | DeepTaskGen vs. Group Average | | | DeepTaskGen vs. Retest Scans | | | DeepTaskGen vs. Linear Model | | |
|---|---|---|---|---|---|---|---|---|---|
| | *t* | *p* | *δ* | *t* | *p* | *δ* | *t* | *p* | *δ* |
| LANGUAGE MATH | -4.996 | -0.119 | 0.003 | 4.227 | 0.339 | 0.003 | 27.981 | 0.85 | 0.003 |
| LANGUAGE STORY | -4.589 | -0.099 | 0.003 | 4.607 | 0.335 | 0.003 | 29.489 | 0.553 | 0.003 |
| LANGUAGE MATH-STORY | -1.367 | -0.012 | 0.179 | -6.111 | -0.395 | 0.003 | 45.423 | 0.859 | 0.003 |
| RELATIONAL MATCH | -2.058 | -0.024 | 0.072 | -2.125 | -0.126 | 0.05 | 50.555 | 0.955 | 0.003 |
| RELATIONAL REL | -1.744 | -0.023 | 0.09 | -3.144 | -0.195 | 0.013 | 54.825 | 0.958 | 0.003 |
| RELATIONAL MATCH-REL | -11.443 | -0.111 | 0.003 | 3.238 | 0.285 | 0.005 | 11.626 | 0.302 | 0.003 |
| SOCIAL RANDOM | -7.177 | -0.111 | 0.003 | -5.168 | -0.324 | 0.003 | 41.246 | 0.941 | 0.003 |
| SOCIAL TOM | -5.418 | -0.101 | 0.003 | -8.297 | -0.499 | 0.003 | 48.664 | 0.989 | 0.003 |
| SOCIAL TOM-RANDOM | -9.617 | -0.149 | 0.003 | 4.289 | 0.302 | 0.003 | 24.233 | 0.788 | 0.003 |
| EMOTION FACES | -8.689 | -0.128 | 0.003 | 6.237 | 0.332 | 0.003 | 28.483 | 0.841 | 0.003 |
| EMOTION SHAPES | -7.318 | -0.108 | 0.003 | 6.256 | 0.298 | 0.003 | 26.222 | 0.726 | 0.003 |
| EMOTION FACES-SHAPES | -9.908 | -0.147 | 0.003 | 1.009 | 0.122 | 0.297 | 31.08 | 0.905 | 0.003 |
| WM 2BK_BODY | -1.987 | -0.045 | 0.062 | 3.174 | 0.204 | 0.003 | 27.361 | 0.836 | 0.003 |
| WM 2BK_FACE | -0.15 | 0.005 | 0.919 | 1.331 | 0.06 | 0.235 | 32.019 | 0.913 | 0.003 |
| WM 2BK_PLACE | -1.058 | -0.01 | 0.341 | 4.454 | 0.224 | 0.003 | 38.255 | 0.904 | 0.003 |
| WM 2BK_TOOL | -1.413 | -0.015 | 0.192 | 1.917 | 0.124 | 0.09 | 25.06 | 0.761 | 0.003 |
| WM 0BK_BODY | -2.186 | -0.014 | 0.046 | 5.316 | 0.329 | 0.003 | 23.219 | 0.758 | 0.003 |

| | | | | | | | | | |
|---|---|---|---|---|---|---|---|---|---|
| WM 0BK_FACE | -2.568 | -0.039 | 0.023 | 7.866 | 0.461 | 0.003 | 27.307 | 0.849 | 0.003 |
| WM 0BK_PLACE | -0.972 | -0.024 | 0.35 | 5.448 | 0.378 | 0.003 | 46.263 | 0.911 | 0.003 |
| WM 0BK_TOOL | -1.339 | -0.01 | 0.199 | 7.088 | 0.486 | 0.003 | 32.31 | 0.888 | 0.003 |
| WM 2BK | -0.507 | -0.014 | 0.643 | -2.218 | -0.302 | 0.052 | 35.731 | 0.892 | 0.003 |
| WM 0BK | -0.82 | -0.015 | 0.432 | 1.267 | 0.051 | 0.235 | 37.632 | 0.888 | 0.003 |
| WM 2BK-0BK | -0.456 | -0.011 | 0.663 | 1.745 | 0.061 | 0.108 | 14.1 | 0.448 | 0.003 |
| WM BODY | -2.044 | -0.034 | 0.048 | 1.138 | 0.053 | 0.342 | 26.863 | 0.804 | 0.003 |
| WM FACE | -1.006 | -0.01 | 0.344 | 0.564 | -0.069 | 0.613 | 32.556 | 0.9 | 0.003 |
| WM PLACE | -0.904 | -0.014 | 0.39 | 2.166 | 0.093 | 0.07 | 52.646 | 0.993 | 0.003 |
| WM TOOL | -0.985 | -0.018 | 0.337 | 1.432 | 0.089 | 0.223 | 32.486 | 0.854 | 0.003 |
| WM BODY-AVG | -6.535 | -0.098 | 0.003 | 11.004 | 0.621 | 0.003 | 19.991 | 0.775 | 0.003 |
| WM FACE-AVG | -8.533 | -0.083 | 0.003 | 6.66 | 0.377 | 0.003 | 21.899 | 0.717 | 0.003 |
| WM PLACE-AVG | -7.555 | -0.066 | 0.003 | 8.988 | 0.528 | 0.003 | 24.026 | 0.779 | 0.003 |
| WM TOOL-AVG | -7.475 | -0.089 | 0.003 | 7.225 | 0.584 | 0.003 | 18.31 | 0.477 | 0.003 |
| MOTOR CUE | -6.986 | -0.091 | 0.003 | 3.557 | 0.273 | 0.003 | 35.202 | 0.819 | 0.003 |
| MOTOR LF | -5.479 | -0.068 | 0.003 | 4.813 | 0.377 | 0.003 | 26.598 | 0.767 | 0.003 |
| MOTOR RF | -7.021 | -0.111 | 0.003 | 5.078 | 0.379 | 0.003 | 22.516 | 0.717 | 0.003 |
| MOTOR LH | -6.661 | -0.077 | 0.003 | 4.974 | 0.302 | 0.003 | 24.779 | 0.702 | 0.003 |
| MOTOR RH | -6.013 | -0.086 | 0.003 | 5.429 | 0.433 | 0.003 | 22.357 | 0.686 | 0.003 |
| MOTOR T | -6.994 | -0.094 | 0.003 | 8.026 | 0.479 | 0.003 | 24.727 | 0.715 | 0.003 |
| MOTOR AVG | -4.245 | -0.07 | 0.003 | 2.982 | 0.227 | 0.008 | 27.722 | 0.788 | 0.003 |
| MOTOR CUE-AVG | -7.689 | -0.08 | 0.003 | 0.924 | 0.108 | 0.395 | 41.548 | 0.807 | 0.003 |
| MOTOR LF-AVG | -11.464 | -0.139 | 0.003 | 10.545 | 0.763 | 0.003 | 19.938 | 0.763 | 0.003 |
| MOTOR LH-AVG | -12.7 | -0.156 | 0.003 | 10.4 | 0.669 | 0.003 | 20.203 | 0.545 | 0.003 |
| MOTOR RF-AVG | -12.943 | -0.143 | 0.003 | 13.923 | 0.749 | 0.003 | 18.696 | 0.732 | 0.003 |
| MOTOR RH-AVG | -12.459 | -0.145 | 0.003 | 11.155 | 0.678 | 0.003 | 20.569 | 0.511 | 0.003 |
| MOTOR T-AVG | -14.479 | -0.108 | 0.003 | 8.646 | 0.563 | 0.003 | 22.381 | 0.55 | 0.003 |
| GAMBLING PUNISH | -2.401 | -0.047 | 0.016 | 0.756 | 0.016 | 0.508 | 43.316 | 0.909 | 0.003 |
| GAMBLING REWARD | -1.888 | -0.039 | 0.064 | 0.44 | 0.005 | 0.704 | 45.671 | 0.954 | 0.003 |

| | | | | | | | | | |
|---|---|---|---|---|---|---|---|---|---|
| GAMBLING PUNISH-REWARD | -6.999 | -0.265 | 0.003 | 4.227 | 0.339 | 0.003 | 27.981 | 0.85 | 0.003 |

**Supplementary Table 3:** Paired t-tests with permutation testing ($P = 1000$) were performed to compare DeepTaskGen's reconstruction performance with various baseline methods, including group-average task contrast maps, retest scans, and the linear model on HCP-YA. The sample size for this comparison was 39. *p* values are FDR corrected for 3 pairwise comparisons across 47 task contrasts. Cliff's Delta ($\delta$) was used to measure effect size.

| Task Contrast | DeepTaskGen vs. Group Average | | | DeepTaskGen vs. Retest Scans | | | DeepTaskGen vs. Linear Model | | |
|---|---|---|---|---|---|---|---|---|---|
| | *t* | *p* | *δ* | *t* | *p* | *δ* | *t* | *p* | *δ* |
| LANGUAGE MATH | 11.608 | 0.002 | 0.949 | -9.596 | 0.002 | -0.97 | 8.008 | 0.002 | 0.704 |
| LANGUAGE STORY | 10.268 | 0.002 | 0.846 | -7.628 | 0.002 | -0.886 | 7.799 | 0.002 | 0.661 |
| LANGUAGE MATH-STORY | 20.287 | 0.002 | 1 | -16.298 | 0.002 | -1 | 11.147 | 0.002 | 0.72 |
| RELATIONAL MATCH | 17.639 | 0.002 | 1 | -18.616 | 0.002 | -0.997 | 8.974 | 0.002 | 0.805 |
| RELATIONAL REL | 19.094 | 0.002 | 1 | -20.284 | 0.002 | -1 | 9.496 | 0.002 | 0.808 |
| RELATIONAL MATCH-REL | 4.593 | 0.002 | 0.487 | -5.755 | 0.002 | -0.78 | 2.479 | 0.032 | 0.204 |
| SOCIAL RANDOM | 10.275 | 0.002 | 0.897 | -22.657 | 0.002 | -1 | 4.279 | 0.002 | 0.508 |
| SOCIAL TOM | 12.872 | 0.002 | 0.949 | -28.626 | 0.002 | -1 | 5.368 | 0.002 | 0.577 |
| SOCIAL TOM-RANDOM | 7.303 | 0.002 | 0.846 | -7.39 | 0.002 | -0.832 | 2.817 | 0.015 | 0.258 |
| EMOTION FACES | 10.316 | 0.002 | 0.949 | -13.586 | 0.002 | -0.938 | 4.741 | 0.002 | 0.49 |
| EMOTION SHAPES | 7.361 | 0.002 | 0.744 | -11.191 | 0.002 | -0.844 | 3.356 | 0.004 | 0.37 |
| EMOTION FACES-SHAPES | 9.82 | 0.002 | 0.897 | -10.697 | 0.002 | -0.997 | 4.426 | 0.002 | 0.482 |
| WM 2BK_BODY | 13.248 | 0.002 | 0.949 | -10.754 | 0.002 | -0.913 | 8.622 | 0.002 | 0.744 |
| WM 2BK_FACE | 15.787 | 0.002 | 1 | -11.151 | 0.002 | -0.838 | 10.323 | 0.002 | 0.808 |
| WM 2BK_PLACE | 14.629 | 0.002 | 0.949 | -11.014 | 0.002 | -0.95 | 8.446 | 0.002 | 0.771 |
| WM 2BK_TOOL | 13.03 | 0.002 | 0.949 | -10.594 | 0.002 | -0.871 | 8.22 | 0.002 | 0.695 |
| WM 0BK_BODY | 10.845 | 0.002 | 0.846 | -9.003 | 0.002 | -0.807 | 6.228 | 0.002 | 0.637 |
| WM 0BK_FACE | 13.443 | 0.002 | 0.949 | -9.776 | 0.002 | -0.854 | 6.838 | 0.002 | 0.708 |
| WM 0BK_PLACE | 14.396 | 0.002 | 0.949 | -10.52 | 0.002 | -0.811 | 7.519 | 0.002 | 0.663 |
| WM 0BK_TOOL | 12.971 | 0.002 | 0.949 | -9.386 | 0.002 | -0.851 | 6.43 | 0.002 | 0.641 |
| WM 2BK | 16.803 | 0.002 | 1 | -14.447 | 0.002 | -0.949 | 9.949 | 0.002 | 0.817 |
| WM 0BK | 14.339 | 0.002 | 0.949 | -13.561 | 0.002 | -0.913 | 7.533 | 0.002 | 0.725 |
| WM 2BK-0BK | 8.813 | 0.002 | 0.897 | -6.878 | 0.002 | -0.608 | 5.558 | 0.002 | 0.394 |

| | | | | | | | | | |
|---|---|---|---|---|---|---|---|---|---|
| WM BODY | 13.467 | 0.002 | 0.949 | -12.345 | 0.002 | -0.876 | 8.052 | 0.002 | 0.732 |
| WM FACE | 16.113 | 0.002 | 0.949 | -12.982 | 0.002 | -0.909 | 9.278 | 0.002 | 0.809 |
| WM PLACE | 15.775 | 0.002 | 1 | -13.284 | 0.002 | -0.943 | 8.567 | 0.002 | 0.748 |
| WM TOOL | 14.828 | 0.002 | 1 | -12.586 | 0.002 | -0.941 | 7.92 | 0.002 | 0.725 |
| WM BODY-AVG | 5.448 | 0.002 | 0.59 | -7.098 | 0.002 | -0.786 | 2.714 | 0.015 | 0.293 |
| WM FACE-AVG | 7.433 | 0.002 | 0.641 | -7.921 | 0.002 | -0.855 | 3.324 | 0.004 | 0.408 |
| WM PLACE-AVG | 6.569 | 0.002 | 0.795 | -8.481 | 0.002 | -0.804 | 2.414 | 0.036 | 0.337 |
| WM TOOL-AVG | 5.09 | 0.002 | 0.641 | -6.142 | 0.002 | -0.663 | 2.075 | 0.058 | 0.248 |
| MOTOR CUE | 12.452 | 0.002 | 0.949 | -11.308 | 0.002 | -0.946 | 7.891 | 0.002 | 0.708 |
| MOTOR LF | 9.191 | 0.002 | 0.949 | -7.807 | 0.002 | -0.749 | 5.788 | 0.002 | 0.488 |
| MOTOR RF | 8.827 | 0.002 | 0.897 | -7.102 | 0.002 | -0.804 | 4.613 | 0.002 | 0.487 |
| MOTOR LH | 9.154 | 0.002 | 0.897 | -7.259 | 0.002 | -0.779 | 5.161 | 0.002 | 0.486 |
| MOTOR RH | 6.78 | 0.002 | 0.795 | -6.967 | 0.002 | -0.801 | 3.621 | 0.002 | 0.324 |
| MOTOR T | 7.652 | 0.002 | 0.846 | -8.14 | 0.002 | -0.942 | 3.082 | 0.002 | 0.282 |
| MOTOR AVG | 9.475 | 0.002 | 1 | -8.579 | 0.002 | -0.905 | 4.917 | 0.002 | 0.452 |
| MOTOR CUE-AVG | 12.945 | 0.002 | 1 | -16.503 | 0.002 | -1 | 7.409 | 0.002 | 0.645 |
| MOTOR LF-AVG | 4.165 | 0.002 | 0.385 | -5.623 | 0.002 | -0.74 | 1.106 | 0.243 | 0.106 |
| MOTOR LH-AVG | 3.062 | 0.007 | 0.385 | -6.383 | 0.002 | -0.695 | 0.88 | 0.363 | 0.09 |
| MOTOR RF-AVG | 5.46 | 0.002 | 0.59 | -4.714 | 0.002 | -0.54 | 1.896 | 0.068 | 0.25 |
| MOTOR RH-AVG | 2.514 | 0.019 | 0.282 | -7.143 | 0.002 | -0.694 | -0.594 | 0.557 | -0.061 |
| MOTOR T-AVG | 4.392 | 0.004 | 0.487 | -9.104 | 0.002 | -0.992 | -0.429 | 0.653 | -0.043 |
| GAMBLING PUNISH | 16.275 | 0.002 | 1 | -13.436 | 0.002 | -0.966 | 8.211 | 0.002 | 0.695 |
| GAMBLING REWARD | 18.129 | 0.002 | 1 | -15.527 | 0.002 | -0.996 | 7.722 | 0.002 | 0.713 |
| GAMBLING PUNISH-REWARD | 2.676 | 0.002 | 0.333 | -9.596 | 0.002 | -0.97 | 8.008 | 0.002 | 0.704 |

**Supplementary Table 4:** Paired t-tests with permutation testing ($P$ = *1000*) were performed to compare DeepTaskGen's Diagonality Index scores with various baseline methods, including group-average task contrast maps, retest scans, and the linear model on HCP-YA. The sample size for this comparison was 39. $p$ values are FDR corrected for 3 pairwise comparisons across 47 task contrasts. Cliff's Delta ($\delta$) was used to measure effect size.

| Task Contrast | DeepTaskGen vs. Group Average | | | DeepTaskGen vs. Retest Scans | | | DeepTaskGen vs. Linear Model | | |
|---|---|---|---|---|---|---|---|---|---|
| | *t* | *p* | *δ* | *t* | *p* | *δ* | *t* | *p* | *δ* |
| LANGUAGE MATH | -2.636 | 0.012 | -0.06 | 2.628 | 0.023 | 0.223 | 17.284 | 0.003 | 0.553 |
| LANGUAGE STORY | -1.51 | 0.188 | -0.03 | 3.783 | 0.003 | 0.295 | 15.528 | 0.003 | 0.502 |
| LANGUAGE MATH-STORY | -0.472 | 0.679 | -0.005 | -6.854 | 0.003 | -0.41 | 25.105 | 0.003 | 0.761 |
| RELATIONAL MATCH | -1.13 | 0.318 | -0.015 | -0.073 | 0.901 | -0.031 | 32.03 | 0.003 | 0.804 |
| RELATIONAL REL | -1.17 | 0.318 | -0.015 | -1.347 | 0.206 | -0.09 | 29.418 | 0.003 | 0.761 |
| RELATIONAL MATCH-REL | -4.431 | 0.006 | -0.07 | 2.193 | 0.069 | 0.114 | 5.657 | 0.003 | 0.233 |
| SOCIAL RANDOM | -5.097 | 0.003 | -0.072 | -1.401 | 0.185 | -0.093 | 30.753 | 0.003 | 0.888 |
| SOCIAL TOM | -3.996 | 0.003 | -0.062 | -3.6 | 0.006 | -0.26 | 29.973 | 0.003 | 0.871 |
| SOCIAL TOM-RANDOM | -5.751 | 0.003 | -0.07 | 2.008 | 0.069 | 0.21 | 13.003 | 0.003 | 0.479 |
| EMOTION FACES | -3.666 | 0.006 | -0.041 | 6.123 | 0.003 | 0.327 | 23.305 | 0.003 | 0.628 |
| EMOTION SHAPES | -4.593 | 0.003 | -0.049 | 7.534 | 0.003 | 0.339 | 19.015 | 0.003 | 0.546 |
| EMOTION FACES-SHAPES | -7.912 | 0.003 | -0.157 | 0.254 | 0.853 | 0.116 | 20.776 | 0.003 | 0.663 |
| WM 2BK_BODY | -1.342 | 0.251 | -0.019 | 3.549 | 0.003 | 0.264 | 17.632 | 0.003 | 0.642 |
| WM 2BK_FACE | -0.287 | 0.822 | -0.009 | 2.268 | 0.03 | 0.147 | 23.297 | 0.003 | 0.819 |
| WM 2BK_PLACE | -0.369 | 0.772 | -0.01 | 3.724 | 0.003 | 0.274 | 22.059 | 0.003 | 0.745 |
| WM 2BK_TOOL | -2.047 | 0.087 | -0.016 | 1.817 | 0.104 | 0.114 | 16.789 | 0.003 | 0.549 |
| WM 0BK_BODY | -0.7 | 0.531 | -0.007 | 4.35 | 0.003 | 0.265 | 15.486 | 0.003 | 0.491 |
| WM 0BK_FACE | 0.038 | 0.967 | -0.015 | 6.005 | 0.003 | 0.485 | 17.403 | 0.003 | 0.711 |
| WM 0BK_PLACE | -2.235 | 0.043 | -0.043 | 4.034 | 0.003 | 0.31 | 24.358 | 0.003 | 0.813 |
| WM 0BK_TOOL | -0.688 | 0.5 | -0.002 | 6.069 | 0.003 | 0.408 | 18.755 | 0.003 | 0.612 |
| WM 2BK | -0.877 | 0.444 | -0.022 | -0.192 | 0.859 | -0.035 | 24.572 | 0.003 | 0.841 |
| WM 0BK | -1.123 | 0.309 | -0.027 | 2.535 | 0.021 | 0.165 | 23.702 | 0.003 | 0.82 |
| WM 2BK-0BK | -0.499 | 0.676 | -0.001 | 1.265 | 0.286 | 0.078 | 8.584 | 0.003 | 0.294 |

| | | | | | | | | | |
|---|---|---|---|---|---|---|---|---|---|
| WM BODY | -1.434 | 0.214 | -0.012 | 2.61 | 0.021 | 0.16 | 19.199 | 0.003 | 0.684 |
| WM FACE | 0.129 | 0.901 | -0.002 | 2.454 | 0.03 | 0.135 | 24.23 | 0.003 | 0.846 |
| WM PLACE | -0.728 | 0.503 | -0.037 | 2.681 | 0.023 | 0.191 | 27.206 | 0.003 | 0.854 |
| WM TOOL | -0.891 | 0.444 | -0.005 | 2.262 | 0.038 | 0.126 | 21.363 | 0.003 | 0.715 |
| WM BODY-AVG | -3.975 | 0.006 | -0.057 | 6.991 | 0.003 | 0.386 | 12.542 | 0.003 | 0.486 |
| WM FACE-AVG | -5.979 | 0.003 | -0.062 | 5.217 | 0.003 | 0.295 | 15.62 | 0.003 | 0.46 |
| WM PLACE-AVG | -3.093 | 0.009 | -0.043 | 5.854 | 0.003 | 0.454 | 16.196 | 0.003 | 0.554 |
| WM TOOL-AVG | -2.282 | 0.04 | -0.049 | 3.271 | 0.003 | 0.35 | 8.526 | 0.003 | 0.35 |
| MOTOR CUE | -5.969 | 0.003 | -0.111 | 4.52 | 0.003 | 0.319 | 27.407 | 0.003 | 0.744 |
| MOTOR LF | -3.765 | 0.006 | -0.059 | 5.372 | 0.003 | 0.402 | 14.637 | 0.003 | 0.645 |
| MOTOR RF | -1.999 | 0.05 | -0.055 | 2.315 | 0.047 | 0.249 | 15.864 | 0.003 | 0.603 |
| MOTOR LH | -3.698 | 0.006 | -0.055 | 4.322 | 0.003 | 0.349 | 17.082 | 0.003 | 0.591 |
| MOTOR RH | -1.87 | 0.086 | -0.028 | 7.107 | 0.003 | 0.482 | 14.194 | 0.003 | 0.515 |
| MOTOR T | -3.466 | 0.003 | -0.045 | 4.267 | 0.003 | 0.261 | 18.168 | 0.003 | 0.495 |
| MOTOR AVG | -1.414 | 0.203 | -0.02 | 3.403 | 0.006 | 0.275 | 17.813 | 0.003 | 0.734 |
| MOTOR CUE-AVG | -4.995 | 0.003 | -0.051 | 3.407 | 0.006 | 0.248 | 23.216 | 0.003 | 0.67 |
| MOTOR LF-AVG | -5.065 | 0.003 | -0.081 | 6.604 | 0.003 | 0.596 | 7.385 | 0.003 | 0.236 |
| MOTOR LH-AVG | -8.699 | 0.003 | -0.093 | 4.659 | 0.003 | 0.34 | 4.297 | 0.006 | 0.09 |
| MOTOR RF-AVG | -8.18 | 0.003 | -0.11 | 5.968 | 0.003 | 0.436 | 7.07 | 0.003 | 0.145 |
| MOTOR RH-AVG | -4.931 | 0.003 | -0.07 | 6.586 | 0.003 | 0.423 | 5.286 | 0.003 | 0.101 |
| MOTOR T-AVG | -6.943 | 0.003 | -0.157 | 2.397 | 0.023 | 0.249 | 11.082 | 0.003 | 0.566 |
| GAMBLING PUNISH | -2.211 | 0.033 | -0.034 | 1.78 | 0.118 | 0.086 | 28.519 | 0.003 | 0.862 |
| GAMBLING REWARD | -2.439 | 0.025 | -0.04 | 2.025 | 0.057 | 0.211 | 30.51 | 0.003 | 0.771 |
| GAMBLING PUNISH-REWARD | -3.412 | 0.003 | -0.072 | 2.628 | 0.023 | 0.223 | 17.284 | 0.003 | 0.553 |

**Supplementary Table 5:** Paired t-tests with permutation testing ($P$ = *1000*) were performed to compare DeepTaskGen's Dice AUC score with various baseline methods, including group-average task contrast maps, retest scans, and the linear model on HCP-YA. The sample size for this comparison was 39. *p* values are FDR corrected for 3 pairwise comparisons across 47 task contrasts. Cliff's Delta ($\delta$) was used to measure effect size.

| Task Contrast | Group Average | Retest Scans | Linear Model | DeepTaskGen |
|---|---|---|---|---|
| LANGUAGE MATH | 0 | **0.337** | 0.111 | 0.256 |
| LANGUAGE STORY | 0 | **0.281** | 0.119 | 0.243 |
| LANGUAGE MATH-STORY | 0 | **0.671** | 0.341 | 0.601 |
| RELATIONAL MATCH | 0 | **0.696** | 0.078 | 0.518 |
| RELATIONAL REL | 0 | **0.735** | 0.120 | 0.547 |
| RELATIONAL MATCH-REL | 0 | **0.114** | 0.032 | 0.051 |
| SOCIAL RANDOM | 0 | **0.711** | 0.089 | 0.310 |
| SOCIAL TOM | 0 | **0.751** | 0.131 | 0.411 |
| SOCIAL TOM-RANDOM | 0 | **0.260** | 0.080 | 0.135 |
| EMOTION FACES | 0 | **0.478** | 0.053 | 0.247 |
| EMOTION SHAPES | 0 | **0.401** | 0.051 | 0.196 |
| EMOTION FACES-SHAPES | 0 | **0.460** | 0.025 | 0.153 |
| WM 2BK_BODY | 0 | **0.489** | 0.064 | 0.416 |
| WM 2BK_FACE | 0 | **0.533** | 0.103 | 0.474 |
| WM 2BK_PLACE | 0 | **0.562** | 0.083 | 0.516 |
| WM 2BK_TOOL | 0 | **0.498** | 0.062 | 0.390 |
| WM 0BK_BODY | 0 | **0.357** | 0.077 | 0.269 |
| WM 0BK_FACE | 0 | **0.392** | 0.056 | 0.277 |
| WM 0BK_PLACE | 0 | **0.544** | 0.037 | 0.393 |
| WM 0BK_TOOL | 0 | **0.449** | 0.081 | 0.409 |
| WM 2BK | 0 | **0.685** | 0.098 | 0.555 |
| WM 0BK | 0 | **0.641** | 0.076 | 0.494 |
| WM 2BK-0BK | 0 | **0.279** | 0.149 | 0.227 |
| WM BODY | 0 | **0.565** | 0.100 | 0.436 |
| WM FACE | 0 | **0.591** | 0.097 | 0.484 |
| WM PLACE | 0 | **0.635** | 0.065 | 0.562 |
| WM TOOL | 0 | **0.597** | 0.096 | 0.498 |

| | | | | |
|---|---|---|---|---|
| WM BODY-AVG | 0 | **0.159** | 0.018 | 0.082 |
| WM FACE-AVG | 0 | **0.263** | 0.007 | 0.104 |
| WM PLACE-AVG | 0 | **0.248** | 0.021 | 0.155 |
| WM TOOL-AVG | 0 | **0.081** | 0.009 | 0.040 |
| MOTOR CUE | 0 | **0.481** | 0.099 | 0.382 |
| MOTOR LF | 0 | **0.272** | 0.078 | 0.175 |
| MOTOR RF | 0 | **0.220** | 0.041 | 0.104 |
| MOTOR LH | 0 | **0.258** | 0.071 | 0.175 |
| MOTOR RH | 0 | **0.254** | 0.048 | 0.079 |
| MOTOR T | 0 | **0.284** | 0.033 | 0.149 |
| MOTOR AVG | 0 | **0.347** | 0.060 | 0.165 |
| MOTOR CUE-AVG | 0 | **0.579** | 0.066 | 0.341 |
| MOTOR LF-AVG | 0 | **0.091** | 0.006 | 0.035 |
| MOTOR LH-AVG | 0 | **0.134** | 0.032 | 0.040 |
| MOTOR RF-AVG | 0 | **0.083** | 0.039 | 0.017 |
| MOTOR RH-AVG | 0 | **0.104** | 0.015 | 0.028 |
| MOTOR T-AVG | 0 | **0.219** | 0.023 | 0.020 |
| GAMBLING PUNISH | 0 | **0.565** | 0.117 | 0.458 |
| GAMBLING REWARD | 0 | **0.624** | 0.135 | 0.475 |
| GAMBLING PUNISH-REWARD | 0 | **0.017** | 0.011 | 0.011 |

**Supplementary Table 6:** The discriminability scores computed using the fingerprinting method outlined in the main manuscript. The table presents the results obtained from analyses performed on the HCP-YA dataset. The highest discriminability score for each task contrast is highlighted, indicating the method that achieved the best performance in differentiating between individuals based on their task contrast maps.

| Task Contrast | DeepTaskGen Fine-tuned vs. Linear Model | | | DeepTaskGen Fine-tuned vs. DeepTaskGen Non-Fine-tuned | | |
|---|---|---|---|---|---|---|
| | *t* | *p* | *δ* | *t* | *p* | *δ* |
| EMOTION FACES-SHAPES | .018 | .995 | - | **-11.059** | **.004** | **-.624** |
| GAMBLING REWARD | **10.225** | **.004** | **.340** | **3.675** | **.005** | **.142** |

**Supplementary Table 7:** Paired t-tests with permutation testing ($P = 1000$) were performed to compare the reconstruction performance of the fine-tuned DeepTaskGen model with the linear model and the non-fine-tuned DeepTaskGen model on HCP-D. FDR corrected (across model and contrasts) significant tests are highlighted. The sample size for this comparison was 64. Cliff's Delta ($\delta$) was used to measure effect size.

| Task Contrast | DeepTaskGen Fine-tuned vs. Linear Model | | | DeepTaskGen Fine-tuned vs. DeepTaskGen Non-Fine-tuned | | |
|---|---|---|---|---|---|---|
| | *t* | *p* | *δ* | *t* | *p* | *δ* |
| EMOTION FACES-SHAPES | **3.998** | **.003** | **.347** | **4.392** | **.003** | **.333** |
| GAMBLING REWARD | 1.362 | .200 | - | **5.743** | **.003** | **.356** |

**Supplementary Table 8:** Paired t-tests with permutation testing ($P = 1000$) were performed to compare the Diagonality Index scores of the fine-tuned DeepTaskGen model with the linear model and the non-fine-tuned DeepTaskGen model on HCP-D. FDR corrected (across model and contrasts) significant tests are highlighted. The sample size for this comparison was 64. Cliff's Delta ($\delta$) was used to measure effect size.

| Task Contrast | DeepTaskGen Fine-tuned vs. Linear Model | | | DeepTaskGen Fine-tuned vs. DeepTaskGen Non-Fine-tuned | | |
|---|---|---|---|---|---|---|
| | *t* | *p* | *δ* | *t* | *p* | *δ* |
| EMOTION FACES-SHAPES | **-2.738** | **.004** | **-.211** | **-8.955** | **.003** | **-.559** |
| GAMBLING REWARD | **7.231** | **.003** | **.208** | **5.876** | **.003** | **.145** |

**Supplementary Table 9:** Paired t-tests with permutation testing ($P = 1000$) were performed to compare the Dice AUC of the fine-tuned DeepTaskGen model with the linear model and the non-fine-tuned DeepTaskGen model on HCP-D. FDR corrected (across model and contrasts) significant tests are highlighted. The sample size for this comparison was 64. Cliff's Delta ($\delta$) was used to measure effect size.

| Task Contrast | DeepTaskGen Fine-tuned | DeepTaskGen Non-Fine-tuned | Linear Model |
|---|---|---|---|
| EMOTION FACES-SHAPES | .137 | **.284** | .274 |
| GAMBLING REWARD | .141 | .069 | **.222** |

**Supplementary Table 10:** The discriminability scores for various models computed using the fingerprinting method outlined in the main manuscript. The table presents the results obtained from analyses performed on the HCP-D dataset. The highest discriminability score for each task contrast is highlighted, indicating the method that achieved the best performance in differentiating between individuals based on their task contrast maps.

| Measure | UK Biobank variable IDs |
|---|---|
| Age | 21003 |
| Sex | 31 |
| Fluid Intelligence | 20016 |
| Dominant Hand Strength | 46, 47, 1707 |
| Overall Health | 2178 |
| Alcohol Use Frequency | 1558 |
| Weekly Beer Intake | 1588 |
| Depression | 20002-0.1286 |
| Hypertension | 20002-0.1065 |
| Neuroticism | 20127 |
| PHQ-9 | 20507, 20508, 20510, 20511, 20513, 20514, 20517, 20518, 20519 |
| GAD-7 | 20505, 20506, 20509, 20512, 20515, 20516, 20520 |
| RDS-4 | 2050, 2060, 2070, 2080 |

**Supplementary Table 11:** Subjects' measures available in the UK Biobank. Abbreviations: PHQ-9: Patient Health Questionnaire-9; GAD-7: General Anxiety Disorder-7; RDS-4: Recent Depressive Symptoms-4.

| Task Contrast | | Age | | Sex | | Fluid Intelligence | | Grip Strength | | Overall Health | | Alcohol Use Frequency | | Beer Use Frequency | |
|---|---|---|---|---|---|---|---|---|---|---|---|---|---|---|---|
| | | $\mu_{cv}$ | *p* | $\mu_{cv}$ | *p* | $\mu_{cv}$ | *p* | $\mu_{cv}$ | *p* | $\mu_{cv}$ | *p* | $\mu_{cv}$ | *p* | $\mu_{cv}$ | *p* |
| Actual | RESTING STATE | **.413** | **.001** | **.781** | **.001** | **.113** | **.002** | **.357** | **.001** | .006 | .418 | .041 | .072 | **.131** | **.001** |
| | EMOTION FACES-SHAPES | **.378** | **.001** | **.704** | **.001** | **.118** | **.002** | **.364** | **.001** | **.066** | **.013** | -.016 | .727 | .043 | .107 |
| Predicted | EMOTION FACES-SHAPES | **.459** | **.001** | **.837** | **.001** | **.094** | **.003** | **.349** | **.001** | .037 | .114 | **.072** | **.008** | **.212** | **.001** |
| | GAMBLING REWARD | **.452** | **.001** | **.828** | **.001** | **.054** | **.040** | **.401** | **.001** | .015 | .329 | .044 | .076 | **.168** | **.001** |
| | LANGUAGE MATH-STORY | **.446** | **.001** | **.817** | **.001** | **.094** | **.001** | **.418** | **.001** | .017 | .302 | **.112** | **.001** | **.181** | **.001** |
| | MOTOR AVG | **.538** | **.001** | **.836** | **.001** | **.076** | **.009** | **.356** | **.001** | .013 | .308 | **.073** | **.010** | **.182** | **.001** |
| | RELATIONAL REL | **.475** | **.001** | **.820** | **.001** | **.056** | **.037** | **.450** | **.001** | .028 | .188 | **.055** | **.033** | **.160** | **.001** |
| | SOCIAL TOM-RANDOM | **.476** | **.001** | **.827** | **.001** | **.111** | **.002** | **.349** | **.001** | .035 | .135 | **.078** | **.007** | **.190** | **.001** |
| | WM 2BK-0BK | **.487** | **.001** | **.827** | **.001** | **.067** | **.012** | **.429** | **.001** | -.014 | .676 | **.053** | **.025** | **.182** | **.001** |

**Supplementary Table 12. Prediction performances for demographic, cognitive and behavioral measures.** Out-of-sample performance was evaluated using a 5-fold cross-validation framework and permutation testing. $\boldsymbol{\mu_{cv}}$ represents the mean CV scores. Balanced accuracy was used for sex, while Pearson's correlation coefficient assessed the remaining variables. Significant predictions ($p < .05$) are highlighted. Comparisons between actual and synthetic data, which achieved significant predictions, were made using permutation testing ($p = 1000$), and the results are presented in Supplementary Tables 14-18.

| Task Contrast | | Depression | | Hypertension | | GAD-7 | | RDS-4 | | PHQ-9 | | Neuroticism | |
|---|---|---|---|---|---|---|---|---|---|---|---|---|---|
| | | $\mu_{cv}$ | $p$ | $\mu_{cv}$ | $p$ | $\mu_{cv}$ | $p$ | $\mu_{cv}$ | $p$ | $\mu_{cv}$ | $p$ | $\mu_{cv}$ | $p$ |
| Actual | RESTING STATE | .504 | .368 | .508 | .250 | **.083** | **.008** | .018 | .289 | .050 | .083 | .032 | .152 |
| | EMOTION FACES-SHAPES | .502 | .345 | **.526** | **.017** | .031 | .183 | .012 | .328 | .008 | .399 | .048 | .083 |
| Predicted | EMOTION FACES-SHAPES | .513 | .405 | .517 | .087 | .013 | .377 | .029 | .166 | **.062** | **.038** | .015 | .314 |
| | GAMBLING REWARD | .502 | .405 | **.553** | **.001** | .043 | .118 | .033 | .150 | .016 | .342 | .035 | .132 |
| | LANGUAGE MATH-STORY | .486 | .945 | **.539** | **.003** | -.002 | .515 | .018 | .262 | .016 | .311 | .019 | .282 |
| | MOTOR AVG | .502 | .409 | **.524** | **.030** | **.091** | **.005** | .016 | .188 | **.085** | **.008** | .026 | .224 |
| | RELATIONAL REL | .509 | .155 | **.535** | **.005** | .053 | .065 | **.052** | **.049** | .031 | .191 | .047 | .075 |
| | SOCIAL TOM-RANDOM | .510 | .104 | .512 | .148 | .043 | .105 | -.004 | .536 | .039 | .163 | .019 | .296 |
| | WM 2BK-0BK | .508 | .174 | **.539** | **.001** | .034 | .159 | .041 | .009 | **.074** | **.015** | -.018 | .711 |

**Supplementary Table 13. Prediction performances for physical and mental health measures.** Out-of-sample performance was evaluated using a 5-fold cross-validation framework and permutation testing. $\boldsymbol{\mu_{cv}}$ represents the mean CV scores. Balanced accuracy was used for depression and hypertension classification, while Pearson's correlation coefficient assessed the remaining variables. Significant predictions ($p < .05$) are highlighted. Comparisons between actual and synthetic data, which achieved significant predictions, were made using permutation testing ($P = 1000$), and the results are presented in Supplementary Tables 19,20.

| Task Contrast | | vs. Actual EMOTION FACES-SHAPES | | | vs. Resting State Connectome | | |
|---|---|---|---|---|---|---|---|
| | | *t* | *p* | *δ* | *t* | *p* | *δ* |
| Predicted | EMOTION FACES-SHAPES | 3.94 | .050 | - | 2.55 | .076 | - |
| | GAMBLING REWARD | 3.79 | .050 | - | 2.38 | .097 | - |
| | LANGUAGE MATH-STORY | **6.51** | **.030** | **1.0** | 2.92 | .066 | - |
| | MOTOR AVG | **6.59** | **.030** | **1.0** | **5.52** | **.038** | **1.0** |
| | RELATIONAL REL | **5.14** | **.038** | **1.0** | 3.97 | .504 | - |
| | SOCIAL TOM-RANDOM | 3.43 | .050 | - | 2.45 | .076 | - |
| | WM 2BK-0BK | **5.29** | **.038** | **1.0** | 4.14 | .050 | - |
| Actual | Resting State Connectome | **10.19** | **.030** | **.68** | | | |

**Supplementary Table 14: Age prediction.** The results of paired t-tests with permutation testing $(P = 1000)$, comparing the brain age prediction performance of predicted task contrast maps, actual EMOTION FACES-SHAPES task contrast maps, and resting-state connectome data from the UK Biobank dataset. Significant differences $(p < .05, FDR\ corrected)$ are highlighted in bold. Cliff's Delta ($\delta$) was used to measure effect size.

| Task Contrast | | vs. Actual EMOTION FACES-SHAPES | | | vs. Resting State Connectome | | |
|---|---|---|---|---|---|---|---|
| | | *t* | *p* | *δ* | *t* | *p* | *δ* |
| Predicted | EMOTION FACES-SHAPES | **9.08** | **.012** | **1.0** | **6.79** | **.009** | **1.0** |
| | GAMBLING REWARD | **6.08** | **.012** | **1.0** | **6.63** | **.012** | **.84** |
| | LANGUAGE MATH-STORY | **8.21** | **.007** | **1.0** | **3.67** | **.034** | **.84** |
| | MOTOR AVG | **9.77** | **.007** | **1.0** | **3.53** | **.030** | **.92** |
| | RELATIONAL REL | **6.55** | **.012** | **1.0** | **12.8** | **.007** | **.84** |
| | SOCIAL TOM-RANDOM | **7.14** | **.012** | **1.0** | **4.02** | **.029** | **.84** |
| | WM 2BK-0BK | **9.26** | **.007** | **1.0** | 2.65 | .055 | - |
| Actual | Resting State Connectome | **3.76** | **.030** | **1.0** | | | |

**Supplementary Table 15: Sex classification.** The results of paired t-tests with permutation testing ($P = 1000$), comparing the sex classification performance of predicted task contrast maps, actual EMOTION FACES-SHAPES task contrast maps, and resting-state connectome data from the UK Biobank dataset. Significant differences ($p < .05, FDR\ corrected$) are highlighted in bold. Cliff's Delta ($\delta$) was used to measure effect size.

| Task Contrast | | vs. Actual EMOTION FACES-SHAPES | | vs. Resting State Connectome | |
|---|---|---|---|---|---|
| | | *t* | *p* | *t* | *p* |
| Predicted | EMOTION FACES-SHAPES | -.78 | .433 | -.91 | .433 |
| | GAMBLING REWARD | -2.23 | .378 | -2.71 | .378 |
| | LANGUAGE MATH-STORY | -.63 | .433 | -.64 | .433 |
| | MOTOR AVG | -1.19 | .433 | -1.31 | .433 |
| | RELATIONAL REL | -2.54 | .378 | -2.22 | .378 |
| | SOCIAL TOM-RANDOM | -.47 | .458 | -.06 | .495 |
| | WM 2BK-0BK | -1.92 | .402 | -1.66 | .407 |
| Actual | Resting State Connectome | -.11 | .495 | | |

**Supplementary Table 16: Fluid intelligence prediction.** The results of paired t-tests with permutation testing ($P = 1000$), comparing the fluid intelligence prediction performance of predicted task contrast maps, actual EMOTION FACES-SHAPES task contrast maps, and resting-state connectome data from the UK Biobank dataset. No significant difference was found ($p < .05, FDR\ corrected$).

| Task Contrast | | vs. Actual EMOTION FACES-SHAPES | | vs. Resting State Connectome | |
|---|---|---|---|---|---|
| | | *t* | *p* | *t* | *p* |
| Predicted | EMOTION FACES-SHAPES | -.31 | .495 | -.58 | .495 |
| | GAMBLING REWARD | 1.09 | .466 | 2.20 | .466 |
| | LANGUAGE MATH-STORY | 1.64 | .466 | 1.65 | .466 |
| | MOTOR AVG | -.12 | .495 | -.02 | .495 |
| | RELATIONAL REL | 1.64 | .466 | 3.42 | .466 |
| | SOCIAL TOM-RANDOM | -.32 | .495 | -.22 | .495 |
| | WM 2BK-0BK | 1.13 | .466 | 2.40 | .466 |
| Actual | Resting State Connectome | -.13 | .495 | | |

**Supplementary Table 17: Dominant hand grip strength prediction.** The results of paired t-tests with permutation testing $(P = 1000)$, comparing the dominant hand grip strength prediction performance of predicted task contrast maps, actual EMOTION FACES-SHAPES task contrast maps, and resting-state connectome data from the UK Biobank dataset. No significant difference was found $(p < .05, FDR\ corrected)$.

| Task Contrast | | vs. Actual EMOTION FACES-SHAPES | | vs. Resting State Connectome | |
|---|---|---|---|---|---|
| | | *t* | *p* | *t* | *p* |
| Predicted | EMOTION FACES-SHAPES | - | - | 4.38 | .050 |
| | GAMBLING REWARD | - | - | 1.33 | .241 |
| | LANGUAGE MATH-STORY | - | - | 1.36 | .250 |
| | MOTOR AVG | - | - | 1.18 | .252 |
| | RELATIONAL REL | - | - | .94 | .252 |
| | SOCIAL TOM-RANDOM | - | - | 1.50 | .241 |
| | WM 2BK-0BK | - | - | 1.91 | .226 |
| Actual | Resting State Connectome | - | - | - | - |

**Supplementary Table 18: Weekly beer intake prediction.** The results of paired t-tests with permutation testing ($P = 1000$), comparing the weekly beer intake prediction performance of predicted task contrast maps, actual EMOTION FACES-SHAPES task contrast maps, and resting-state connectome data from the UK Biobank dataset. Actual EMOTION FACES-SHAPES did not survive the permutation test. No significant difference was found ($p < .05, FDR\ corrected$).

| Task Contrast | | vs. Actual EMOTION FACES-SHAPES | | vs. Resting State Connectome | |
|---|---|---|---|---|---|
| | | *t* | *p* | *t* | *p* |
| Predicted | EMOTION FACES-SHAPES | -.51 | .418 | - | - |
| | GAMBLING REWARD | 1.95 | .230 | - | - |
| | LANGUAGE MATH-STORY | 1.41 | .268 | - | - |
| | MOTOR AVG | -.21 | .452 | - | - |
| | RELATIONAL REL | .42 | .418 | - | - |
| | SOCIAL TOM-RANDOM | -3.39 | .206 | - | - |
| | WM 2BK-0BK | .96 | .348 | - | - |
| Actual | Resting State Connectome | - | - | - | - |

**Supplementary Table 19: Hypertension diagnosis classification.** The results of paired t-tests with permutation testing $(P = 1000)$, comparing the hypertension diagnosis classification performance of predicted task contrast maps, actual EMOTION FACES-SHAPES task contrast maps, and resting-state connectome data from the UK Biobank dataset. Actual resting-state connectome did not survive the permutation test. No significant difference was found $(p < .05, FDR\ corrected)$.

| Task Contrast | | vs. Actual EMOTION FACES-SHAPES | | vs. Resting State Connectome | |
|---|---|---|---|---|---|
| | | *t* | *p* | *t* | *p* |
| Predicted | EMOTION FACES-SHAPES | - | - | - | - |
| | GAMBLING REWARD | - | - | - | - |
| | LANGUAGE MATH-STORY | - | - | - | - |
| | MOTOR AVG | - | - | 2.08 | .357 |
| | RELATIONAL REL | - | - | - | - |
| | SOCIAL TOM-RANDOM | - | - | - | - |
| | WM 2BK-0BK | - | - | - | - |
| Actual | Resting State Connectome | - | - | | |

**Supplementary Table 20: GAD-7 prediction.** The results of paired t-tests with permutation testing $(P = 1000)$ comparing the subjects' overall health performance of predicted task contrast maps, actual EMOTION FACES-SHAPES task contrast maps, and resting-state connectome data from the UK Biobank dataset. Significant differences $(p < .05, FDR\ corrected)$ are highlighted in bold. Only synthetic MOTOR AVG and actual resting-state connectome significantly predicted GAD-7 scores.

### References


1. Abraham, A. *et al.* Machine learning for neuroimaging with scikit-learn. *Front. Neuroinform.* **8**, (2014).
2. Pruim, R. H. R. *et al.* ICA-AROMA: A robust ICA-based strategy for removing motion artifacts from fMRI data. *NeuroImage* **112**, 267–277 (2015).
3. Dice, L. R. Measures of the Amount of Ecologic Association Between Species. *Ecology* **26**, 297–302 (1945).
4. Ngo, G. H., Khosla, M., Jamison, K., Kuceyeski, A. & Sabuncu, M. R. Predicting individual task contrasts from resting-state functional connectivity using a surface-based convolutional network. *NeuroImage* **248**, 118849 (2022).
5. Finn, E. S. *et al.* Functional connectome fingerprinting: Identifying individuals using patterns of brain connectivity. *Nature Neuroscience* **18**, 1664–1671 (2015).
6. Tavor, I. *et al.* Task-free MRI predicts individual differences in brain activity during task performance. *Science* **352**, 216–220 (2016).
